# Disentangling Protein Function via Decoupled Information Theoretic Selection of Key Tuning Residues

**DOI:** 10.1101/2025.05.28.653817

**Authors:** Haris Saeed, Aidong Yang, Wei E. Huang

## Abstract

Rational protein engineering requires identifying residues that modulate function without disrupting functionality, a key challenge in protein engineering. Existing computational methods struggle to distinguish genuine functional sites from positions coevolving due to structural constraints, leading to high false-discovery rates. Here we present an information-theoretic decoupling framework that, without machine learning, isolates key tuning residues by computationally “denoising” sequence data, iteratively removing confounding evolutionary signals to reveal underlying functional sites. We validated this framework across 10 datasets spanning enzymes, fluorescent proteins, and antibodies. In a nanobody-antigen binding case study, our method identified *>* 25% (6/20) of verified binding-critical residues (*p* = 0.031), while the best of five benchmarked tools found zero. Performance was consistent across all datasets, with supervised variants achieving large effect sizes (Hedges’ *g >* 0.7, *p <* 0.01) and unsupervised variants also showing gains (*g >* 0.2, *p <* 0.05) over benchmarks. This interpretable framework provides a generalizable method to accelerate protein design, from focusing antibody maturation to optimizing biocatalysts.

## 1 Introduction

A central ambition in computational biology is to move beyond static protein analysis and toward dynamic design, navigating the vast sequence landscape to engineer functions and alter protein’s functional and environmental properties. We propose that this engineering challenge can be framed as a navigable Markov process, where mutations are guided steps toward an optimal functional state (Figure 1b). Building such a model, however, requires a high-fidelity map of the underlying sequence-function landscape. While coevolutionary methods like EVmutation have been successfully applied as epistasis models to guide protein design [1], a fundamental challenge remains: identifying the specific “levers”—the key tuning residues that govern this process.

**Figure 1:**
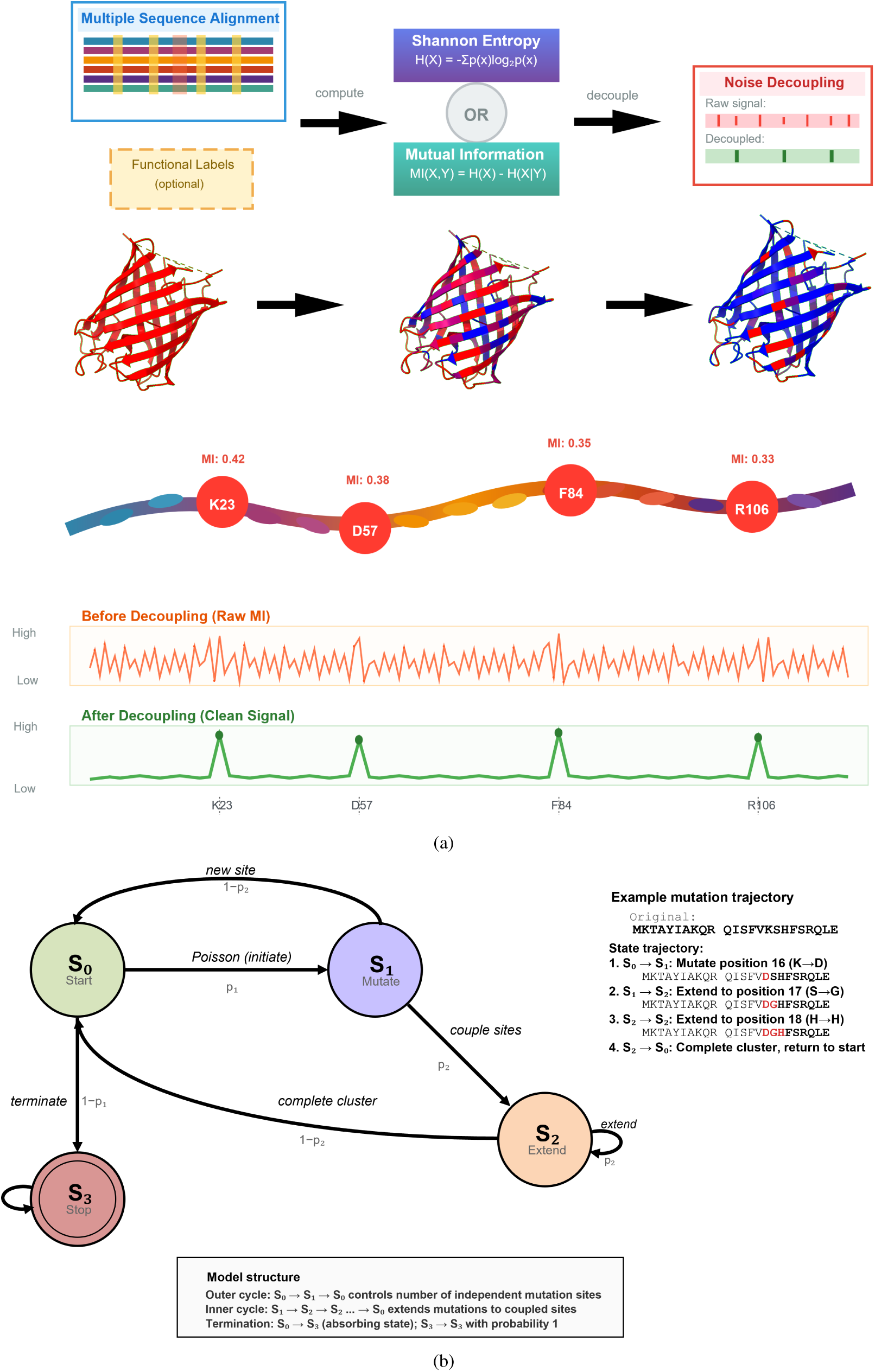
Framework for disentangling protein function. a Conceptual schematic of the analysis pipeline, illustrated with a synthetic GFP example: from the high-level workflow through to a flattened amino acid chain, and the corresponding residue trace before and after decoupling. b Application of the residue trace and interaction matrices to guide a genetic algorithm, where mutagenesis is represented as a Markov process with the traces and coupling matrices forming the basis of the state transition matrices

We define these key tuning residues as “rheostats”, positions where mutations can modulate a specific property, such as an enzyme’s *k*_cat_ or an antibody’s affinity, without catastrophically abolishing the protein’s core function [2–4]. These rheostats are the primary targets for engineering [5, 6], but finding them experimentally is a major bottleneck. A comprehensive high-throughput screen on even a small protein can require testing ∼4,000 variants and cost over $50,000, making a purely experimental approach prohibitive.

This cost has driven the development of computational tools to predict these sites. Existing methods span from structure-based (Hotspot Wizard [7]) and conservation-based (SIFT [8], Rate4Site [9]) to coevolutionary (EVmutation [10]) and deep learning models (DeepSequence [11]). These tools, however, often struggle to reliably discover key tuning residues. We propose this is due to an inability to distinguish functional tuning from structural maintenance. Residues often co-evolve simply to maintain the protein’s fold, and this strong structural signal can mask the more subtle signal of functional modulation. This defines a precise feature selection problem, where the goal is to find a subset of residues F ⊂ {1*, . . ., N* } that maximizes a functional objective *J*:

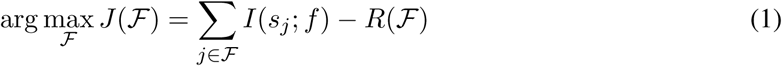

where *I*(*s_j_*; *f*) is the functional information at position *j* about property *f*, and *R*(F) is the redundant information from evolutionary coupling.

To explicitly solve this optimization problem, we propose a new, two-criterion hypothesis for what defines a key tuning residue. We posit a residue is a functional rheostat if and only if it is (1) **functionally correlated** with the target property (maximizing *I*(*s_j_*; *f*)), and (2) an **’Informational Hub’** that retains unique predictive information after confounding couplings are accounted for (minimizing *R*(F)). Here, we develop and test a flexible, information-theoretic framework (Figure 1a) built directly upon this hypothesis. We validated our framework across eight distinct protein families, benchmarking it against five widely used tools. Our results show that this decoupling-aware approach outperforms existing methods (*p <* 0.05 both supervised and unsupervised), providing mechanistically transparent parameters suitable for building the navigable Markov process we envision for protein design and optimization.

## 2 Results

### 2.1 A Four-Stage Pipeline to Test the Two-Criterion Hypothesis

To test our two-criterion hypothesis (Eq. 1), we developed a modular, four-stage pipeline (Fig. 1). This framework was designed to systematically test two major hypotheses: first, that a biochemically-aware encoding is superior to standard string representations, and second, that an explicit decoupling step is necessary to remove coupling noise.

#### 2.1.1 Stage 1: The Biochemical Encoding Hypothesis

The first stage of the pipeline addresses the encoding of the MSA. Standard, string-based (categorical) representations treat all substitutions as equally different, which is biochemically unrealistic. A conservative leucine ↔ isoleucine substitution is treated identically to a radical leucine ↔ aspartic acid one. We hypothesized that this representation would be a major source of noise.

This hypothesis was guided by preliminary analysis (Figure 2), which shows that standard Hamming distances blur protein family structure, while BLOSUM62-weighted distances reveal clear clustering (compare panels 2a and 2b). The BLOSUM62 histograms (panel 2d) show a clear separation between inter- and intra-family distances, which is reduced in the Hamming histograms (panel 2c).

**Figure 2:**
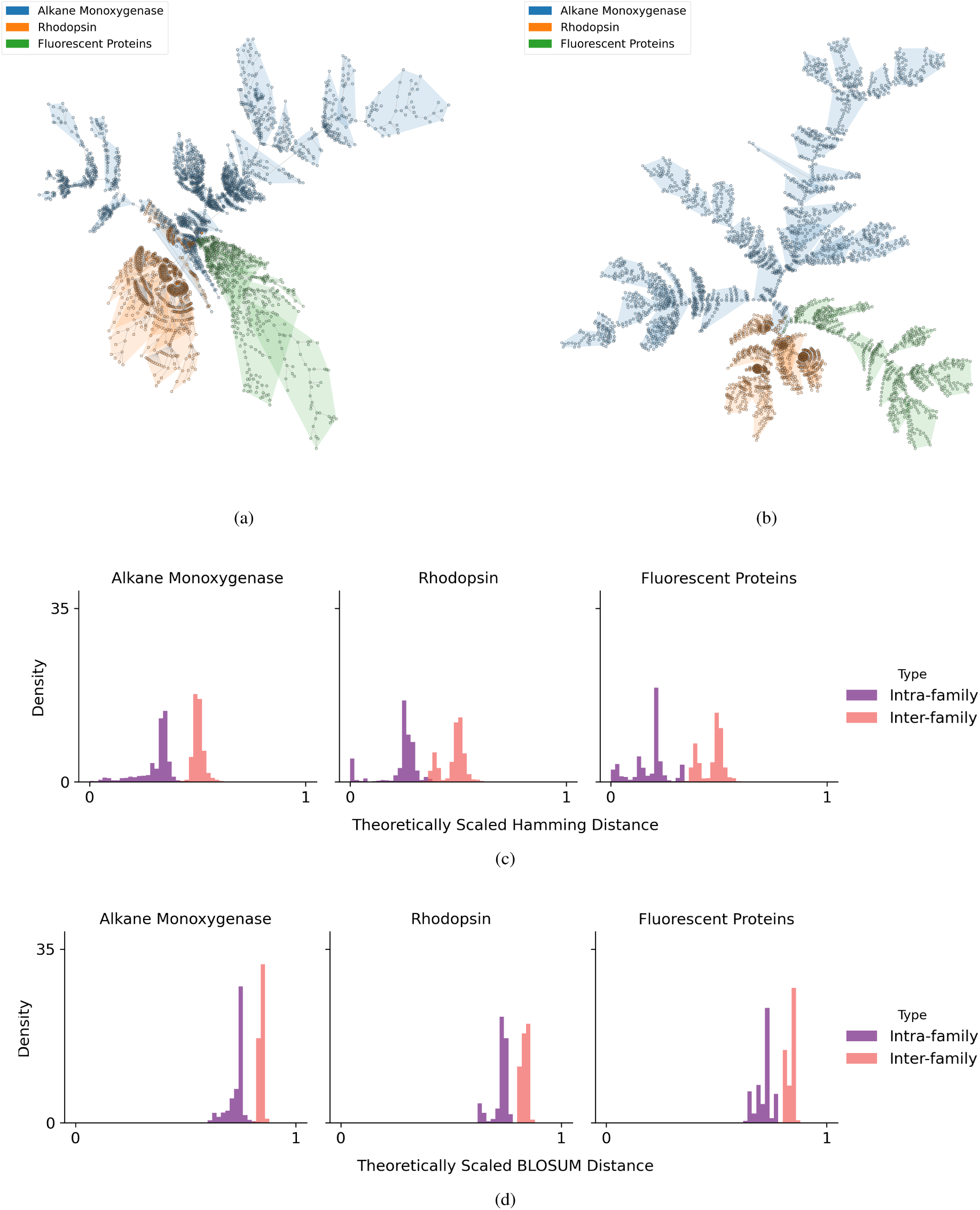
Biochemically-informed distance metrics reveal sharper protein family boundaries. Comparison of Hamming versus BLOSUM62-weighted distances across Alkane Monooxygenases (blue), Rhodopsins (orange), and Fluorescent Proteins (green). a Hamming distance minimum spanning tree shows extensive cross-family bridging. b BLOSUM62 tree produces clearer family-specific clusters. c Hamming distance distributions show broad overlap between within- and between-family distances. d BLOSUM62 distributions show sharper Gaussian separation. BLOSUM62 weighting by evolutionary exchangeability provides superior signal for coupling analysis, justifying physicochemical encoding.

To test this hypothesis, we compare two encoding strategies: (1) a standard categorical (string) encoding, and (2) a *cluster-then-separate* physicochemical (P-C) encoding that groups biochemically similar residues (e.g., {L, I, V}) into single symbols. (The full implementation of this P-C method is detailed in Methods).

#### 2.1.2 Stage 2: Initial Scoring for Functional Correlation

After encoding, **Stage 2** of the pipeline generates an initial, or “raw,” importance ranking to test for **Criterion 1 (Functional Correlation)**. For supervised tasks, we measure the association between a position *X_i_* and a target property *Y* using Mutual Information:

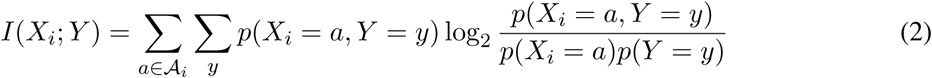

For unsupervised tasks, we quantify sequence variability using Shannon entropy:

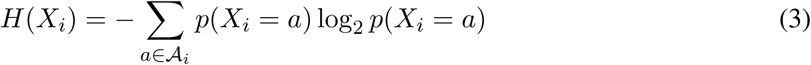

The theory is such that in the absence of labelled data, residue locations with low variability (low entropy) are more likely to be critical to maintenance of functionality, and thus less amenable to tuning. Thus in the absence of labels we use unsupervised entropy based measures as a proxy for functional correlation. However, regardless of the encoding used, these initial scores are still confounded by evolutionary coupling. They fail to separate true functional information (*I*(*s_j_*; *f*)) from structural redundancy (*R*(F)), which necessitates the pipeline’s second key hypothesis.

#### 2.1.3 Stage 3: Decoupling to Isolate Informational Hubs

The pipeline’s core innovation is **Stage 3 (Decoupling)**, which explicitly tests for **Criterion 2 (The Informational Hub)**. To solve the *R*(F) term in our objective function, we developed an iterative algorithm to isolate each residue’s independent predictive power. First, we construct a coupling matrix **C** using one of three measures: Cramér’s V (symmetric association):

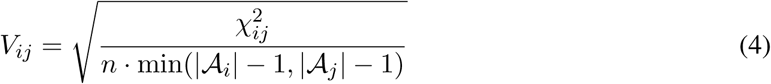

Theil’s U (asymmetric predictive power):

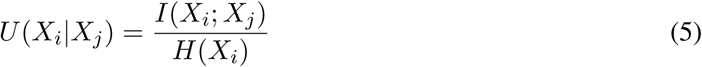

or Statistical Coupling Analysis (SCA). We then apply an iterative procedure, analogous to “network deconvolution” [12], to refine the initial scores. Starting with the highest-ranked residue *i*, we leave its score unchanged. For every other residue *j*, we reduce its score *r_j_* in proportion to its coupling with *i*: 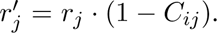 This is repeated down the ranking, ensuring each final score is adjusted to remove contributions already explainable by correlations with higher-ranked residues.

#### 2.1.4 Stage 4: Evaluation

Finally, **Stage 4 (Evaluation)** assesses the resulting “decoupled importance fingerprints”. We thus present the results of this pipeline, beginning with the effect of the decoupling algorithm itself.

### 2.2 Decoupling as Denoising

From a signal-processing perspective, residue importance traces resemble noisy signals in which functionally important positions appear as peaks amid background fluctuations. Evolutionary coupling introduces correlated noise: substitutions at one position induce variation at others through co-evolutionary constraints. This effect is evident in protein families with strong co-evolutionary signatures, where coupled fluctuations can obscure genuine functional signals.

To quantify the impact of decoupling, we characterized noise properties of importance traces before and after decoupling across all datasets, encoding schemes (standard categorical and physicochemical clustering), and information measures (e.g. Shannon entropy, Tsallis entropy etc.). Figure 3d illustrates a representative transformation: raw entropy exhibits a diffuse, broad distribution, while decoupled traces show sharper, localized peaks.

**Figure 3:**
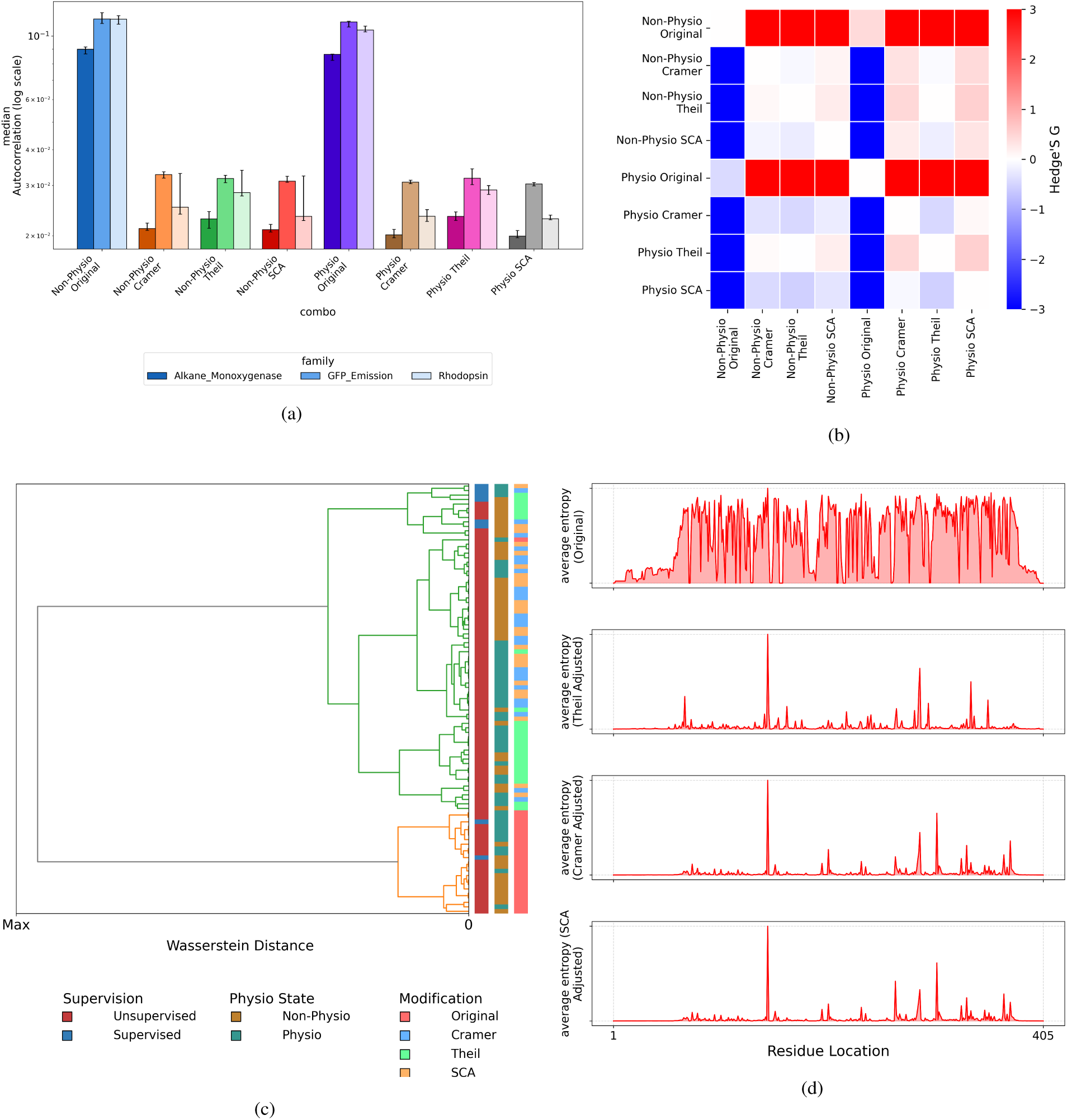
Decoupling improves performance across families and measures. a Cross-validated performance across three families. Decoupled methods outperform non-decoupled counterparts, with largest gains in the Alkane Monooxygenase dataset. b Effect sizes for physicochemical vs. standard encoding. Positive values in columns indicate physicochemical advantage, strongest for decoupled methods (Hedges’ *g >* 2 when comparing to base measures). c Method similarity clustering. Decoupled methods form distinct cluster (top), separate from non-decoupled (central), indicating characteristic signal transformation. Bars on the right show supervision mode (supervised in red vs. unsupervised in blue), encoding type (physicochemical in blue vs. standard in yellow), and decoupling status (decoupled in green, blue and yellow vs. non-decoupled in pink). d Shannon entropy transformation for GFP. Raw entropy (top) shows diffuse peaks; Theil’s U (2nd), Cramer’s V (3rd) and SCA (bottom) transforms sharpen peaks and reduce noise. SCA produces nearly identical results to Cramér’s V. Decoupling provides significant gains (paired *t*-test: *p <* 0.01, Hedges’ *g >* 5.8, *n* = 3 families).

This visual “denoising” translated into a large and consistent statistical effect. Decoupling improved signal quality across all protein families and information measures (Figure 3a), with the largest gains in the diverse Alkane Monooxygenases dataset. The magnitude of this improvement was exceptionally large, with Hedges’ *g* values exceeding 5 for all decoupling approaches (n=3 families) (Figure 3b). This effect size, which exceeds Cohen’s benchmark for “large” effects (*g >* 0.8), shows that decoupling has a large and practical impact. This gain was consistent across encoding choices: while physicochemical clustering offered some baseline improvements, both encoding schemes benefited greatly from the decoupling process.

Hierarchical clustering of the methods themselves shows why decoupling works (Figure 3c). The methods do not group by information measure (e.g., shannon entropy vs. Tsallis entropy), but by their processing pipeline. All non-decoupled methods—regardless of encoding or supervision—cluster together, supporting our hypothesis that they all suffer from the same correlated noise. In contrast, the decoupled methods form a distinct super-cluster, which then separates into logical branches based on decoupling type (Theil’s U vs. Cramér’s V) and encoding. This separation shows that decoupling is the most important processing step: it removes the shared, dominant noise, allowing the more subtle differences between information measures and encodings to emerge.

### 2.3 Decoupled Rankings Strongly Agree with Literature-Annotated Functional Sites

Key tuning residues control functional variation across protein families: mutations at these positions drive functional diversity, while changes elsewhere are neutral or destabilizing. If our hypothesis is correct, our information-theoretic methods should identify these residues from sequence data alone. We tested this prediction by comparing our rankings against literature-reported functional sites in three diverse protein families.

These benchmark families present varying conditions that test method robustness. First, annotation scope varies: Alkane Monooxygenase residues were mapped in a single homolog [13], Fluorescent Protein residues in GFP and close variants [14–18], while Rhodopsin residues represent family-wide consensus positions [19]. Second, alignment quality differs substantially: Rhodopsins maintain high conservation (dense, high-quality alignment), Fluorescent Proteins show moderate divergence (moderate alignment quality), while Alkane Monooxygenases span diverse lineages with ambiguous homology (sparse, challenging alignment). (Quantitative alignment metrics are in Supplementary document S1).

To evaluate prediction quality, we developed a Wasserstein Concentration Score (WCS), as standard metrics like AUROC and AUPRC fail to reward “near misses” (see Supplementary document S4 for AUROC/AUPRC comparisons, where our methods also outperform). The WCS combines two measures: (1) concentration, quantifying what fraction of the total importance score is captured by the top *N* literature-annotated positions, and (2) distributional distance, measuring how closely the importance distribution matches an ideal signal at those key residues (defined in Methods, Eq. 3). The final score, WCS = concentration*/*(1 + *W_d_*) (where *W_d_*is the Wasserstein distance), rewards methods that both concentrate predictive power in a small subset of residues and correctly localize that subset to the functionally relevant positions. Given our test data, we expected strongest performance on Rhodopsins (family-wide annotations, high-quality alignment) and weakest on Alkane Monooxygenases (single-species annotations, poor alignment).

Across all three datasets, our developed methods from both physicochemical and standard encoding pipelines outperformed existing tools (Figure 4). We observed large effect sizes, with Hedges’ *g* ranging from 0.6 to 0.9 (95% CI: 0.47–0.9, *p* = 0.002 to 0.1) when comparing our best methods to the best-performing benchmark tool (Figure 4c). As predicted, this performance scaled with both annotation and alignment quality: Rhodopsin (max WCS = 0.48) exceeded Fluorescent Proteins (0.43) and Alkane Monooxygenases (0.35). While the absolute WCS values (all *<* 0.5) reflect the reality of incomplete experimental annotations where most positions remain unstudied, the relative outperformance of our methods is clear. The Alkane Monooxygenase family represents a worst-case scenario for any sequencebased method, yet even here, our methods showed a positive improvement over existing baselines.

**Figure 4:**
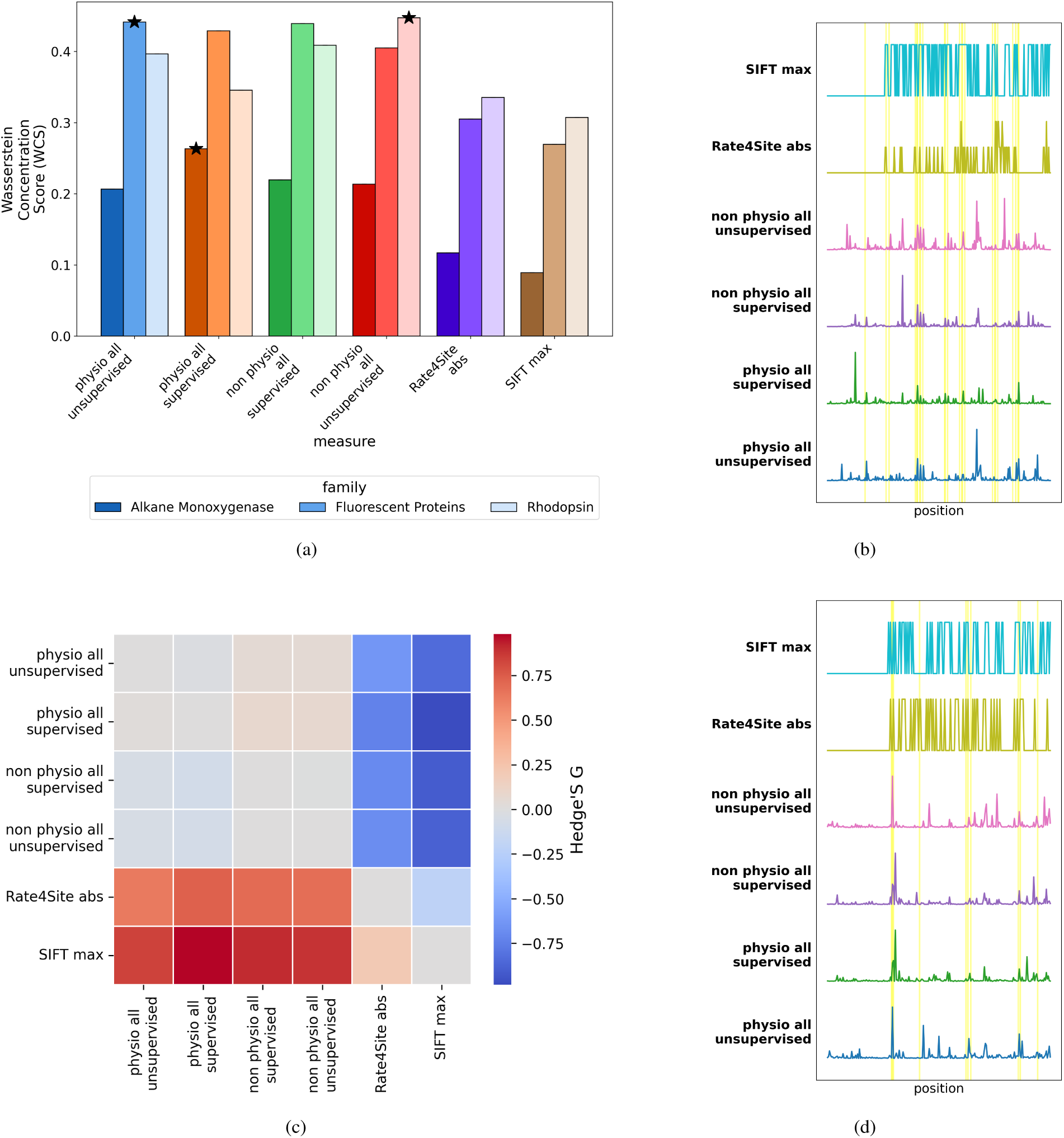
Developed methods outperform existing tools despite incomplete annotations. Literature-based validation against experimentally validated key residues (15±8 per family). a Wasserstein Concentration Scores (WCS) across three families. All methods achieve modest absolute scores (*<* 0.5), but developed methods exceed existing tools, especially in well-characterized Rhodopsins and in fluorescent proteins. b Importance traces for Rhodopsins. All top methods localize known functional regions (yellow bands), but developed methods produce sharper, more discriminative peaks. c Pairwise effect sizes. Warm colors in columns indicate consistent outperformance; developed methods cluster in high-performance region (top-left). d Importance traces for Fluorescent Protein. All top methods localize known functional regions (yellow bands), but developed methods produce sharper, more discriminative peaks.

### 2.4 Decoupled Rankings are More Informative for Machine Learning

To complement literature-based annotations, we define a data-driven ‘informativeness’ metric that directly quantifies how well residue rankings predict functional properties. For each ranking, we construct nested feature sets by progressively including residues in ranked order. A Random Forest regressor trained on each subset yields a performance curve (Fig. 5a). Optimal rankings produce steep initial gains followed by a plateau, indicating that top-ranked positions capture the most predictive signal. We quantify ranking quality as the area under this performance curve (AUPC) after enforcing weak monotonicity, yielding a score from 0 (poor) to 1 (ideal).

**Figure 5:**
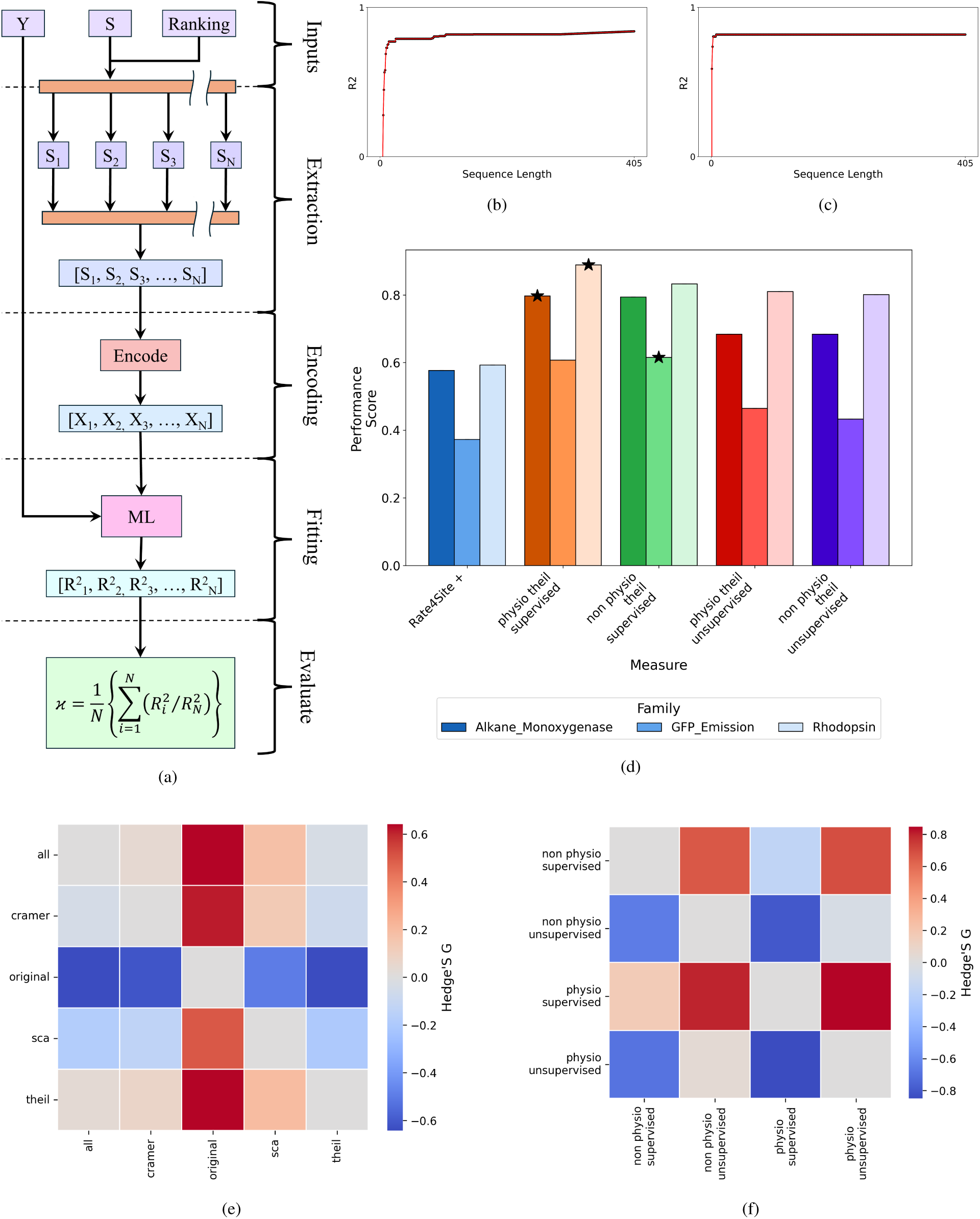
Top-ranked residues from developed methods concentrate functional information. Machine learning validation using Random Forest regressors trained cumulatively on top-ranked residue subsets. All methods evaluated identically using ESM2 embeddings. a Pipeline schematic: ranking quality quantified by area under *R*^2^ curve. Ideal methods produce steep initial rises. b Rate4Site (existing tool): slower increase indicates poorer discovery. c Supervised (decoupled and Physicochemically encoded): steep rise reaching *R*^2^ *>* 0.8 with *<* 5% of residues, demonstrating successful discovery. e Impact of decoupling: consistent gains across all families and metrics with low effect sizes when comparing different decoupling methods. f Effect sizes for modifications, examining the impact of supervision and physicochemical encoding. Supervision provides consistent gains; physicochemical encoding benefits primarily under supervision

For this validation, we used static ESM2 embeddings as a uniform, fixed feature set for all methods. This ensures a fair comparison that tests only the quality of the residue ranking, not the feature encoding itself (see Methods). For instance, with fluorescent proteins, our best method (Theil Adjusted Mutual Information Physio) achieved an *R*^2^ ∼ 0.80 using only the top 11 residues (Fig. 5c). In contrast, the best existing tool (Rate4Site) required 19 residues to reach comparable performance (Fig. 5b). Across all three protein families, our methods outperformed existing tools (Fig. 5d). Our supervised approaches showed large effect size improvements over Rate4Site (Hedges’ *g* = 1.64 − 1.83, 95% CI: 1.61-1.79, *n* = 3 families), and our best unsupervised approach also achieved intermediate gains (Hedges’ *g* = 0.66 − 0.67, 95% CI: 0.56-0.61).

Having established this performance gain, we performed ablation analyses to isolate and quantify the contributions of our two key innovations: (1) *decoupling* to remove confounding evolutionary background signals, and (2) *physicochemical encoding* to replace categorical amino acid representations.

1. **Decoupling provides the largest individual gain.** Isolating the decoupling component revealed performance gains (Hedges’ *g* = 0.5–0.65) across all protein families and metrics (Fig. 5e). Removing evolutionary coupling improves performance, independent of the encoding scheme or base metric.
2. **Physicochemical encoding requires supervision.** Substituting categorical encodings with physicochemically derived categories yielded modest, supervision-dependent improvements (Fig. 5f). The benefit was evident under supervision (Hedges’ *g* ≈ 0.3) but negligible for unsupervised methods, suggesting these features primarily enhance learning when guided by functional labels.
3. **Supervision and encoding interact.** As expected, supervised models consistently outperformed their unsupervised counterparts (Hedges’ *g* ≈ 0.7), confirming the value of functional annotations (Fig. 5f). This benefit was amplified by the physicochemical encoding, indicating a positive interaction between supervision and biochemically informative representations.
4. **Combined effects exceed additive expectations.** The strongest effect was observed when combining both innovations under supervision. The fully integrated model (decoupled, physicochemical, supervised) outperformed the baseline (non-decoupled, categorical, unsupervised) with a large effect size (Hedges’ *g* = 1.15). This combined gain exceeds the sum of the individual component effects: decoupling suppresses evolutionary noise, while physicochemical encoding captures functional relationships obscured by categorical representations.

### 2.5 Quantifying the Interplay of Supervision and Alignment Quality

To characterize the relationship between supervision and data quality, we quantified the performance of supervised versus unsupervised methods in two distinct contexts: a single, densely-aligned protein family and a set of diverse, sparsely-aligned families.

First, to isolate the effect of supervision, we analyzed a high-quality, densely-aligned family (Green Fluorescent Protein) across three target properties (Fig. 6c). In this setting (Fig. 6e), supervised methods consistently outperformed their unsupervised counterparts. The unsupervised scores were approximately one-third lower, indicating that access to functional labels provides a performance benefit even when sequence data is abundant and homology is clear.

**Figure 6:**
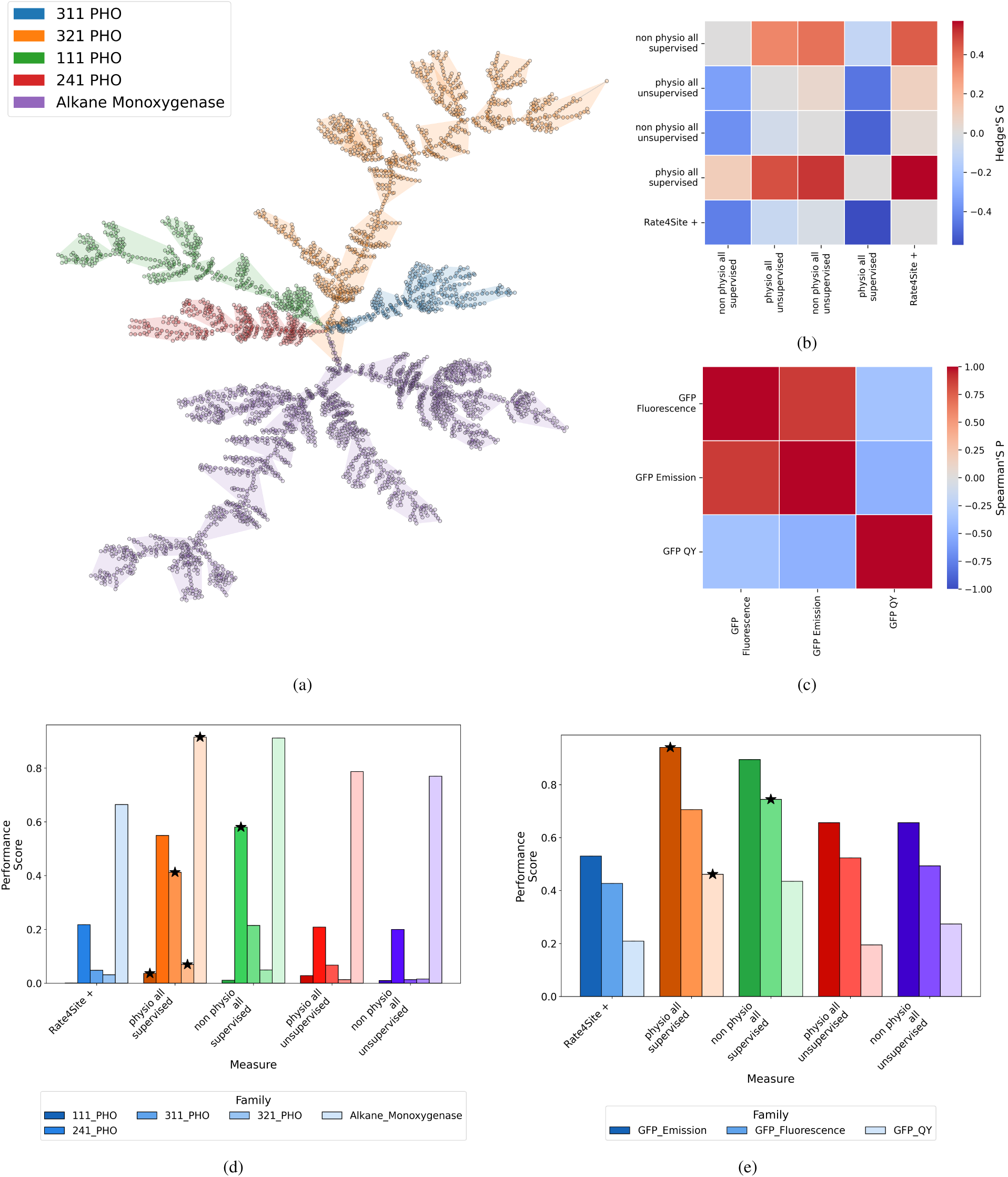
Alignment quality determines supervised vs. unsupervised method choice. Enzyme families (sparse alignments) vs. GFP multi-property analysis (dense alignment). a Enzyme family tree shows large divergence and sparse coverage—challenging for unsupervised methods. b Supervised advantage increases with family divergence (warm colors). Even unsupervised decoupled methods exceed existing tools (*g* ≈ 0.5). d Family-specific distributions confirm systematic supervised advantage in divergent families. c GFP property correlations: fluorescence-emission highly correlated (*r >* 0.8), quantum yield weakly correlated. e In well-aligned GFP, unsupervised (orange) matches supervised (blue) across all properties. Alignment density, not property type, determines method choice.

We then quantified this performance gap in a more challenging, sparse-alignment context using diverse enzyme families (Fig. 6a). Here, the performance gap between supervised and unsupervised methods widened considerably. The unsupervised method performance was typically less than half that of the supervised counterparts (Fig. 6b, 6d). Despite this large gap, our unsupervised decoupled methods still achieved large average effect sizes (*>* 0.5) over the leading existing tool, Rate4Site, showing a clear baseline improvement from our framework.

Together, these two analyses provide a quantitative guideline. The GFP analysis, by holding alignment quality constant, isolates the independent value of supervision. The enzyme analysis confirms that the magnitude of this performance gap is strongly modulated by data quality, widening considerably as alignment quality degrades.

### 2.6 Case Study: Decoupled Rankings Identify Key Binding Residues in SARS-CoV-2 Nanobodies

To validate our framework on a practical binding problem, we analyzed a dataset of nanobodies targeting the SARS-CoV-2 spike protein. This system provides a clear benchmark: we can compare the predictions of our sequence-only method to an independent, structure-based “ground truth” derived from computational docking (ClusPro 2.0 [20]). The hypothesis is that our method, without 3D information, can enrich for the same interfacial and energetically-critical positions identified by the structural approach.

We first established this structural ground truth via molecular docking simulations. We defined two reference sets: (1) Interfacial residues (48 total), those within 10 Å of the antigen surface (Figure 7a, purple); and (2) Key binding residues (20 total), a subset of the interfacial positions where *in silico* mutation scanning showed a large shift in binding energy (Figure 7a, red). These 20 residues represent the most important sites for binding.

**Figure 7:**
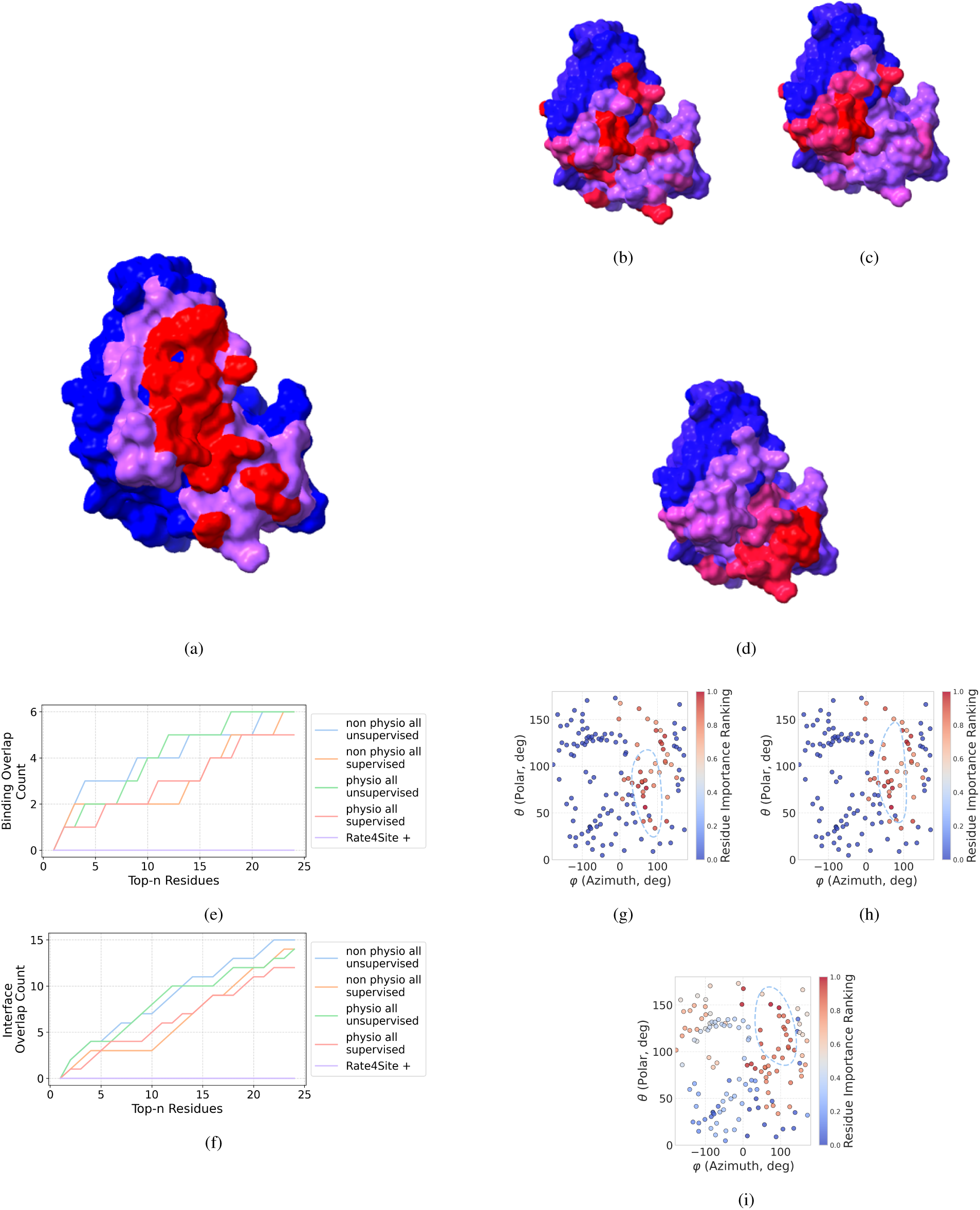
Sequence-only predictions recover 67% of binding-critical residues. Nanobody-SARS-CoV-2 validation with ground truth from docking and mutational scanning. a Reference: interfacial region (purple, 10 Å), binding-critical residues (red). b–d Top-ranked residues (red spheres): example developed methods b–d cluster at interface, while Rate4Site, d identifies distal surface. e Discovery curves: within top 10% (24 positions), developed methods capture 5+ residues, Rate4Site zero. f Interfacial analysis: developed methods 12+ positions, Rate4Site zero. g–i Surface projections (red = high, blue = low): developed methods show coherent interface patches

Visual inspection of the results (Figure 7) and their 3D surface projections (Figures 7g–7i) shows a clear distinction. Our developed methods (e.g., hard voting, KL divergence) concentrate high-importance scores on the binding interface, consistent with the structural ground truth. In contrast, Rate4Site assigns its highest importance to a distal surface cluster, a prediction inconsistent with the energetic and spatial evidence.

We quantified this by analyzing the top 10% of ranked positions (24 residues) from each method. Our developed methods consistently identified 5 or more of the 20 key binding residues (a sensitivity of ≥25%). The best existing method, Rate4Site, identified zero (Figure 7e). This enrichment is statistically significant: the probability of finding 5 or more of the 20 key sites in a random draw of 24 (out of 240) is *p* = 0.031 (hypergeometric test). A parallel analysis for the 48 interfacial residues showed the same pattern, with our methods recovering 12+ positions (¿50% sensitivity) while Rate4Site again identified none (Figure 7f).

These results have direct practical implications for reducing experimental burden. A comprehensive saturation mutagenesis of the 240-residue nanobody would require testing ∼4,560 variants. A structure-guided approach, testing only the 48 interfacial residues, still requires ∼912 variants. Our sequence-only method, by ranking the top 24 positions, identifies a subset of these sites while requiring only ∼456 variants. This represents a 90% reduction in experimental effort versus the comprehensive scan and a 50% reduction versus the structure-guided approach. This provides a sequence-only filter to focus experimental efforts, especially when structural information is unavailable.

## 3 Discussion

We have reframed key residue identification as an information-theoretic feature selection problem. Our central advance is a **training-free, decoupling framework** that isolates functional signals from evolutionary noise. This offers a distinct approach from many modern ML tools (e.g., AlphaFold, ESM) which, while useful for structure prediction or variant scoring, do not explicitly identify family-wide functional *tuning sites*. Our method provides mechanistic interpretability and explicit coupling matrices, all derived from sequence data alone.

### 3.1 Why Decoupling Works: The Informational Hub Hypothesis

Our framework’s success appears to hinge on solving the problem of *confounded coupling*. We hypothesized that key tuning residues act as “informational hubs” that predict the state of other residues. However, this signal is normally obscured by structural co-evolution. Our iterative decoupling algorithm acts as a “denoising” filter that isolates informationally unique positions.

The nanobody case study provides support for this. Our decoupled rankings enriched for residues that are (1) clustered at the binding interface and (2) energetically important for binding, as validated by independent computational docking. This suggests our information-theoretic approach is identifying positions that are biophysically and functionally relevant.

### 3.2 Practical Guidelines for Method Selection

Our analyses suggest clear guidelines for researchers:

- **Supervision provides a consistent benefit:** When functional labels are available, supervised methods (Mutual Information) provide a performance benefit. We observed this even in high-quality, dense alignments (like GFP, where scores improved by ∼33%) and noted the gap widens in sparse, diverse alignments (enzymes, where supervised scores were often *>* 50% higher).
- **Unsupervised methods are a viable proxy:** When labels are unavailable, our *unsupervised* decoupled methods still outperformed existing benchmarks (like Rate4Site). This makes them a reliable tool for exploratory analysis, especially for well-aligned families.

### 3.3 A Path to Reduced Experimental Burden

While the absolute prediction accuracy for any sequence-based tool remains a challenge (often due to sparse experimental validation), the *relative* gain of our method has practical implications.

In the nanobody case study, our method identified **over 25% (5/20) of the key binding residues** within the top 10% of its predictions, while the best existing tool identified none in that same tier. This ability to enrich for key sites can translate to a meaningful reduction in experimental cost.

For example, a comprehensive scan of the 240-residue nanobody requires ∼4,560 variants. A structure-guided approach, testing the 48 interfacial residues, still requires ∼912 variants. Our sequence-only method, by focusing on the top 24 positions (10%), requires only ∼456 variants. This represents a **90% reduction in effort versus a comprehensive scan and a 50% reduction versus a structure-guided approach**, while enriching for the most important binding sites.

### 3.4 Limitations and Future Work

This framework has limitations. First, it targets *tuning residues* (variable sites) and is not designed to find *absolutely conserved* catalytic sites. Second, our validation relies on literature annotations (which are sparse) and computational docking (which is not experimental ground truth).

Future work should focus on experimentally validating these predictions. The ultimate goal is to integrate this framework into engineering workflows. The importance scores and coupling matrices provide a natural foundation for **rational library design**, potentially formalizing directed evolution as a guided Markov process. By using our rankings to focus experimental mutagenesis, this work provides a direct, sequence-only path to more efficient protein engineering.

### 3.5 Conclusion

Reframing residue identification as hypothesis-driven feature selection yields a generalizable framework that matches or exceeds black-box model performance while maintaining full interpretability. We establish and validate a unified theory of key tuning residues as informational hubs, providing both immediate utility for protein engineering and a rigorous foundation for machine learning-guided evolution.

Our results demonstrate consistent and statistically well-supported improvements across all evaluated metrics. When looking at performance in downstream regressors, supervised methods achieved Hedges’ *g* = 0.83 (95% CI: 0.47 to 1.40, *p* = 0.00013) for physicochemical-encoded approaches and *g* = 0.71 (95% CI: 0.40 to 1.18, *p* = 0.00034) for standard-encoded variants when compared to the best existing tool (Rate4Site). Unsupervised methods also demonstrated meaningful but lesser improvements (*g* ≈ 0.2, *p* ≈ 0.02-0.03). Additionally, decoupling alone provided large gains (Hedges’ *g* = 5.8-6.2 with 95% CI: *g* = 4.8-5.3, *n* = 3 families), demonstrating the importance of removing evolutionary coupling. In the nanobody-antigen binding case study, our methods identified ¿25% of binding-critical residues (6/20) within the top 10% of predictions (hypergeometric test, *p* = 0.031), compared to zero recovered by the best existing method, illustrating practical utility for protein engineering applications.

The framework’s flexibility, operating in both supervised and unsupervised modes, ensures its broad applicability. It provides explicit coupling matrices and mechanistic interpretability—features often absent in ‘black-box’ ML models. This enables a direct, sequence-only path to more efficient protein engineering by helping to focus experimental mutagenesis and providing a foundation for rational library design. As sequence databases grow larger and experimental resources remain limited, our transparent, efficient mathematical approach provides a useful tool for translating evolutionary information into functional insight.

## 4 Methodology

### 4.1 Test Datasets

We used seven non-structural datasets: three diverse protein families and four enzyme families. These datasets include protein sequences and corresponding target variables. All datasets were fully labeled, meaning no sequences had missing functional data.

- **Diverse Families:** Rhodopsins (884 sequences) [19], GFP variants (611 sequences) [14], and Alkane Monooxygenases (1880 sequences) [21]. Rhodopsins and GFP datasets were obtained from published literature, while the Alkane Monooxygenase dataset was compiled from a BLAST search around a reference protein with experimentally tested key tuning residues [13, 21–30]. For alkane monooxygenases we used deep learning predicted optimal pH [31] as the target variable owing to data availability. Table 1 summarizes these three families, their target variables, known key residues from literature (ground truth), and residues predicted by our best-performing supervised method.
- **Enzyme Families:** We selected four additional enzyme families with varying alignment quality and sequence diversity from the BRENDA database, requiring at least 300 sequences per EC class (first three digits), a known functional property, and complete accession numbers. The four families were: EC 1.1.1 (Aldehyde Dehydrogenases, 496 sequences), EC 2.4.1 (Glycosyltransferases, 381 sequences), EC 3.1.1 (Carboxylesterases, 343 sequences), and EC 3.2.1 (Glycoside Hydrolases, 1,123 sequences). Optimal pH was used as the target variable for all enzyme families owing to data availability in the BRENDA database [32].
- **Nanobodies:** This dataset comprises 3,127 unique nanobody sequences with measured binding to the SARS-CoV-2 spike protein [33].

**Table 1:** Summary of Protein Families and Target Variables with Key Residues. Column ‘Key Residues’ lists residues identified as functionally important in experimental literature (ground truth). Column ‘Supervised Physio Decoupled’ lists the top-ranked residues predicted by our supervised mutual information method with decoupling and physiochemical encoding.

| **Protein Family** | **Target Variable** | **Key Residues (Ground Truth)** | **Citations** | **Size** | **Supervised Physio Decoupled** |
| --- | --- | --- | --- | --- | --- |
| Rhodopsins | Absorption wavelength | M52, V80, A84, Y119, D121, W122, T125, V126, L129, M158, I159, G162, G178, S181, T182, F185, W222, Y225, P226, Y229, Y249, D253, A256, K257 | [19,22,36] | 884 | F38, N43, W95, L101, S110, K112, D121, T125, L129, E143, S168, T172, T173, L200, V201, R202, S237, V243, V245, D253, A256, L269, D275, H282 |
| GFP and Variants | Excitation Wavelength | Q68, Y69, G70, R97, W145, K147, E150, H199, I201, Q219 | [14–18,22] | 611 | Q68, N71, R72, L127, V163, E165, H199, G207, M214, P227 |
| Alkane Monooxygenases | Optimum pH | H44, H45, C106, N113, S227 | [13,21–30] | 1880 | L34, P120, Q124, E248, K263 |

### 4.2 MSA Generation and Benchmark Tool Execution

For each protein family, a single Multiple Sequence Alignment (MSA) was generated using Kalign (v3.3.1) [34] with automatically determined parameters. Gap characters were treated as a distinct 21st amino acid category in all entropy and mutual information calculations; alignment columns consisting entirely of gaps were removed prior to analysis. **Critically, to ensure fair comparison across all methods including the nanobody case study, this identical MSA was used as direct input for all sequence-based methods evaluated: our framework and five benchmark tools (DeepSequence, Rate4Site, EVmutation, HotspotWizard, and SIFT).** This unified alignment input ensures that performance differences reflect algorithmic choices (entropy vs. mutual information, decoupling strategies, feature selection) rather than preprocessing variations. For the nanobody-antigen binding case study specifically, the same MSA alignment used for our sequence-based rankings was employed to extract alignment-based features for Rate4Site and other sequence-only baselines, confirming methodological consistency. Code to reproduce all alignments and tool executions is provided in the GitHub repository.

### 4.3 Execution Of Existing Tools

We used the following tools as benchmarks:

- **DeepSequence** (v0.2.6) [11] was run using default parameters.
- **Rate4Site** (v3.0.0) [9] was run using default parameters.
- **EVmutation** (v0.1.1) [10] was run using default parameters.
- **SIFT** (v5.1.1) [8] was run using default parameters.
- **HotspotWizard** (v3.0) [7] was run using default parameters.

Where multiple scores were produced per residue (e.g. EVmutation), we used the following pooling methods: (1) mean score, (2) max score, (3) min score, (4) mean of absolute scores, (5) max of absolute scores, (6) min of absolute scores. The best-performing pooling method for each tool was used in the final analysis. Code to reproduce all tool executions is provided in the GitHub repository, and principles behind each tool are outlined in Supplementary Proofs S5.

### 4.4 Information Theoretic Analysis

Our framework’s first stage involved ranking residues using information-theoretic measures computed directly from sequence alignments-**this is a purely mathematical, training-free approach requiring no parameter optimization or learning from functional labels**. In the supervised case, we used **Mutual Information (MI)** to directly compute statistical associations between sequence positions and functional properties. In the unsupervised case, we employed **Hard Voting as our primary method**: an ensemble approach that aggregates rankings from multiple entropy-based measures to enhance robustness.

**Hard Voting Algorithm** treats the problem as a mixture of experts, where each entropy measure provides an independent ranking of residues. The algorithm proceeds in five steps:
list
1. **Score Computation**: Compute importance scores 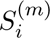 for each residue *i* across all *M* component entropy measures *m* ∈ {1*, . . ., M* } (e.g., Shannon, Renyi, Tsallis, KL divergence, etc.) independently from the sequence alignment.
2. **Per-trace Normalization**: Normalize each measure’s scores to preserve per-measure relative importance while equalizing contribution magnitudes: 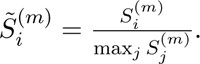 This ensures no single measure dominates due to scale differences while maintaining rank ordering within each measure.
3. **Rank Conversion**: Convert normalized scores to ranks within each measure. A residue with the highest normalized score receives rank 1 (best), the second-highest receives rank 2, etc. Ties are handled via secondary rank assignment based on original score ordering. Formally, 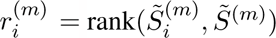 where ties broken by score value.
4. **Majority Voting with Tiebreaking**: For each position *i*, count how many measures assign each rank value. Identify the mode (most frequent rank). In case of ties between rank values with equal vote counts, break ties using the median score from all measures that voted for any tied rank: 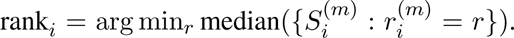 This tiebreaker prioritizes ranks supported by higher-scoring measures while averaging out measure-specific noise.
5. **Final Normalization**: Normalize the aggregated ranks to a continuous score in the range [0, 1] by inverting rank (low rank → high score): 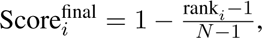 where *N* is the total number of residues. This produces a final importance ranking combining evidence across all measures.

#### Implementation Details

The ensemble aggregates 11 different entropy-based measures (see list below), ensuring stable performance through wisdom-of-crowds principles. Each measure is computed independently without learned parameters. Statistical significance of the aggregated ranking is assessed via bias-corrected, non-parametric bootstrap with 10,000 resamples to compute 95% confidence intervals around effect sizes. Complete implementation is provided in `hard voting agglom.py` in the repository, which includes detailed debug output showing original scores, rank assignments, vote counts, tiebreaker logic, and final aggregated scores for full transparency.

The component measures in our Hard Voting ensemble include:

- **Shannon Entropy** A measure of uncertainty in a probability distribution.
- **Renyi Entropy** A generalization of Shannon entropy that introduces a parameter to adjust sensitivity to distribution tails.
- **Tsallis Entropy** Another generalization of Shannon entropy, useful for non-extensive systems.
- **Kullback-Leibler Divergence** A measure of how one probability distribution diverges from a second, expected probability distribution.
- **Neumann Entropy** A quantum analog of Shannon entropy, applicable to quantum states.
- **Gini Impurity** A measure of statistical dispersion intended to represent the income inequality within a nation or a social group.
- **Kolmogorov Complexity** A measure of the computational resources needed to specify a dataset.
- **Jensen Shannon Divergence** A method of measuring the similarity between two probability distributions.
- **Average Entropy** A measure of the average uncertainty across a distribution.
- **Soft Voting** A weighted voting scheme where measures contribute according to their confidence.

These component measures represent diverse information-theoretic concepts including uncertainty (Shannon, Renyi, Tsallis), divergence from reference distributions (Kullback-Leibler, Jensen-Shannon), impurity (Gini), and complexity (Kolmogorov). When combined with decoupling, Hard Voting provides consistent performance (Supplementary Figures S1.9-S1.10). **Throughout this manuscript, “our method” or “developed method” refers to Hard Voting with decoupling as the primary approach**, with individual component measures presented in supplementary materials for completeness. For detailed mathematical definitions and rationales behind each component measure, see Supplementary Proofs S5.

### 4.5 Input Encoding Schemes

We compared two distinct input encoding schemes:

1. **Base Sequence Encoding:** The standard categorical representation of the 20 amino acids.
2. **Physicochemical Pre-encoding:** A novel scheme where each amino acid is represented as a vector of 16 physicochemical properties. Then, for each MSA position, residues are clustered using k-means, with the optimal number of clusters determined automatically by maximizing the silhouette score.

The physicochemical properties used for encoding were selected based on their relevance to protein structure and function, including hydrophobicity, charge, polarity, molecular weight, and others. This encoding aims to capture underlying biochemical similarities between amino acids that may not be evident in a purely categorical representation. Clustering residues at each position helps to reduce noise and highlight functionally relevant patterns.

#### Detailed Amino Acid to Physicochemical Cluster Conversion

Each of the 20 canonical amino acids is represented by a 16-dimensional feature vector encoding key physicochemical descriptors. These properties include:

- **Continuous properties**: Molecular weight (g/mol), charge at pH 7.4, isoelectric point, hydropathy index (Kyte-Doolittle scale), volume (Å ^3^), bulkiness (Zimmerman), flexibility (Vihinen), hydrogen bond donors (sidechain), hydrogen bond acceptors (sidechain).
- **Binary categorical properties**: Is aliphatic (A, G, I, L, V), is aromatic (F, W, Y), is polar (N, Q, S, T), contains sulfur (C, M), is acidic (D, E), is basic (H, K, R), is cyclic (P).

For example, leucine (L) is encoded as:

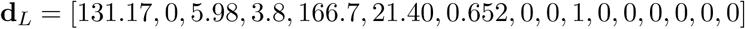

while aspartic acid (D) is:

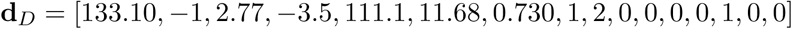

The clustering algorithm operates position-wise on the MSA:

1. **Feature extraction**: For position *i* in the MSA with *N* sequences, extract the amino acids 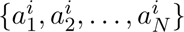 and map each to its 16-dimensional property vector, forming a feature matrix **X***^i^* ∈ ℝ*^N^* ^×16^.
2. **Preprocessing**: Handle gaps and unknown residues by median imputation, then standardize features using z-score normalization:

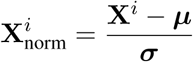 where ***µ*** and ***σ*** are computed column-wise across the *N* sequences.
3. **Optimal cluster determination**: Apply k-means clustering for *k* ∈ [2*, k*_max_] (default *k*_max_ = 10) and compute silhouette scores to determine optimal cluster count:

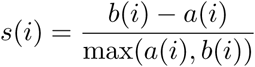 where *a*(*i*) is the mean intra-cluster distance and *b*(*i*) is the mean nearest-cluster distance for sample *i*. The optimal *k*^∗^ maximizes the average silhouette score across all sequences at position *i*:

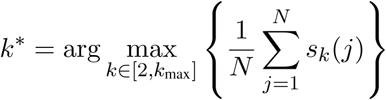

This approach provides a unified, simplified framework with theoretical guarantees for convergence and cluster quality. Detailed algorithmic specifications, including initialization strategies and convergence criteria, are provided in Supplementary Section S5.
4. **Cluster assignment**: Apply k-means with the optimal *k*^∗^ to partition amino acids at position *i* into biochemically similar groups. Each amino acid is replaced by its cluster label *c* ∈ {1, 2*, . . ., k*^∗^}.
5. **Position-specific encoding**: The cluster assignments effectively reduce the alphabet size from 21 (20 amino acids + gap) to typically 2-6 clusters per position, with the exact number determined by the biochemical diversity observed at that site.

This position-specific clustering adapts to local biochemical constraints: conserved catalytic sites often collapse to 2-3 clusters (capturing strict chemical requirements), while variable surface loops may retain 5-7 clusters (reflecting tolerance for diverse sidechains). By grouping biochemically equivalent substitutions-such as L↔I↔V (hydrophobic, aliphatic, similar volume)-the encoding reduces noise from functionally neutral mutations while preserving signal from chemically consequential changes like D↔K (charge reversal). This improves downstream coupling analysis by reducing spurious correlations arising from arbitrary symbolic differences between biochemically similar residues. For detailed property values and validation of clustering stability across families, see Supplementary Table S5.1.

### 4.6 Removing the Effect of Coupling

To test Criterion 2 of our hypothesis (Informational Hub), we employ a decoupling algorithm. We use three complementary measures for this. **Theil’s U** is used as a measure of predictive power, assessing how much the uncertainty of one residue’s identity is reduced by knowing another’s. The **Chi-squared (***χ*^2^) statistic, from which Cramér’s V is derived, is used as a symmetric measure of association. **Statistical Coupling Analysis (SCA)** quantifies the coevolutionary relationship between positions through conservation-weighted correlation of amino acid distributions. These measures are computed for all pairs of residues in the MSA, resulting in a coupling matrix that quantifies the strength of association between every pair of positions. Theil’s U is directional, capturing how well position *i* predicts position *j*, while Cramér’s V and SCA are symmetric, reflecting mutual association without directionality. All three measures are bounded between 0 and 1 (or normalized to this range for SCA), ensuring numerical stability. Empirical comparison across our datasets demonstrates that SCA and Cramér’s V produce nearly identical effect sizes (Supplementary Figure SX), suggesting they capture similar aspects of pairwise coupling despite different mathematical formulations. Coupling matrix computation was GPU-parallelized to handle large protein families efficiently.

They are defined as follows:

- Theil’s U:

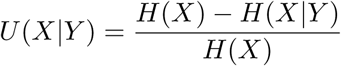

where *H*(*X*) is the entropy of variable *X* and *H*(*X*|*Y*) is the conditional entropy of *X* given *Y* .
- Cramér’s V:

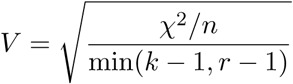

where *χ*^2^ is the chi-squared statistic, *n* is the total number of observations, and *k* and *r* are the number of categories for the two variables.
- Statistical Coupling Analysis (SCA):

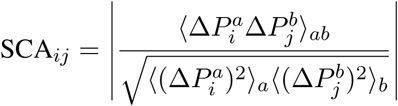

where 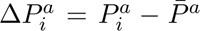 represents the deviation of amino acid frequency *a* at position *i* from the mean frequency across all positions, and ⟨·⟩ denotes averaging over amino acid types. This quantifies conservation-weighted covariation between positions.

The decoupling procedure operates iteratively to isolate each residue’s independent contribution. Residues are processed in descending order of their initial importance scores *S*(*i*), which are normalized at initialization to sum to unity. The ranking order is fixed at the start; residues are not re-ranked between iterations. At each iteration *t*, the top-ranked remaining residue *i_t_* retains its score unchanged, and all lower-ranked residues *j* have their scores adjusted to remove the component attributable to coupling with *i_t_*:

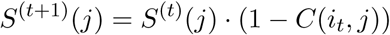

where *C*(*i_t_, j*) is the coupling measure (Theil’s U, Cramér’s V, or SCA) between residues *i_t_* and *j*. This multiplicative adjustment down-weights residue *j* proportionally to its coupling strength with *i_t_*, ensuring that subsequent rankings reflect unique functional contributions rather than redundant covariation.

We chose multiplicative down-weighting over alternative linear or subtractive schemes for three reasons: (1) it guarantees preservation of non-negativity (since 0 ≤ *C*(*i, j*) ≤ 1, we have 0 ≤ *S*^(*t*+1)^(*j*) ≤ *S*^(*t*)^(*j*)), avoiding artifacts from negative scores; (2) it produces cumulative attenuation across iterations-residues coupled to multiple high-ranked positions are progressively down-weighted in a compounding manner, naturally reflecting their redundancy; and (3) the normalization-preserving property (scores normalized at start remain proportionally valid after each multiplicative update) ensures interpretability throughout the iterative process. The procedure terminates after *N* iterations, yielding a decoupled importance fingerprint. Computational complexity is *O*(*N* ^2^) for coupling matrix construction (parallelized on GPU) and *O*(*N* ^2^) for the iterative decoupling procedure. For the largest family analyzed (1,880 sequences, 482 residues), total computation time was approximately 45 seconds on an NVIDIA A100 GPU. For detailed algorithms and convergence proofs, please see Supplementary Proofs S5.1.5.4.

### 4.7 Performance Evaluation

We evaluated methods using two complementary metrics: literature-based Coherence (assessing alignment with experimentally validated key residues) and data-driven machine learning informativeness (measuring predictive power for functional properties). Effect sizes were quantified using Hedges’ *g* as the primary statistical metric. Given the limited family count (n=3-7), which reduces statistical power for significance testing, we prioritize effect size reporting over p-values. We employed paired t-tests with prior normality assessment (Shapiro-Wilk tests), and report 95% confidence intervals on Hedges’ *g* values computed via bias-corrected bootstrap (10,000 resamples). All raw scores are configuration-specific: each combination of base measure (e.g., Shannon entropy, mutual information), encoding type (categorical or physicochemical), and decoupling type (none, Cramér’s V, Theil’s U, or SCA) produces a distinct score, enabling systematic evaluation of methodological modifications.formation), encoding type (categorical or physicochemical), and decoupling type (none, Cramér’s V, Theil’s U, or SCA) produces a distinct score, enabling systematic evaluation of methodological modifications. Where multiple pairwise comparisons were performed, p-values were adjusted using the Holm–Bonferroni method to control the family-wise error rate; reported p-values are Holm–Bonferroni-adjusted unless otherwise stated.

#### 4.7.1 Wasserstein Concentration Score (WCS) (mathematical definition)

Let *S* = (*S*_1_*, . . ., S_N_*) be the raw per-residue importance scores produced by a method for a sequence of length *N* . Let *K* = {*k*_1_*, . . ., k_n_*} denote the set of literature-annotated key residue indices.

**Step 1: Normalize scores to form a probability distribution** First, normalize the raw scores so they sum to unity:

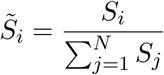

If all scores are zero or equal (degenerate case), assign uniform weights: *S̃_i_*= 1*/N* .

**Step 2: Compute concentration in top-***n* **positions** The concentration measures what fraction of the total importance mass is captured by the top *n* positions (where *n* = |*K*| is the number of literature-annotated key residues):

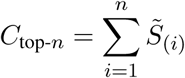

where 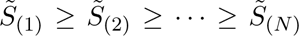 denotes the scores sorted in descending order. This quantifies how concentrated the importance signal is in the top-ranked positions.

**Step 3: Construct target pulse train** Define an ideal distribution that places unit weight at each key residue position:

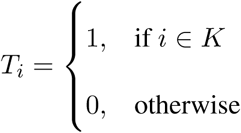

Normalize to form a probability distribution:

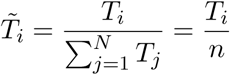

**Step 4: Compute Wasserstein distance** The Wasserstein distance (also known as Earth Mover’s Distance) measures the distributional difference between the predicted importance curve *S̃* and the target pulse train *T̃*:

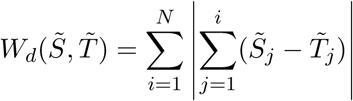

This quantifies how much “probability mass” must be moved to transform the predicted distribution into the ideal distribution.

**Step 5: Compute Wasserstein Concentration Score** The final WCS combines concentration and distributional accuracy:

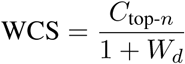

The denominator (1 + *W_d_*) penalizes methods whose top-ranked positions are far from the true key residues. High WCS values indicate both strong concentration of importance in a small subset of residues AND accurate localization of that subset to functionally relevant positions.

##### Interpretation

- WCS ∈ [0, 1], with higher values indicating better performance.
- If the top *n* predictions exactly match the literature-annotated residues, then *W_d_* ≈ 0 and WCS ≈ *C*_top-_*_n_*.
- If predictions are randomly distributed, *W_d_* is large and WCS approaches zero.
- The metric is robust to incomplete literature annotations: it rewards methods that concentrate signal in functionally relevant regions, even if annotations are sparse.

This metric directly tests our hypothesis that effective methods should both concentrate predictive power in a small subset of residues (captured by *C*_top-*n*_) and localize that subset to functionally critical positions (captured by the Wasserstein distance penalty *W_d_*).

#### 4.7.2 Machine Learning-based Informativeness (mathematical definition)

We assessed residue ranking informativeness by training Random Forest regressors to predict target functional properties using only top-ranked residue subsets. The procedure follows:

- Rank all residues by their importance scores (method-specific).
- Select the top *k*% of residues (evaluated adaptively across all k).
- Extract corresponding MSA columns to form a reduced feature set.
- Train a Random Forest regressor on this reduced feature set to predict the target variable.
- Evaluate performance using cross-validated *R*^2^.

This approach quantifies how effectively selected residues capture functional variability, providing an objective, data-driven ranking quality metric independent of literature annotations.

To encode categorical amino acid sequences for regression, we used pretrained per-residue embedding lookup tables from ESM2 (facebook/esm2 t33 650M UR50D) [35], converting each amino acid into a fixed-dimensional numerical vector. For a protein with *m* selected residues and *N* sequences in the MSA, this yields a feature matrix of dimension *N* × (*m* × *d*), where *d* is the embedding dimension. These embeddings capture rich evolutionary and biochemical patterns learned from large-scale protein databases, enabling effective Random Forest learning even with modest sample sizes. Complete details on ESM2 model selection, embedding dimension, and preprocessing steps are provided in Supplementary S5.1.7.

We used Random Forest regressors (100 trees, maximum depth 10) to balance model capacity against overfitting risk; hyperparameters were selected based on prior protein engineering studies and not tuned per-dataset. Cross-validation employed Monte Carlo sampling: 10-fold cross-validation with an additional held-out 20% test set, repeated across five independent random splits. Reported *R*^2^ scores represent mean performance across cross-validation folds. To emphasize early-ranking performance (most relevant for experimental prioritization), we computed a separate squared AUROC-style metric: AUROC^2^, derived from the area under the *R*^2^ vs. top-*k*% curve. This metric rewards methods that achieve high predictive power with minimal residue subsets.

To improve computational efficiency, we used adaptive segment sampling: the ranking curve (0–100%) was divided into segments, with finer sampling applied to regions exhibiting rapid *R*^2^ change. This focuses evaluation on informative curve regions without sacrificing accuracy.

### 4.8 Nanobody binding validation (methods)

We tested our methods on a curated nanobody dataset targeting the SARS-CoV-2 spike protein [33]. After removing duplicates and negative controls, the final set included 3,127 unique sequences with experimentally measured binding. The goal was to evaluate whether sequence-derived importance rankings enrich for residues that mediate antigen recognition and binding energetics.

For structural validation we generated nanobody–spike complex models using ClusPro 2.0 [20]. Rigid-body docking produced an ensemble of candidate poses; representative models (including the top-ranked pose) were inspected and analysed. Full docking settings, pose selection rules and energy scoring thresholds are provided in the supplementary S5.

We used three orthogonal metrics to compare sequence-based rankings with structural evidence. First, interface residues were defined geometrically as nanobody residues with any heavy-atom within 10 Å of the antigen in the chosen docking pose. Second, for each method we computed overlap and enrichment of the top-ranked residues (for example, the top 10%) against the docking-derived interface set; discovery curves and enrichment statistics are reported in Figures 7e and 7f. Third, we decomposed per-residue interaction energies (van der Waals and electrostatic terms) for the most probable pose and examined correlations between energetic contributions, computed ΔG from in-silico point mutations, and sequence-based importance scores.

Together, these analyses provide an orthogonal structural check: geometric proximity, overlap/enrichment, and energetic sensitivity test different aspects of whether sequence-based methods identify residues that are both spatially and energetically relevant for binding.

## 6 Software Availability

All code, datasets, multiple sequence alignments, and trained models are publicly available at our GitHub repository [https://github.com/hs280/NaiveEntropicMeasures]. We provide two complementary interfaces for accessing our pipeline:

**Graphical user interface (GUI):** A lightweight HTML5-based web interface provides interactive control of all pipeline parameters without requiring coding experience. Users can run the interface locally in any web browser by executing a single command. The GUI enables: (1) specification of input alignments, target properties, and output directories; (2) selection of entropy measures and encoding schemes; (3) configuration of pipeline analysis stages; (4) real-time visualization of residue importance profiles and rankings; (5) interactive browsing and inspection of computed PDB structures with per-residue coloration reflecting importance scores; and (6) export of results to multiple formats. The interface includes live logging of pipeline execution, progress tracking, and integrated plot viewers for exploratory analysis. Both algorithmic choices (supervised vs. unsupervised, encoding type, decoupling method, information-theoretic measure) and computational parameters are exposed to users, enabling customization while providing guided defaults for non-expert practitioners.

**Command-line interface (CLI):** For advanced users and high-throughput batch processing, all functionality is available through a modular Python command-line pipeline with full argument support and sensible defaults. Running the entry point with no arguments automatically launches the HTML GUI; providing arguments enables direct CLI execution. Full documentation and usage examples are provided in the repository.

## 7 Availability of Data and Materials

All datasets, multiple sequence alignments, and code are publicly available at our GitHub repository [https://github.com/hs280/NaiveEntropicMeasures].

## Acknowledgements

W.E.H. thanks EPSRC (EP/M002403/1 and EP/N009746/1) for financial support.

## References

[1] Singh, N. et al. A generalized platform for artificial intelligence-powered autonomous enzyme engineering. Nature Communications 16 (2025). URL 10.1038/s41467-025-61209-y.

[2] Warshel, A. Electrostatic origin of the catalytic power of enzymes and the role of preorganized active sites. Journal of Biological Chemistry 276, 33235–33239 (2001).

[3] Agarwal, P. Role of protein dynamics in reaction rate enhancement by enzymes. Journal of the American Chemical Society 124, 1525–1531 (2002).

[4] Ranganathan, R. et al. Evolutionary trace method defines binding surfaces common to protein families. Journal of Molecular Biology 257, 841–858 (1996).

[5] Arnold, F. Combinatorial and computational challenges for biocatalyst design. Nature 409, 253–257 (2001).

[6] Strohl, W. Therapeutic Antibody Engineering: Current and Future Advances Driving the Strongest Growth Area in the Pharmaceutical Industry (Woodhead Publishing, 2015).

[7] Pavelka, A. et al. Hotspot wizard: a web server for identification of hot spots in protein engineering. Nucleic Acids Research 37, W376–W383 (2009).

[8] Capriotti, E. & Fariselli, P. Phd-snpg: a webserver and lightweight tool for scoring single nucleotide variants. Nucleic Acids Research 45, W247–W252 (2017).

[9] Pupko, T., Bell, R. E., Mayrose, I., Glaser, F. & Ben-Tal, N. Rate4site: an algorithmic tool for the identification of functional regions in proteins by surface mapping of evolutionary determinants within their homologues. Bioinformatics 18, S71–S77 (2002). URL 10.1093/bioinformatics/18.suppl_1.S71.

[10] Hopf, T. et al. Mutation effects predicted from sequence co-variation. Nature Biotechnology 35, 128–135 (2017).

[11] Riesselman, A. et al. Deep generative models of genetic variation capture the effects of mutations. Nature Methods 15, 816–822 (2018).

[12] Feizi, S., Marbach, D., Médard, M. & Kellis, M. Network deconvolution as a general method to distinguish direct dependencies in networks. Nature Biotechnology 31, 726–733 (2013). URL https://www.nature.com/articles/nbt.2635.

[13] Romero, E. et al. Same substrate, many reactions: Oxygen activation in flavoenzymes. Chem Rev 118, 1742–1769 (2018).

[14] Fpbase. https://www.fpbase.org/.

[15] Heim, R., Prasher, D. C. & Tsien, R. Y. Wavelength mutations and posttranslational autoxidation of green fluorescent protein. Proceedings of the National Academy of Sciences 91, 12501–12504 (1994).

[16] Tsien, R. The green fluorescent protein. Annual Review of Biochemistry 67, 509–544 (1998).

[17] Ormö, M., et al. Crystal structure of the aequorea victoria green fluorescent protein. Science 273, 1392–1395 (1996).

[18] Pakhomov, A. A. & Martynov, V. I. Gfp family: Structural insights into spectral tuning. Chemistry & Biology 15, 755–764 (2008).

[19] Karasuyama, M., Inoue, K., Nakamura, R., Kandori, H. & Takeuchi, I. Understanding colour tuning rules and predicting absorption wavelengths of microbial rhodopsins by data-driven machinelearning approach. Scientific Reports 8, 16334 (2018).

[20] Kozakov, D. et al. The cluspro web server for protein–protein docking. Nature Protocols 12, 255–278 (2017).

[21] Deep learning prediction of enzyme optimum ph. bioRxiv (2021). URL https://www.biorxiv.org/content/10.1101/2021.06.25.449983v1.

[22] Uniprot. https://www.uniprot.org/.

[23] Ncbi. https://www.ncbi.nlm.nih.gov/.

[24] Huang, S. et al. Identification and characterization of a catalytic base in bacterial luciferase by chemical rescue of a dark mutant. Biochemistry 37, 8614–8614 (1998).

[25] Campbell, Z. et al. Crystal structure of the bacterial luciferase/flavin complex provides insight into the function of the beta subunit. Biochemistry 48, 6085–6094 (2009).

[26] Li, C. et al. Active site hydrophobicity is critical to the bioluminescence activity of vibrio harveyi luciferase. Biochemistry 44, 12970–12977 (2005).

[27] Low, J. et al. Functional roles of conserved residues in the unstructured loop of vibrio harveyi bacterial luciferase. Biochemistry 41, 1724–1731 (2002).

[28] Li, H. et al. Effects of mutations of the his45 residue of vibrio harveyi luciferase on the yield and reactivity of the flavin peroxide intermediate. Biochemistry 38, 4409–4415 (1999).

[29] Tinikul, R. et al. Structure, mechanism, and mutation of bacterial luciferase. Adv Biochem Eng Biotechnol 154, 47–74 (2016).

[30] Eckstein, J. et al. Mechanism of bacterial bioluminescence: 4a,5-dihydroflavin analogs as models for luciferase hydroperoxide intermediates and the effect of substituents at the 8-position of flavin on luciferase kinetics. Biochemistry 32, 404–411 (1993).

[31] Gado, J. E. et al. Machine learning prediction of enzyme optimum ph. Nature Machine Intelligence 7, 716–729 (2025). URL 10.1038/s42256-025-01026-6.

[32] Chang, A. et al. Brenda, the elixir core data resource in 2021: new developments and updates. Nucleic Acids Research 49, D498–D508 (2020). URL 10.1093/nar/gkaa1025. https://academic.oup.com/nar/article-pdf/49/D1/D498/35364686/gkaa1025.pdf.

[33] Engelhart, E. et al. A dataset comprised of binding interactions for 104,972 antibodies against a sars-cov-2 peptide. Scientific Data 9, 653 (2022).

[34] Lassmann, T. Kalign 3: multiple sequence alignment of large datasets. Bioinformatics 36, 1928–1929 (2019).

[35] Lin, Z. et al. Evolutionary-scale prediction of atomic-level protein structure with a language model. Science 379, 1123–1130 (2023). URL https://www.science.org/doi/abs/10.1126/science.ade2574. https://www.science.org/doi/pdf/10.1126/science.ade2574.

[36] Tsujimura, M. & Ishikita, H. Insights into the protein functions and absorption wavelengths of microbial rhodopsins. Journal of Physical Chemistry B 124, 12956–12965 (2020). URL https://pubs.acs.org/doi/10.1021/acs.jpcb.0c08910.

