## Supplementary material for "Disentangling Protein Function via Decoupled Information Theoretic Selection of Key Tuning Residues": S1: Supplementary Figures 1

October 6, 2025

[1]Department of Engineering Science, University of Oxford, Oxford, OX1 3PJ, United Kingdom

*Corresponding author:

#### Overview

This supplementary document provides a comprehensive visual analysis of three diverse protein families: Alkane Monooxygenases, Rhodopsins, and Fluorescent Proteins. These analyses encompass sequence characteristics, evolutionary relationships, conservation patterns, and inter-family structural comparisons. All visualizations were generated using custom analysis pipelines designed for publication-quality Nature-style graphics.

### 1 Sequence Length and Composition Analysis

#### 1.1 Alkane Monooxygenase Family

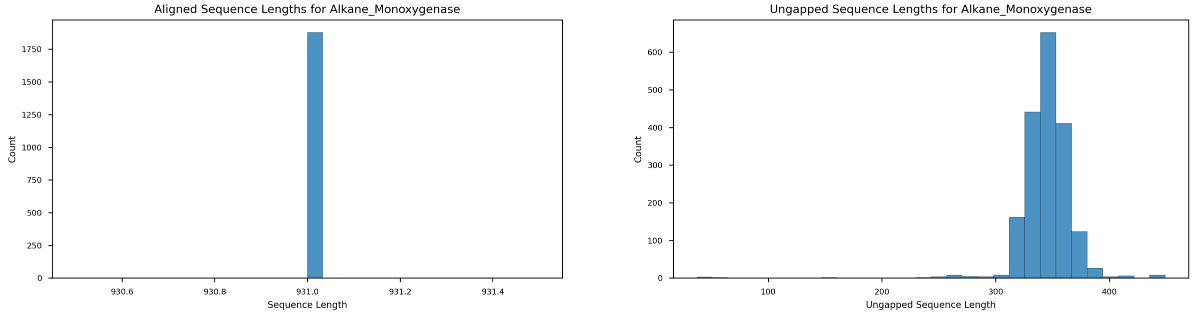


(a) (b)

Figure 1: **Sequence length distributions for the Alkane Monooxygenase family.** **(a)** Distribution of aligned sequence lengths including gaps, showing the post-alignment length distribution across 1,880 sequences. The tight distribution indicates successful alignment with minimal variability in total aligned length. **(b)** Distribution of ungapped sequence lengths, revealing the true variation in protein size after removing alignment-introduced gaps. This represents the actual coding sequence diversity within the family, with most sequences clustering around 350-400 amino acids, consistent with the conserved catalytic core of bacterial luciferases and related monooxygenases.

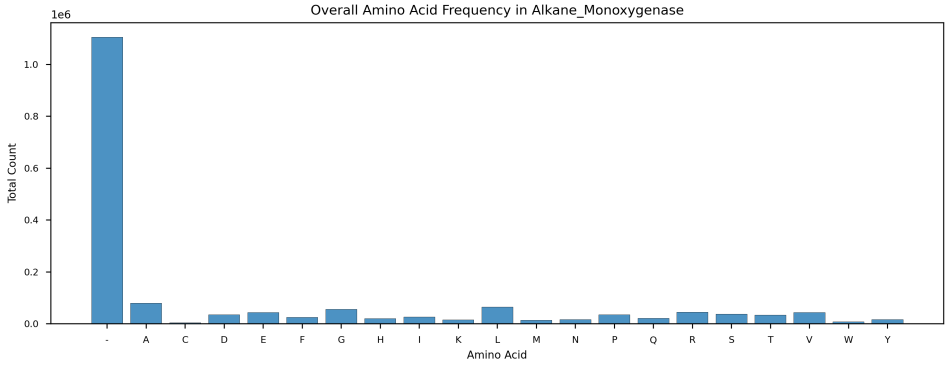


Figure 2: **Overall amino acid composition of the Alkane Monooxygenase family.** Aggregate frequency distribution of all 20 standard amino acids across the entire multiple sequence alignment (1,880 sequences). The distribution reflects the functional requirements of these enzymes: high leucine (L) and alanine (A) content supports the hydrophobic active site environment critical for substrate binding and catalysis; elevated glycine (G) frequency provides structural flexibility in loop regions; and enrichment of charged residues (E, D, K, R) facilitates electrostatic stabilization and potential pH-dependent regulation of enzymatic activity.

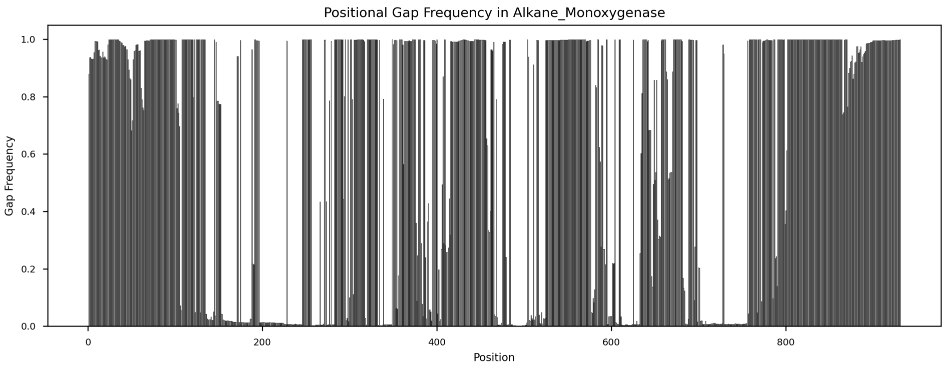


Figure 3: **Positional gap frequency across the Alkane Monooxygenase alignment.** This plot reveals the distribution of alignment gaps at each position, with higher values indicating regions of insertion/deletion events or sequence divergence. Regions with low gap frequency (¡0.1) represent structurally conserved core domains, likely corresponding to the catalytic machinery and essential structural elements. Peaks of high gap frequency (¿0.3) indicate variable loop regions, potential insertion domains, or lineage-specific adaptations. These variable regions may contribute to substrate specificity differences or regulatory mechanisms across different monooxygenase subfamilies.

#### 1.2 Rhodopsin Family

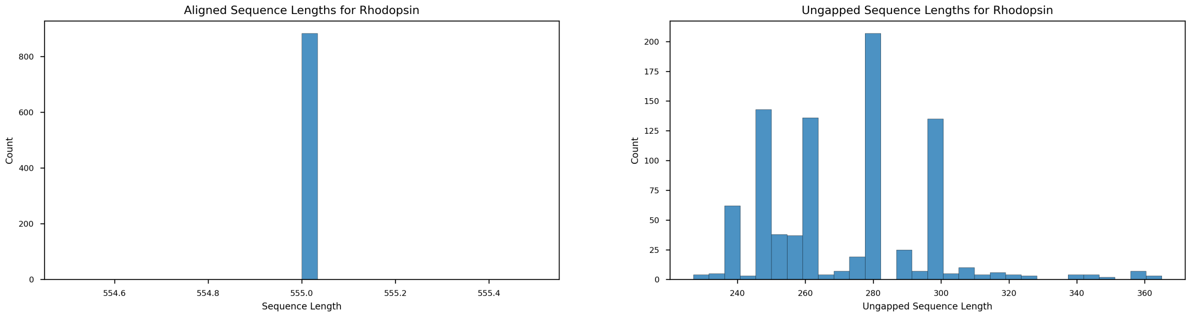


(a) (b)

Figure 4: **Sequence length distributions for the Rhodopsin family.** **(a)** Distribution of aligned sequence lengths for 884 microbial rhodopsin sequences, showing a uniform aligned length characteristic of highly conserved seven-transmembrane helix architecture. **(b)** Ungapped sequence length distribution revealing remarkably low variability, with most sequences clustering tightly around 250 amino acids. This conservation reflects the structural constraints of the canonical 7-TM fold required for retinal binding and proton/ion pumping function. The minimal length variation suggests strong purifying selection maintaining the essential transmembrane architecture across diverse rhodopsin subtypes.

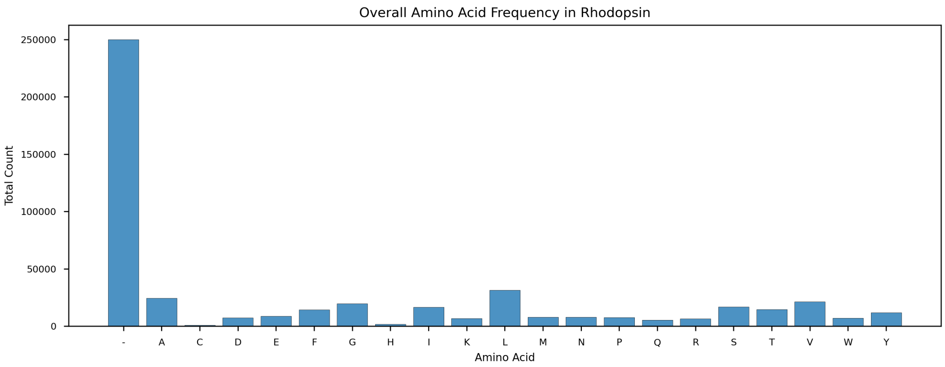


Figure 5: **Overall amino acid composition of the Rhodopsin family.** The amino acid distribution reflects the membrane-embedded nature of rhodopsins: dominant hydrophobic residues (L, A, V, I, F) are consistent with extensive transmembrane helical content; elevated glycine facilitates tight helix packing and flexibility in the retinal-binding pocket; enrichment of charged residues (D, E, K, R) at specific positions forms the proton transfer pathway and counterion complex essential for chromophore stabilization and photocycle function. The relatively high abundance of aromatic residues (F, Y, W) supports both structural stability through - interactions and spectral tuning through interactions with the retinal chromophore.

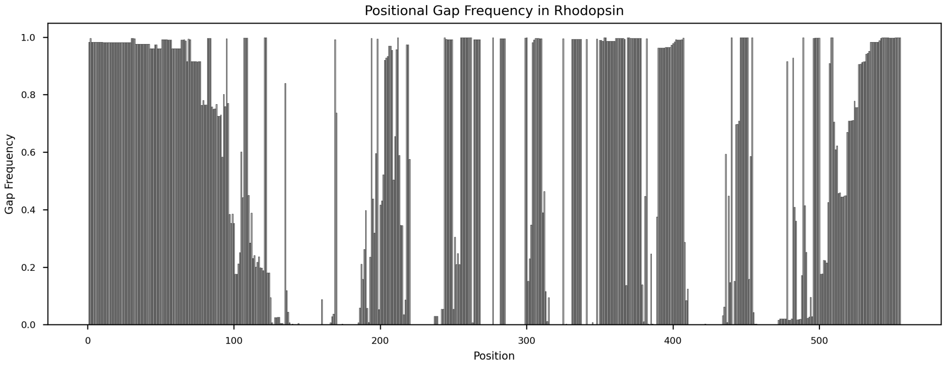


Figure 6: **Positional gap frequency across the Rhodopsin alignment.** The gap distribution reveals the structural organization of microbial rhodopsins. Extended regions of near-zero gap frequency correspond to the seven conserved transmembrane helices (A-G), which maintain their length and register across all family members. Localized peaks of elevated gap frequency mark the connecting loops between helices, particularly the extracellular and cytoplasmic loops, which show greater sequence length variability. The low overall gap frequency (¡0.15 at most positions) underscores the exceptional structural conservation required for maintaining the retinal-binding pocket geometry and proton pumping mechanism across diverse rhodopsin subfamilies.

#### 1.3 Fluorescent Protein Family

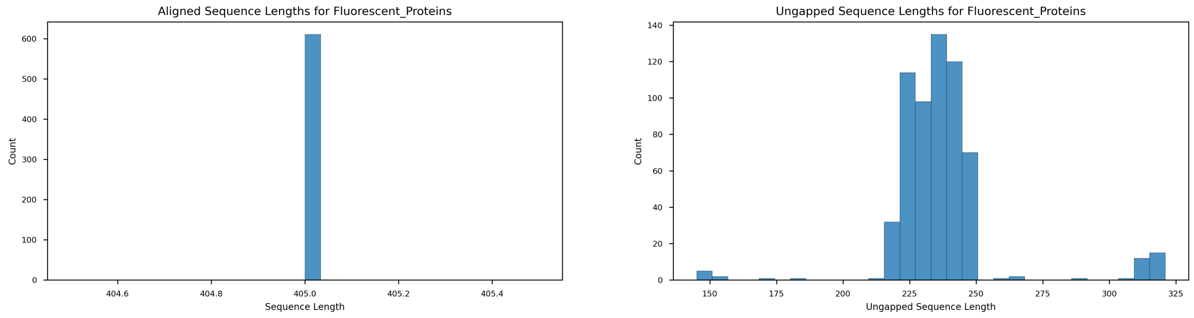


(a) (b)

Figure 7: **Sequence length distributions for the Fluorescent Protein family.** **(a)** Distribution of aligned sequence lengths for 611 GFP-like fluorescent proteins, showing consistent alignment length with minimal variability. **(b)** Ungapped sequence length distribution demonstrating tight clustering around 225-240 amino acids, reflecting the conserved -barrel architecture characteristic of the GFP superfamily. This length conservation is expected given that the 11-stranded -barrel structure and central -helix must be maintained to properly form and protect the autocatalytically generated chromophore. Slight variations likely represent N- or C-terminal extensions that do not disrupt the core barrel structure but may modulate folding kinetics, oligomerization, or photophysical properties.

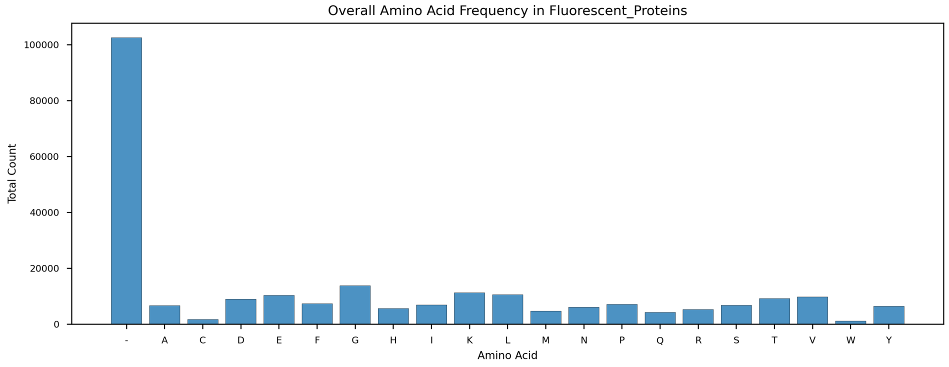


Figure 8: **Overall amino acid composition of the Fluorescent Protein family.** The amino acid distribution reveals features characteristic of -barrel proteins: high glycine content facilitates the tight turns required in the barrel structure; abundant leucine, alanine, and valine form the hydrophobic core that shields the chromophore from solvent quenching; elevated serine and threonine frequencies support the extensive hydrogen bonding network within the barrel that maintains structural rigidity and chromophore planarity. The enrichment of aromatic residues (Y, F) is functionally significant, as tyrosine-65 (in GFP numbering) is typically part of the chromophore-forming triad, and surrounding aromatics modulate fluorescence through -stacking interactions and excited-state stabilization.

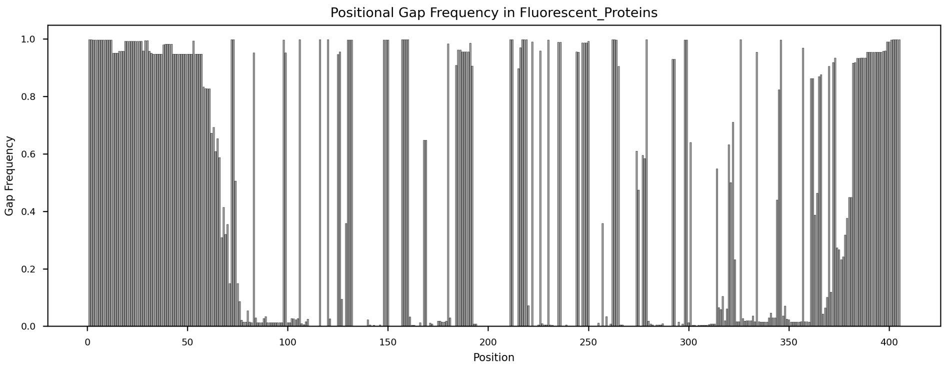


Figure 9: **Positional gap frequency across the Fluorescent Protein alignment.** The gap distribution pattern reflects the modular architecture of fluorescent proteins. Extended stretches of low gap frequency (¡0.05) correspond to the 11 -strands that form the conserved barrel framework. These regions show exceptional length conservation, as alterations would disrupt the barrel geometry and chromophore environment. Localized peaks of higher gap frequency mark the connecting loops between -strands, particularly surface-exposed loops that tolerate insertions/deletions without compromising core structure. The overall low gap content indicates that most evolutionary variation occurs through substitution rather than indel events, preserving the precise barrel architecture required for chromophore formation and optical properties.

### 2 Sequence Divergence and Homology Analysis

#### 2.1 Pairwise Distance Distributions

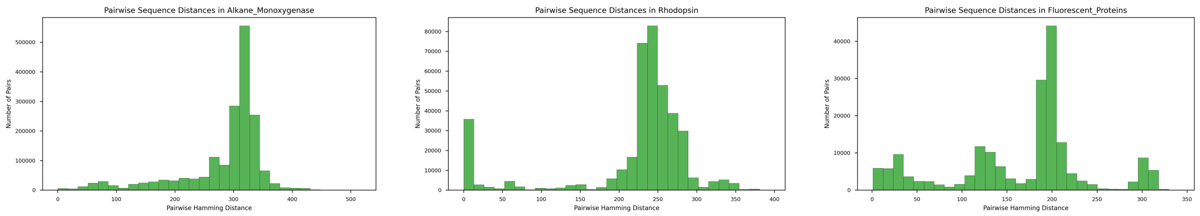


(a) Alkane Monooxygenase

(b) Rhodopsin

(c) Fluorescent Proteins

Figure 10: **Pairwise Hamming distance distributions across protein families.** These histograms quantify sequence divergence by plotting the distribution of pairwise Hamming distances (number of differing positions) between all sequence pairs within each family. **(a)** Alkane Monooxygenases show a broad distribution (mean 150-200 differences), indicating substantial sequence diversity across the family, consistent with adaptation to different substrates and environmental conditions. **(b)** Rhodopsins display an intermediate distribution, reflecting diversification across different proton/ion pumping mechanisms and spectral tuning adaptations. **(c)** Fluorescent Proteins show the tightest distribution, suggesting recent common ancestry and conservation of the core barrel structure, with diversity primarily reflecting spectral variants rather than deep evolutionary divergence.

#### 2.2 Theoretically Scaled Distance Distributions

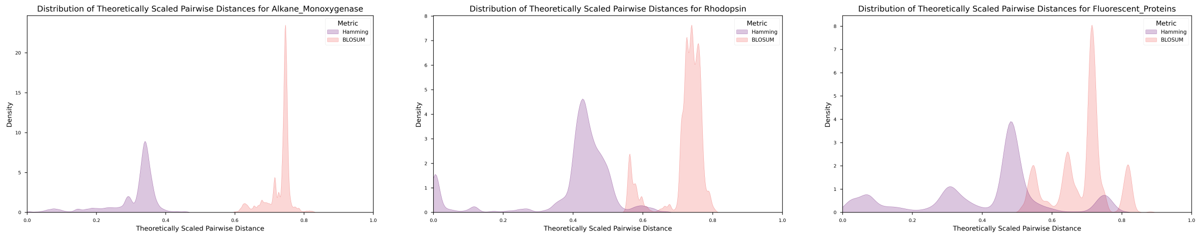


(a) Alkane Monooxygenase (b) Rhodopsin (c) Fluorescent Proteins

Figure 11: **Theoretically scaled distance distributions comparing Hamming and BLOSUM metrics.** These kernel density estimates compare two distance metrics normalized to [0,1]: simple Hamming distance (position-level identity) and BLOSUM62-based biochemical distance (accounting for amino acid similarity). The relationship between these curves reveals the nature of sequence variation. When BLOSUM distances are systematically lower than Hamming distances, it indicates that substitutions are predominantly conservative (biochemically similar residues), suggesting functional constraint. Conversely, similar distributions suggest non-conservative changes or relaxed selection. **(a)** Alkane Monooxygenases show substantial separation between metrics, indicating many conservative substitutions in substrate-binding regions. **(b)** Rhodopsins display moderate separation, consistent with conservation of hydrophobic character in transmembrane regions while allowing spectral tuning substitutions. **(c)** Fluorescent Proteins show the smallest metric separation, suggesting tighter structural constraints.

### 3 Conservation and Variability Analysis

#### 3.1 Positional Entropy Profiles

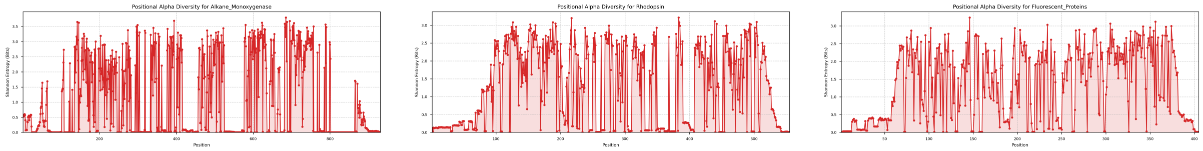

(a) Alkane Monooxygenase (b) Rhodopsin (c) Fluorescent Proteins

Figure 12: **Shannon entropy profiles revealing conservation patterns across alignment positions.** Entropy (bits) quantifies amino acid variability at each position: low entropy (¡1 bit) indicates strong conservation, while high entropy (¿3 bits) indicates high variability. These profiles identify functionally critical positions (low entropy) versus those tolerating substitution. **(a)** Alkane Monooxygenases show heterogeneous entropy with distinct low-entropy peaks marking the active site residues (e.g., His44, His45, Cys106) and flavin-binding motifs, interspersed with high-entropy regions in surface-exposed loops. **(b)** Rhodopsins display characteristic periodic patterns corresponding to transmembrane helices (low entropy) and connecting loops (higher entropy), with exceptionally conserved positions marking retinal-binding lysine and proton transfer pathway residues. **(c)** Fluorescent Proteins show lower overall entropy, with minimal variation in barrel-forming strands and elevated entropy only in surface loops, reflecting the stringent requirements for chromophore formation and barrel integrity.

#### 3.2 Residue Probability Heatmaps

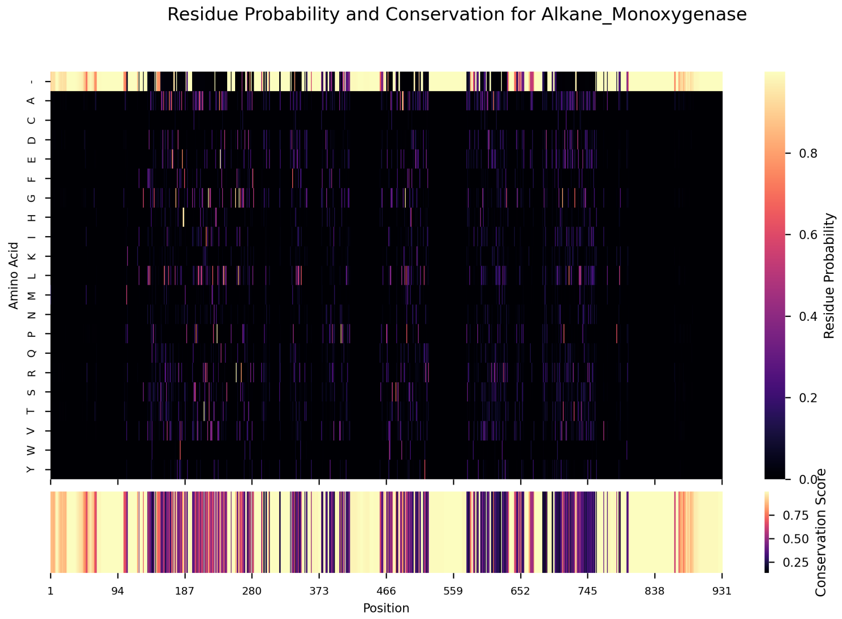


Figure 13: **Residue probability and conservation heatmap for Alkane Monooxygenases.** The upper panel displays the probability of each amino acid (y-axis) at each alignment position (x-axis), with color intensity representing frequency. Vertical ”hot streaks” indicate highly conserved positions dominated by a single residue type, while diffuse columns indicate variable positions. The lower panel shows position-wise conservation scores (inverse of Shannon entropy), where bright regions mark functionally critical, invariant positions. This visualization reveals: (1) absolutely conserved catalytic residues appearing as bright horizontal bands at specific positions; (2) hydrophobic patches corresponding to structural core; (3) variable surface-exposed regions showing mixed residue profiles. The conservation track identifies candidate positions for mutagenesis studies, as highly conserved positions likely play essential catalytic or structural roles.

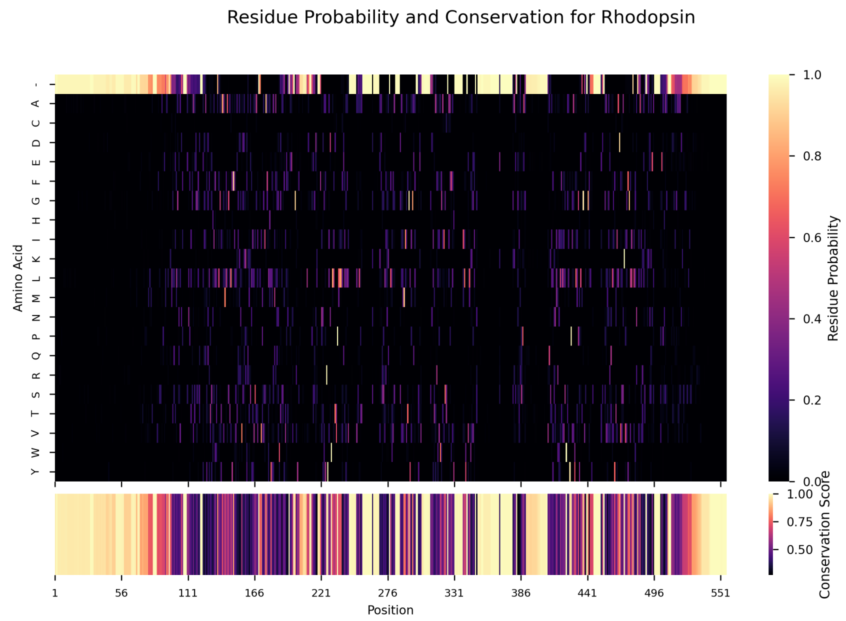


Figure 14: **Residue probability and conservation heatmap for Rhodopsins.** This dual-panel visualization captures the exquisite conservation pattern of microbial rhodopsins. The upper probability heatmap reveals position-specific amino acid preferences: extended stretches dominated by hydrophobic residues (L, V, I, F) mark the seven transmembrane helices; periodic appearance of charged residues (D, E, K, R) at conserved positions identifies the proton transfer pathway; absolute conservation of specific positions (bright vertical bands) marks functionally critical sites including the retinal-binding lysine (Schiff base), the counterion aspartate/glutamate, and aromatic residues involved in spectral tuning. The conservation track (lower panel) shows characteristic high conservation within helices and reduced conservation in loops, consistent with the structural and functional constraints of the rhodopsin fold.

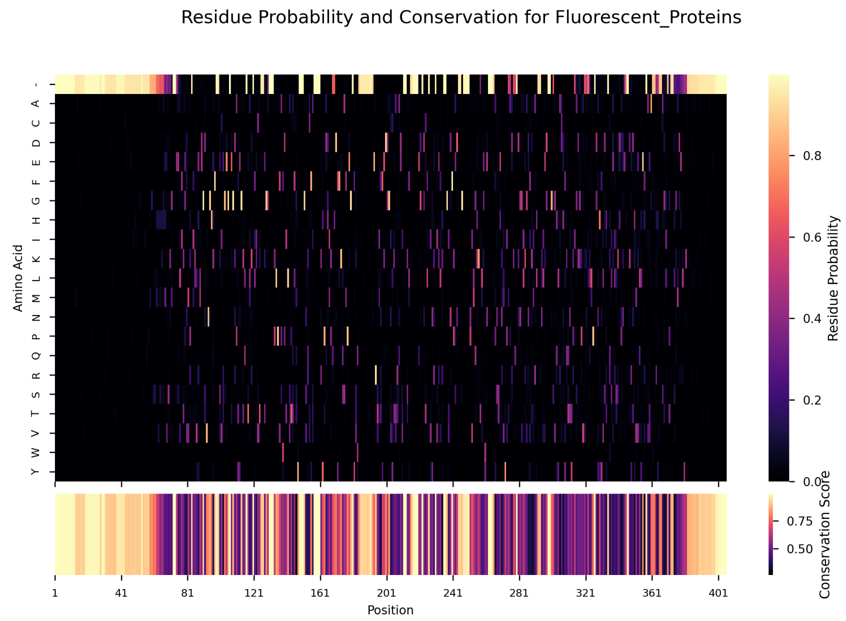


Figure 15: **Residue probability and conservation heatmap for Fluorescent Proteins.** The heatmap illustrates the remarkable conservation of the GFP -barrel architecture. The upper panel shows strong position-specific preferences: glycine-rich regions facilitating tight turns between -strands; alternating patterns of hydrophobic residues forming the barrel interior; absolute conservation at the chromophore-forming triad positions (typically Ser65-Tyr66-Gly67 in GFP numbering), visible as three adjacent bright vertical streaks. The conservation track (lower panel) reveals exceptionally high conservation throughout the barrel-forming regions (¿90% of positions), with only surface-exposed loops showing moderate variability. This pattern reflects the dual requirements of maintaining barrel geometry for chromophore protection while preserving the precise chemical environment for autocatalytic chromophore formation and spectral properties.

### 4 Internal Homology and Distance Matrices

#### 4.1 Hamming Distance-Based Homology

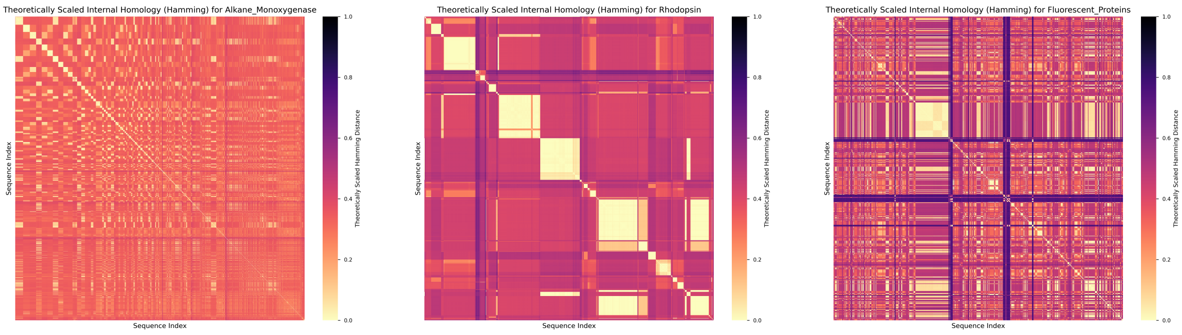


(a) Alkane Monooxygenase (b) Rhodopsin (c) Fluorescent Proteins

Figure 16: **Theoretically scaled Hamming distance matrices revealing internal homology structure.** These heatmaps display all-vs-all pairwise Hamming distances (normalized to [0,1]) for each family, with darker colors indicating higher similarity. Block-like patterns reveal subfamily structure: distinct dark squares along the diagonal indicate tightly related sequence clusters (subfamilies), while off-diagonal patterns quantify inter-cluster relationships. **(a)** Alkane Monooxygenases show clear modular structure with multiple distinct sequence clusters, likely corresponding to different bacterial lineages or substrate specificities. **(b)** Rhodopsins display intermediate clustering, reflecting the major functional classes (proton pumps, chloride pumps, sensory rhodopsins). **(c)** Fluorescent Proteins show the most homogeneous pattern, consistent with a recently diversified family primarily differing in spectral properties rather than deep phylogenetic splits.

#### 4.2 BLOSUM62 Distance-Based Homology

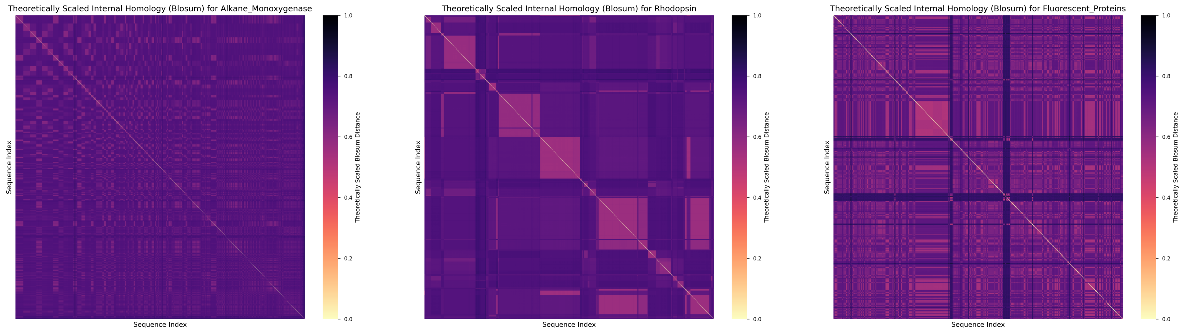


(a) Alkane Monooxygenase (b) Rhodopsin (c) Fluorescent Proteins

Figure 17: **Theoretically scaled BLOSUM62 distance matrices capturing biochemical similarity.** Unlike Hamming distance, BLOSUM62 scoring accounts for biochemical similarity between amino acids, treating conservative substitutions as more similar than radical changes. Comparing these matrices to Hamming-based matrices (Figure 16) reveals whether sequence divergence occurs primarily through conservative or non-conservative substitutions. Generally darker BLOSUM matrices compared to Hamming indicate prevalent conservative substitutions, suggesting functional constraint. **(a)** Alkane Monooxygenases show noticeably darker BLOSUM patterns, indicating many conservative substitutions, particularly in the substrate-binding pocket where hydrophobic character is maintained. **(b)** Rhodopsins display similar darkening in transmembrane regions, consistent with conservation of membrane-spanning hydrophobic character. **(c)** Fluorescent Proteins show less difference between metrics, reflecting tighter overall constraint with fewer permissible substitutions.

### 5 Phylogenetic Relationships and Network Analysis

#### 5.1 Minimum Spanning Trees (Hamming Distance)

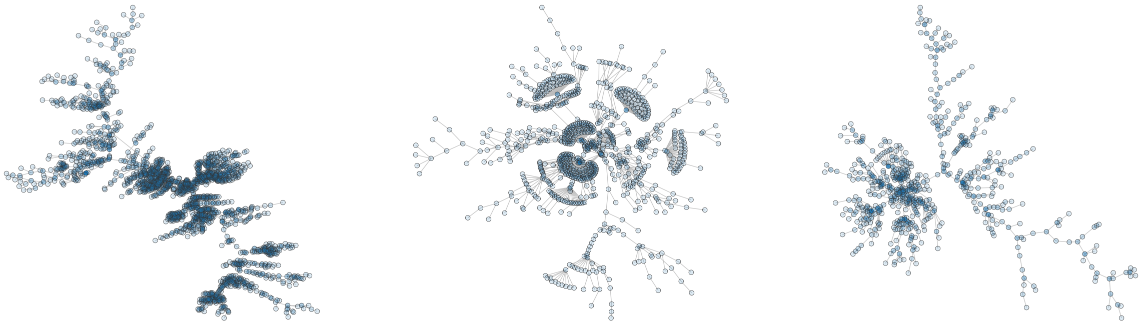

(a) Alkane Monooxygenase (b) Rhodopsin (c) Fluorescent Proteins

Figure 18: **Minimum spanning tree representations based on Hamming distances.** These network visualizations connect sequences via the minimum total distance (Hamming-based), revealing evolutionary relationships and subfamily structure without imposing a bifurcating tree topology. Node opacity reflects network degree (hub sequences with many close neighbors appear darker), identifying ancestral or central sequences. Edge lengths qualitatively represent genetic distance. The overall topology reveals diversification patterns: star-like topologies suggest recent radiation from a common ancestor, while branched structures indicate older, more complex evolutionary histories. **(a)** Alkane Monooxygenases display multiple discrete clusters with hub sequences, consistent with independent diversification in different bacterial lineages. **(b)** Rhodopsins show a more continuous distribution with major hubs, reflecting the diversification of major functional classes. **(c)** Fluorescent Proteins exhibit a more compact structure with less extreme branching, consistent with recent diversification primarily driven by laboratory engineering and natural color variation.

#### 5.2 Minimum Spanning Trees (BLOSUM62 Distance)

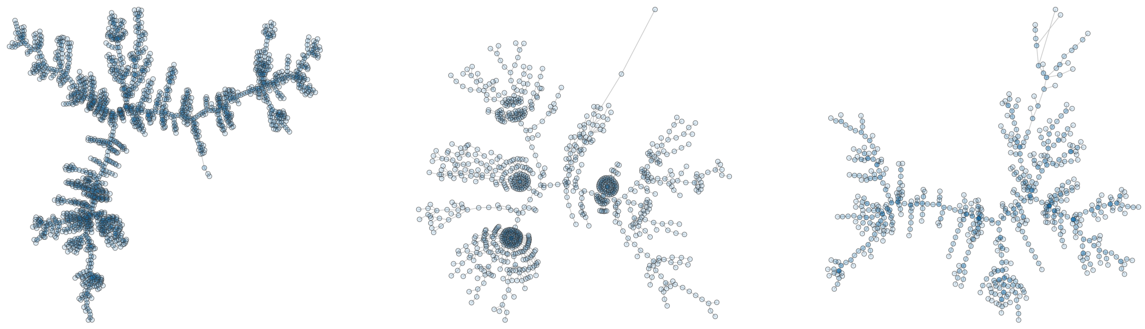

(a) Alkane Monooxygenase (b) Rhodopsin (c) Fluorescent Proteins

Figure 19: **Minimum spanning tree representations based on BLOSUM62 distances.** These MSTs use biochemical similarity rather than strict identity for edge weighting. Comparing to Hamming-based MSTs (Figure 18) reveals how accounting for substitution similarity affects inferred relationships. More compact BLOSUM-based trees suggest that apparent sequence diversity is partly due to conservative substitutions that maintain biochemical properties. Topological changes between Hamming and BLOSUM trees identify sequences differing primarily through conservative vs. radical substitutions. **(a)** Alkane Monooxygenases show somewhat tighter clustering in the BLOSUM tree, indicating functional constraint through biochemical conservation. **(b)** Rhodopsins display similar topology changes, particularly in transmembrane-helix-rich regions. **(c)** Fluorescent Proteins show minimal topological change, consistent with tight structural constraints allowing few substitutions of any type.

### 6 Cross-Family Comparative Analysis

#### 6.1 Cross-Family Minimum Spanning Trees

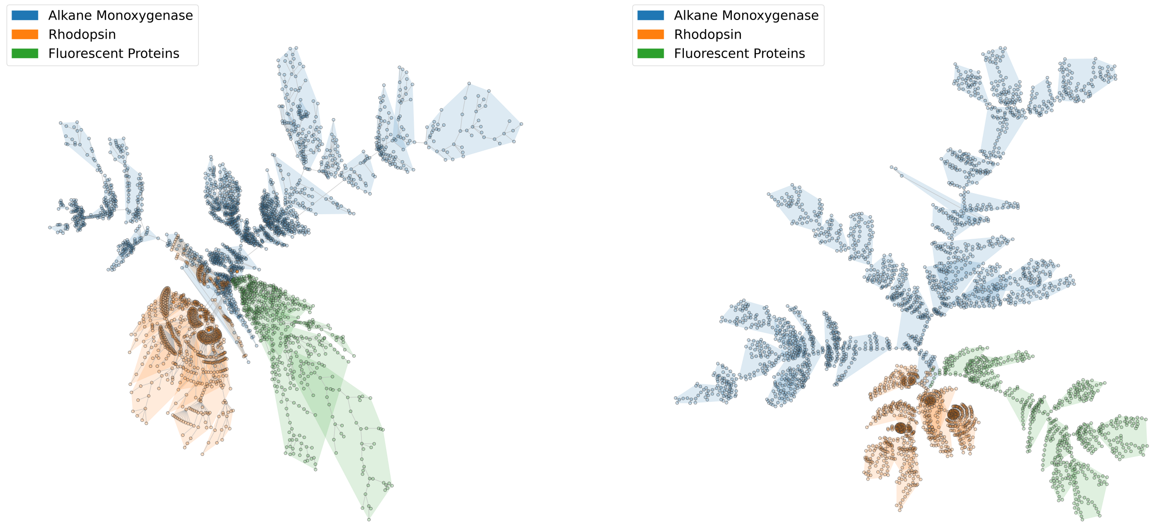


(a) Hamming distance

(b) BLOSUM62 distance

Figure 20: **Cross-family minimum spanning trees with community detection.** These unified MSTs combine all three protein families into a single network, with nodes colored by family membership (Alkane Monooxygenases, Rhodopsins, Fluorescent Proteins). The visualization tests whether sequence-based distances naturally cluster sequences by known family boundaries. Perfect family separation would show three distinct, non-overlapping color clusters; mixing would indicate convergent sequence features or alignment artifacts. Both distance metrics show excellent family separation, validating that: (1) the alignment captures family-specific signatures; (2) these families are evolutionarily distinct with minimal sequence-level convergence; (3) within-family variation is substantially less than between-family divergence. **(a)** Hamming-based MST shows clean separation of all three families with minimal inter-family edges, demonstrating strong phylogenetic signal. **(b)** BLOSUM-based MST maintains similar topology, indicating that family boundaries are robust to whether strict identity or biochemical similarity is used for comparison. The relative positioning of family clusters reflects their overall sequence divergence: Fluorescent Proteins and Rhodopsins (both having -sheet or -helical regularity) may show closer proximity than either to the more sequence-diverse Alkane Monooxygenases.

### Conclusions

This comprehensive visual analysis reveals distinct signatures for each protein family:

**Alkane Monooxygenases** exhibit substantial sequence diversity with clear subfamily structure, reflecting adaptation to diverse substrates and bacterial lineages. The high sequence variability outside conserved catalytic residues suggests this family as an excellent target for protein engineering, with multiple ”tunable” positions that can be modified without disrupting catalytic function.

**Rhodopsins** display exceptional conservation of core transmembrane architecture with localized variability in spectral-tuning and ion-specificity determinants. The periodic conservation pattern (helices vs. loops) provides a clear map for rational mutagenesis studies targeting wavelength shifts or pumping efficiency without compromising the fundamental 7-TM fold.

**Fluorescent Proteins** show the tightest conservation, particularly in barrel-forming regions, consistent with stringent requirements for chromophore formation. The limited sequence diversity outside surface loops suggests that generating novel optical properties may require targeted, structure-guided mutagenesis rather than random diversification.

The cross-family comparisons validate that these families represent genuinely distinct evolutionary lineages with minimal sequence-level convergence, making them ideal test cases for methods development in protein function prediction and design.
