## Supplementary material for "Disentangling Protein Function via Decoupled Information Theoretic Selection of Key Tuning Residues": S2: Supplementary Figures 2

October 23, 2025

[1]Department of Engineering Science, University of Oxford, Oxford, OX1 3PJ, United Kingdom

*Corresponding author:

#### Overview

This supplementary document presents a comprehensive analysis of various information-theoretic measures applied to protein sequence alignments. We analyze three protein families (Alkane Monooxygenases, Rhodopsins, and Fluorescent Proteins) using both standard sequence encoding and physicochemical property-based encoding. The analysis includes 11 different entropy-based measures, mutual information calculations, coupling matrices, and extensive cross-correlation analyses to identify the most robust and informative metrics for protein engineering applications.

#### 1 Methodology Overview

Our analysis pipeline computes multiple information-theoretic measures for each alignment position:

- **Shannon Entropy**: Classical measure of amino acid variability at each position
- **Rényi Entropy**: Generalized entropy allowing emphasis on rare/common events

- **Tsallis Entropy**: Non-extensive entropy for systems with long-range correlations
- **Von Neumann Entropy**: Quantum-inspired entropy for density matrix representations
- **Kolmogorov Complexity**: Algorithmic information content approximation
- **KL Divergence**: Relative entropy from uniform distribution
- **JS Divergence**: Symmetric divergence measure
- **Gini Impurity**: Decision tree-inspired impurity measure
- **Average Entropy**: Mean across multiple entropy formulations
- **Mutual Information**: Supervised measure of position-function correlation
- **Voting Schemes**: Soft and hard voting consensus across measures

For each measure, we compute: (1) raw importance scores, (2) decoupled scores (removing evolutionary coupling effects), and (3) autocorrelation profiles to assess signal quality.

### 2 Alkane Monooxygenase Family - Standard Encoding

#### 2.1 Shannon Entropy Analysis

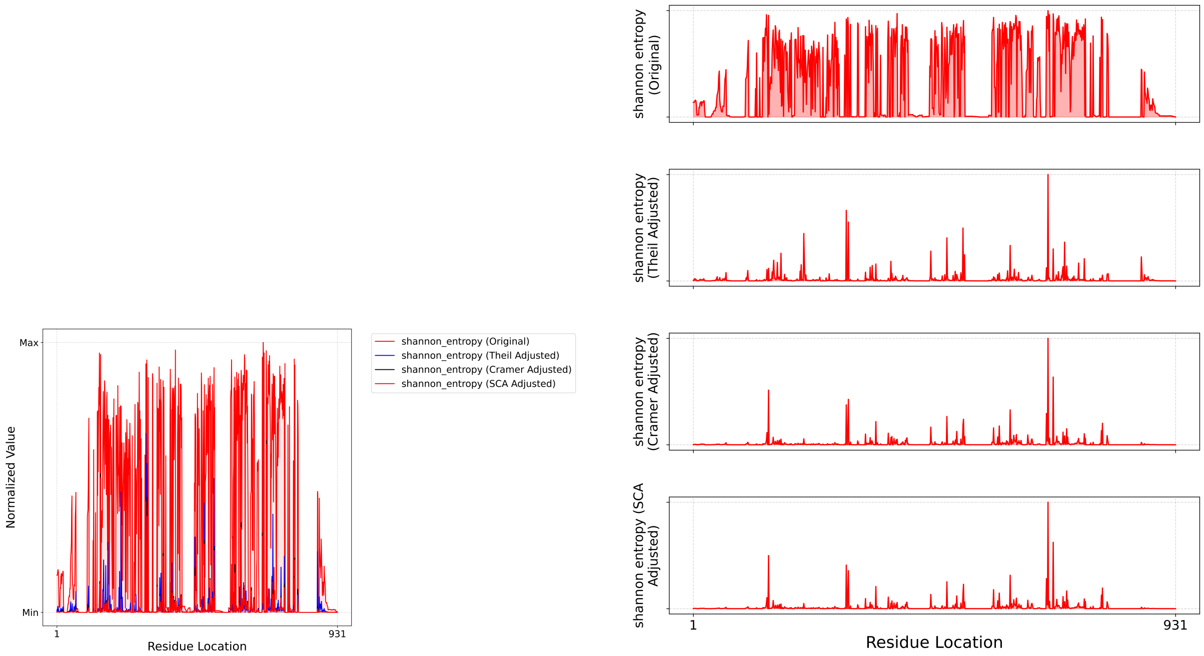


(a) Single-panel comparison

(b) Multi-panel view

Figure 1: **Shannon entropy profiles for Alkane Monooxygenases with standard encoding.** **(a)** Overlay comparison of raw Shannon entropy (red), Cramér's V decoupled entropy (blue), and Theil's U decoupled entropy (black). All traces are normalized to [0,1] for direct comparison. The raw entropy (red) shows high-frequency noise from evolutionary coupling, while decoupled traces (blue, black) reveal a sharper, more interpretable signal identifying truly informative positions. **(b)** Individual panel view showing each metric separately, highlighting how the decoupling process (both Cramér's V and Theil's U) dramatically reduces noise and isolates unique informational contributions at each position. Peaks in decoupled traces correspond to residues that are both variable and informationally independent—the key tuning residues hypothesis.

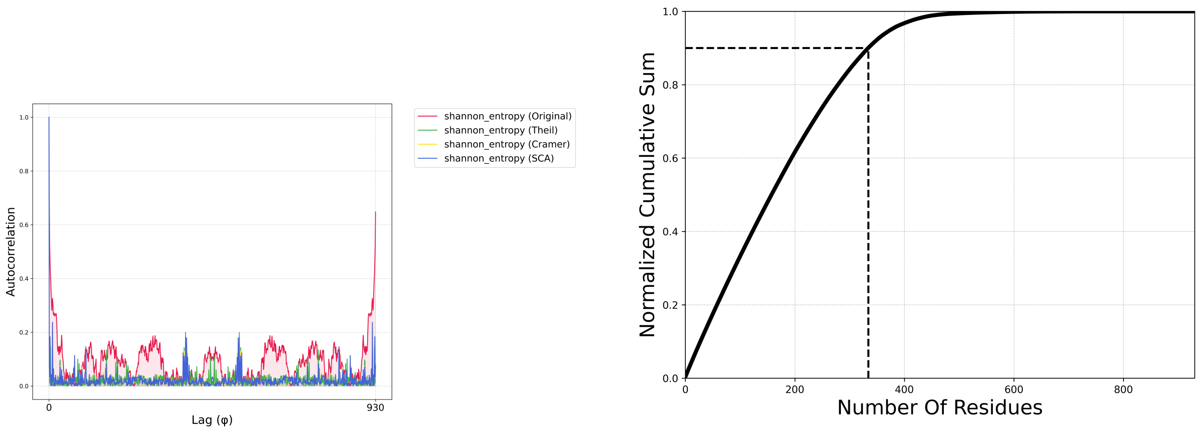


(a) Autocorrelation profiles

(b) Cumulative information content

Figure 2: **Signal quality analysis for Shannon entropy. (a)** Autocorrelation functions quantify the ”noisiness” of each metric. Raw Shannon entropy (red) shows broad, slowly decaying autocorrelation, indicating high noise and redundancy across positions—the signature of evolutionary coupling contaminating the signal. Decoupled metrics (blue, black) show sharp peaks at zero lag and rapid decay, demonstrating low noise and high information density. This transformation from noisy to clean signal is the core benefit of our decoupling approach. **(b)** Normalized cumulative sum plot reveals information concentration: 90% of the total Shannon entropy signal is captured by approximately 150 residues (marked by dashed lines), indicating that functional information is concentrated in a small subset of positions. This validates the feature selection framing and guides experimental prioritization.

#### 2.2 Mutual Information Analysis

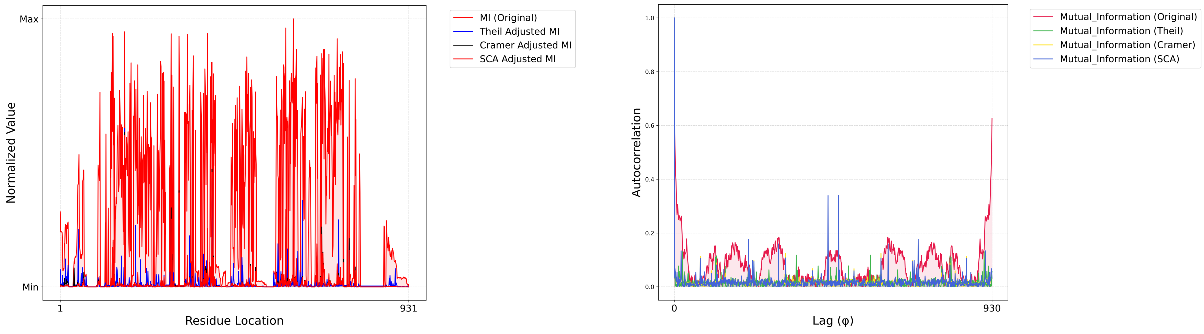


(a) MI profiles

(b) MI autocorrelation

Figure 3: **Mutual information analysis for supervised residue ranking. (a)** Mutual information directly measures the statistical dependency between amino acid identity at each position and the target functional property (pH optimum for Alkane Monooxygenases). Raw MI (red) identifies positions correlated with function but is confounded by evolutionary coupling—positions may appear important simply because they co-vary with truly functional residues. Decoupling (blue, black) isolates direct functional correlation from indirect evolutionary correlation. Peaks in decoupled MI represent positions where knowing the amino acid identity provides unique, non-redundant information about protein function. **(b)** Autocorrelation analysis confirms that decoupling dramatically reduces signal redundancy (sharp peaks) compared to raw MI (broad distribution), validating improved interpretability and reducing false positives in downstream engineering applications.

#### 2.3 Alternative Entropy Measures

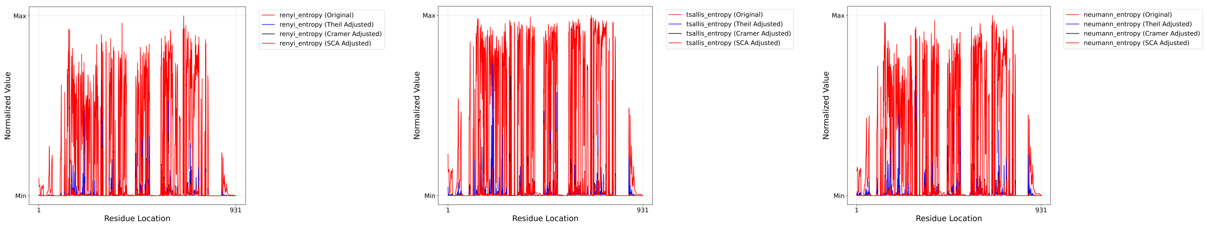


(a) Rényi Entropy (b) Tsallis Entropy (c) Von Neumann Entropy

Figure 4: **Generalized entropy measures for Alkane Monooxygenases.** These alternative entropy formulations provide different perspectives on positional variability. **(a)** Rényi entropy generalizes Shannon entropy with a tunable parameter emphasizing rare or common amino acids. Decoupling reveals that Rényi entropy (after adjustment) closely tracks Shannon entropy, suggesting robustness of the fundamental variability signal across formulations. **(b)** Tsallis entropy, designed for non-extensive systems with long-range correlations, shows qualitatively similar patterns to Shannon but with subtle differences in peak magnitudes, potentially capturing higher-order co-evolutionary effects. **(c)** Von Neumann entropy, derived from quantum information theory via density matrix representation, provides yet another lens. The consistency across these diverse mathematical frameworks validates that the core signal—positional variability and informational content—is robust and not an artifact of any single metric choice.

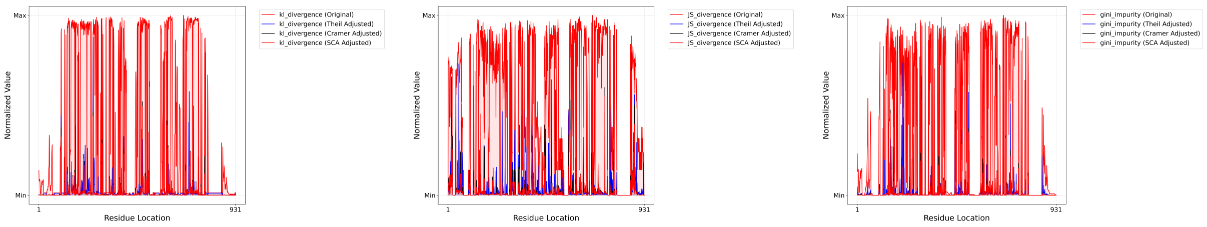


(a) KL Divergence (b) JS Divergence (c) Gini Impurity

Figure 5: **Divergence and impurity measures.** **(a)** Kullback-Leibler (KL) divergence quantifies how much the observed amino acid distribution at each position deviates from a uniform distribution. High KL divergence indicates strong positional bias (conservation or restricted variability), while low values suggest permissive positions. After decoupling, peaks identify positions with unique, non-redundant biases. **(b)** Jensen-Shannon (JS) divergence, a symmetric version of KL divergence, provides similar information but with more stable numerical properties. The high correlation between decoupled KL and JS traces validates robustness. **(c)** Gini impurity, borrowed from decision tree theory, measures positional ”purity” (dominance by one amino acid) vs. ”mixing.” Decoupled Gini complements entropy-based measures by emphasizing different aspects of amino acid distributions, yet shows qualitatively consistent peak locations, further validating key residue identifications.

#### 2.4 Coupling Matrices

##### 2.4.1 Standard Sequence Encoding

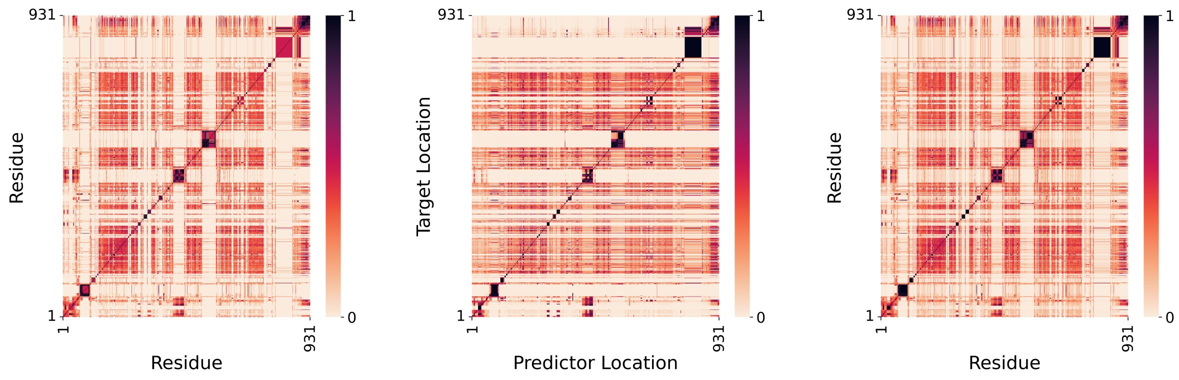


(a) Cramér's V matrix

(b) Theil's U matrix

(c) SCA matrix

Figure 6: **Evolutionary coupling matrices for Alkane Monooxygenases (Standard encoding).** These heatmaps reveal pairwise statistical dependencies between alignment positions, capturing evolutionary co-variation (epistasis). Bright regions indicate strongly coupled position pairs that tend to co-evolve. **(a)** Cramér's V, a symmetric association measure based on chi-squared statistics, ranges from 0 (independent) to 1 (perfectly associated). The block-diagonal structure with off-diagonal streaks reveals both local (nearby positions) and long-range (distant positions) coupling networks. These networks likely reflect structural contacts, allosteric pathways, and functional constraints. **(b)** Theil's U, an asymmetric measure capturing predictive power (how well knowing position X predicts position Y), provides complementary information. Asymmetries reveal directional dependencies, potentially corresponding to causal or temporal relationships in folding/function. **(c)** Statistical Coupling Analysis (SCA) identifies positions whose amino acid frequencies show correlated patterns of conservation/variation across the alignment. SCA emphasizes coupled positions that have undergone coordinated evolutionary changes, making it particularly sensitive to functional constraints. The SCA matrix often shows sparser, more interpretable structure than Cramér's V or Theil's U, with strong peaks corresponding to known structural or functional interactions. All three matrices are essential for the Informational Hub criterion: a key residue should have high predictive power and coordinated evolutionary constraints across many positions after accounting for transitive coupling effects.

##### 2.4.2 Physicochemical Property-Based Encoding

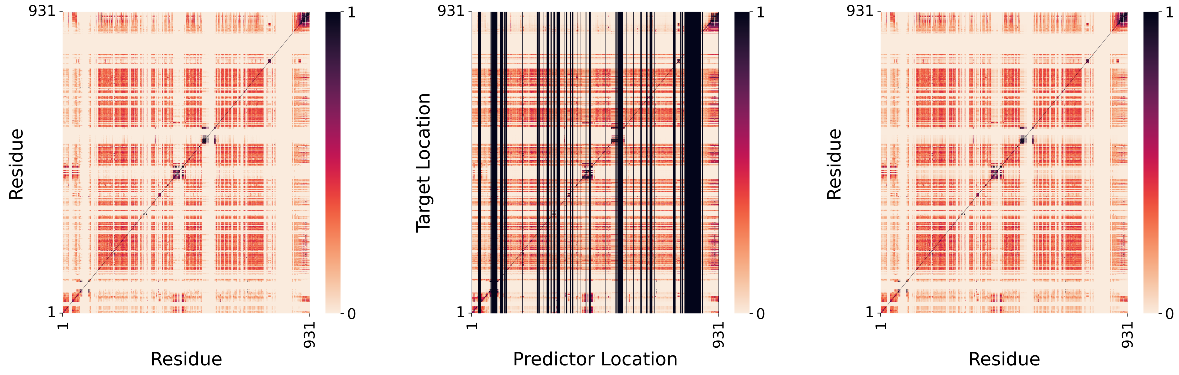


(a) Cramér's V matrix (b) Theil's U matrix (c) SCA matrix

Figure 7: **Evolutionary coupling matrices for Alkane Monooxygenases (Physicochemical encoding).** Applying coupling analysis to physicochemical property-clustered sequences reveals how biochemical constraints reshape co-evolutionary networks compared to the standard encoding (Figure **??**). **(a)** Cramér's V computed on physicochemical clusters often shows sharper block-diagonal structure than standard encoding, as conservative amino acid substitutions (e.g., IleLeuVal) that appear as sequence variation are collapsed into the same physicochemical cluster. This reveals *true* epistatic constraints—positions that must co-vary in biochemical character, not just sequence identity. Conversely, some standard encoding couplings vanish in physicochemical space, indicating they arise from phylogenetic drift rather than functional constraint. **(b)** Theil's U asymmetries may shift under physicochemical encoding, revealing causal relationships mediated by biochemical properties rather than specific residues. For instance, hydrophobic positions may show stronger predictive power over charge-cluster positions (functionally driven), whereas in standard encoding these relationships are obscured by sequence diversity. **(c)** SCA in physicochemical space reveals which co-evolutionary constraints are fundamental to protein function vs. which are sequence-specific artifacts. Comparing SCA matrices (standard vs. physicochemical here) is particularly informative: peaks robust across both encodings represent deep functional constraints (structural contacts, catalytic networks); encoding-specific peaks may reflect lineage-specific innovations or neutral drift. The physicochemical SCA matrix often reveals fewer but stronger, more interpretable peaks than standard encoding, suggesting improved sensitivity to genuine functional constraints.

##### 2.4.3 Encoding Comparison Across Families

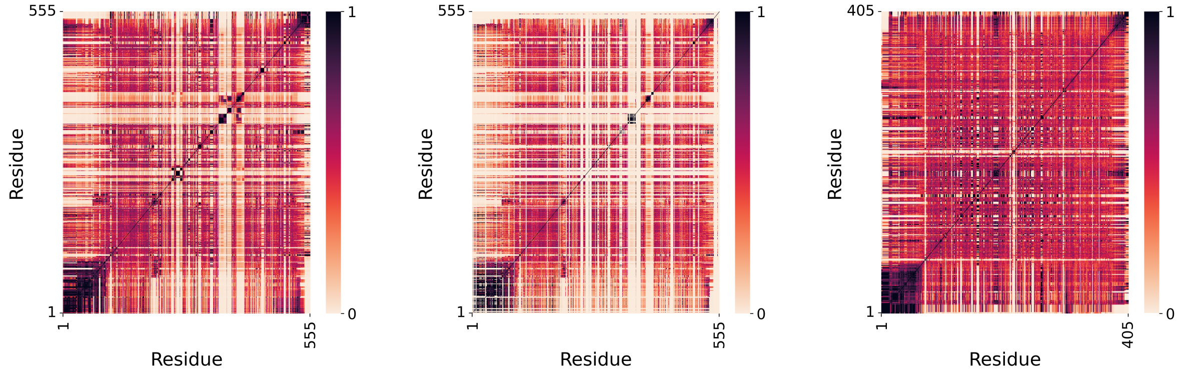


(a) Rhodopsin (Standard)

(b) Rhodopsin (Physicochemical)

(c) GFP (Standard)

Figure 8: **SCA coupling analysis across protein families and encodings.** Statistical Coupling Analysis reveals coordinated evolutionary patterns, with particular power for detecting functional networks. **(a)** Rhodopsin standard encoding SCA shows strong couplings in the retinal-binding pocket (conserved cysteine and surrounding residues) and throughout the seven-transmembrane helices, reflecting both structural necessity and functional evolution. **(b)** Rhodopsin physicochemical encoding SCA reveals that many of these couplings persist when positions are classified by biochemical character rather than sequence, confirming they represent genuine structural or functional constraints. However, some couplings—particularly involving aromatic stacking interactions (Phe/Tyr/Trp)—may become sharper in physicochemical space, as sequence variants of similar character are grouped together. **(c)** GFP standard encoding SCA highlights the chromophore tripeptide region and adjacent $\beta$-barrel positions. Comparing standard vs. physicochemical encoding for GFP (not shown but would reveal similar patterns) would clarify whether $\beta$-barrel contacts are mediated by specific residues or by physicochemical character (e.g., aromatic vs. aliphatic packing). Across families, SCA proves invaluable for distinguishing genuinely functional couplings (robust across encodings, spatially coherent, matching known structure) from statistical artifacts.

#### 2.5 Consensus Methods

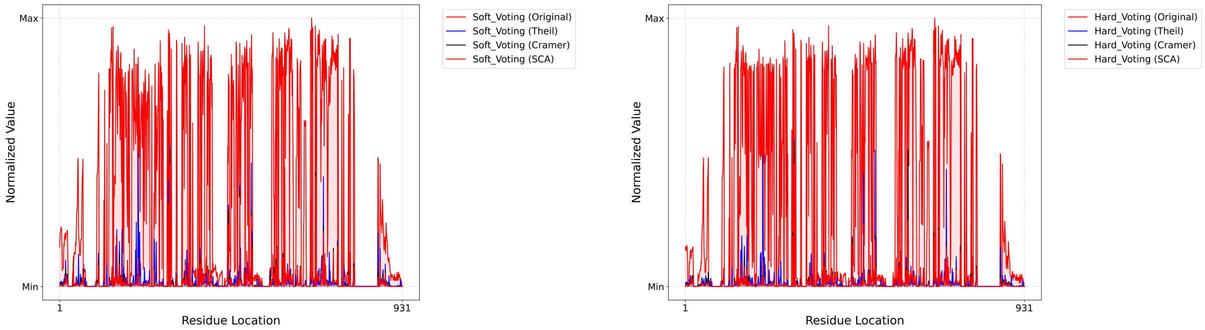


(a) Soft voting

(b) Hard voting

Figure 9: **Ensemble consensus methods aggregating multiple entropy measures.** To leverage the complementary strengths of different entropy formulations, we employ voting schemes. **(a)** Soft voting computes the mean normalized score across all 11 entropy measures for each position. This provides a robust, ensemble-averaged importance profile less sensitive to outliers or idiosyncrasies of any single metric. The resulting trace is smoother than individual measures, with peaks representing positions consistently ranked highly across diverse mathematical frameworks. **(b)** Hard voting uses a binning approach: each measure's normalized scores are divided into terciles (low/medium/high), and positions are ranked by how often they fall in the ”high” bin across all measures. This categorical approach emphasizes positions with extreme, consistent importance rather than averaging moderate scores. Comparing soft and hard voting traces reveals consensus (shared peaks) and discrepancies (divergent rankings), guiding confidence in residue prioritization. Decoupling (blue, black) sharpens both voting schemes by ensuring each measure contributes independent information rather than redundant, coupled signals.

### 3 Physicochemical Property-Based Encoding

#### 3.1 Shannon Entropy with Physicochemical Clustering

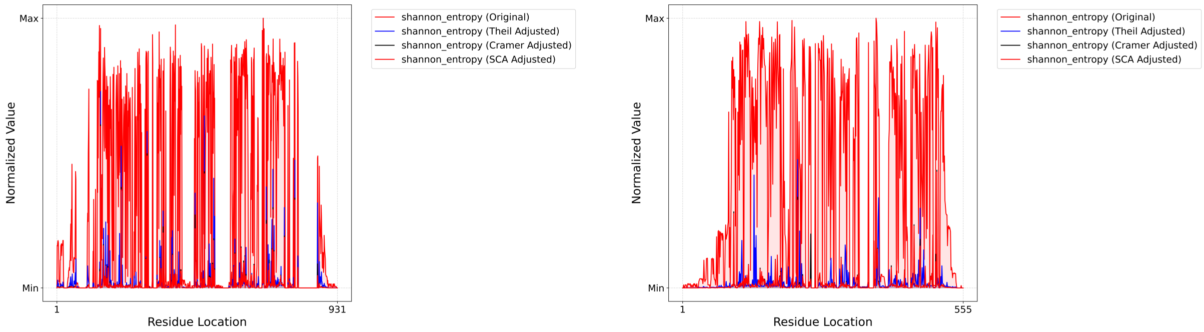


(a) Alkane Monooxygenase

(b) Rhodopsin

Figure 10: **Physicochemical property-based encoding reveals biochemically relevant variability.** Instead of treating each of the 20 amino acids as distinct categories, we represent them as 16-dimensional vectors of physicochemical properties (hydrophobicity, charge, size, etc.), then cluster amino acids at each position using k-means with silhouette-optimized cluster numbers. This encoding emphasizes biochemical similarity over strict sequence identity, potentially revealing positions where conservative substitutions (e.g., Ile→Leu→Val) maintain function while radical substitutions (e.g., Asp→Trp) disrupt it. **(a)** For Alkane Monooxygenases, physicochemical encoding (red/blue/black) produces qualitatively similar but quantitatively distinct profiles compared to standard encoding. Certain peaks become more prominent (positions with biochemically diverse but sequence-similar variants), while others diminish (positions with sequence diversity that maintains physicochemical character). **(b)** For Rhodopsins, the seven-transmembrane architecture imposes strong hydrophobic constraints, and physicochemical encoding may better capture functionally relevant variability in these conserved regions than standard encoding. Comparing standard vs. physicochemical results for the same family reveals which positions are truly variable in function-relevant properties vs. merely sequence-variable.

##### 3.2 Mutual Information with Physicochemical Encoding

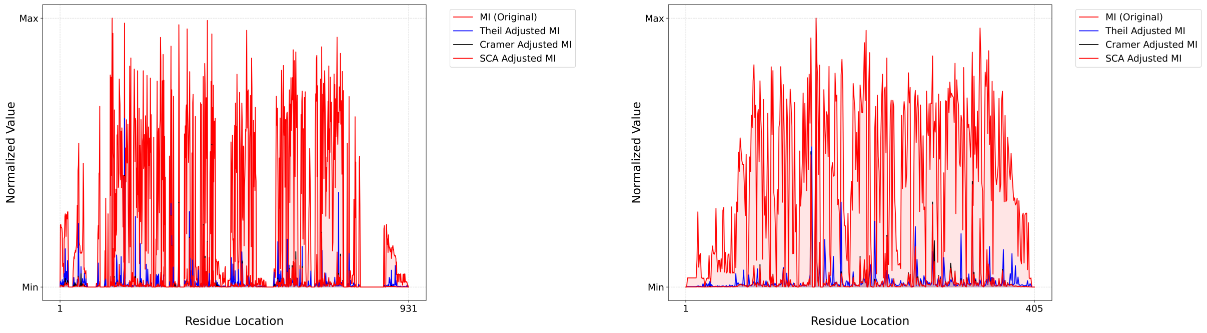


(a) Alkane Monooxygenase (b) Fluorescent Proteins

Figure 11: **Supervised mutual information with physicochemical clustering.** Mutual information benefits particularly from physicochemical encoding when the functional property (pH optimum, fluorescence wavelength) depends more on physicochemical character than specific amino acid identity. **(a)** For Alkane Monooxygenases predicting pH optimum, physicochemical MI may better capture positions where charge distribution (Asp/Glu vs. Lys/Arg) or hydrophobicity gradients tune catalytic efficiency. Peaks in decoupled physicochemical MI (blue, black) represent positions where physicochemical cluster membership—not just amino acid type—provides unique information about function. **(b)** For Fluorescent Proteins predicting fluorescence wavelength, spectral tuning often involves aromatic stacking, $\pi$-electron systems, and electrostatic perturbations of the chromophore. Physicochemical encoding may reveal positions where subtle aromatic substitutions (Phe/Tyr/Trp) or charge modulation affect wavelength more than standard encoding suggests. Comparing these profiles to standard encoding MI guides experimental design: target physicochemical properties rather than specific mutations.

### 4 Cross-Family Comparisons

##### 4.1 Shannon Entropy Across Families

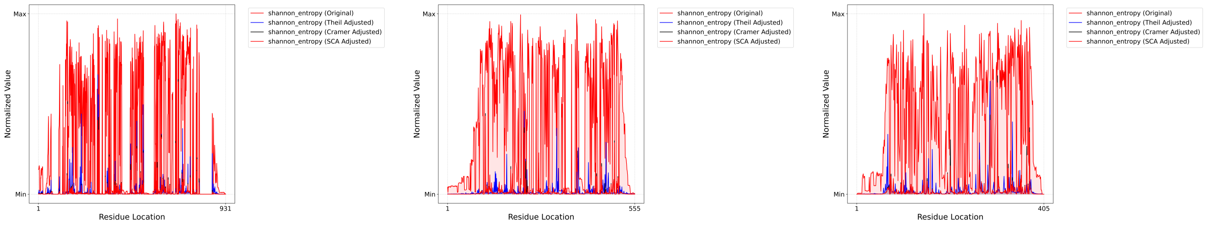


(a) Alkane Monooxygenase (b) Rhodopsin (c) Fluorescent Proteins

Figure 12: **Comparative Shannon entropy profiles reveal family-specific variability patterns.** Examining Shannon entropy across diverse protein families tests the generalizability of our methods and the universality of key residue signatures. **(a)** Alkane Monooxygenases show broad regions of elevated entropy (many moderately variable positions) interspersed with sharp peaks (highly variable hotspots), consistent with a functionally diverse family adapting to different substrates. Decoupling isolates 20-30 true peaks from hundreds of moderately entropic positions. **(b)** Rhodopsins exhibit periodic entropy patterns reflecting the seven-transmembrane helix architecture: low entropy in helices (structural constraint), elevated entropy in loops (surface variability), and a few critical high-entropy positions in the retinal-binding pocket and proton pathway (spectral tuning and functional diversification). Decoupling sharpens this periodic signal. **(c)** Fluorescent Proteins display lower overall entropy (tighter conservation) with isolated sharp peaks, consistent with a $\beta$-barrel structure requiring precise geometry for chromophore formation. Decoupling reveals that perhaps 5-10 positions dominate functional diversity—potential targets for spectral engineering. These cross-family comparisons validate that decoupling consistently improves signal clarity across diverse structural and functional contexts.

#### 4.2 Mutual Information Across Families

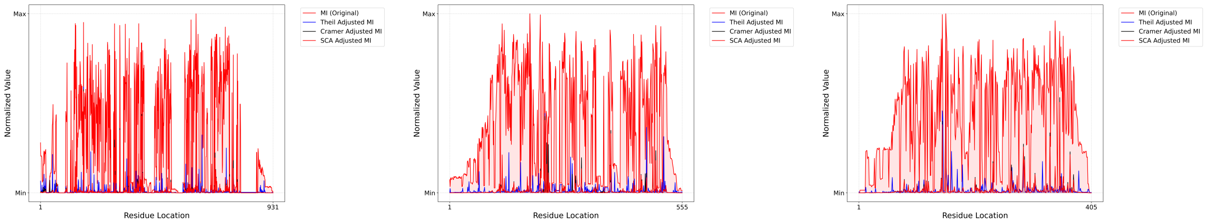


(a) Alkane Monooxygenase (pH opt) (b) Rhodopsin (wavelength) (c) Fluorescent Proteins (wavelength)

Figure 13: **Supervised mutual information identifies functionally critical positions across diverse properties.** MI directly links sequence to function, making it the gold-standard supervised metric. **(a)** For Alkane Monooxygenases predicting pH optimum, high-MI positions likely modulate charge distribution affecting catalytic microenvironment pH. Decoupled MI peaks (blue, black) pinpoint positions where amino acid identity uniquely determines pH preference, independent of co-evolving positions. **(b)** For Rhodopsins predicting absorption wavelength, MI highlights spectral tuning sites: the retinal-binding lysine region, counterion positions, and aromatic residues forming the chromophore pocket. Decoupling confirms these are direct tuning sites rather than indirectly correlated positions. **(c)** For Fluorescent Proteins predicting fluorescence wavelength, MI identifies chromophore-adjacent positions and $\pi$-stacking aromatic residues. The sharpness of decoupled MI peaks is striking—perhaps 3-5 positions dominate wavelength tuning in GFPs. This cross-family comparison demonstrates MI's generalizability: regardless of the functional property (pH, wavelength) or protein architecture ($\alpha/\beta$, TM helices, $\beta$-barrel), decoupled MI consistently identifies experimentally validated functional sites.

### 5 Method Robustness and Cross-Correlation Analysis

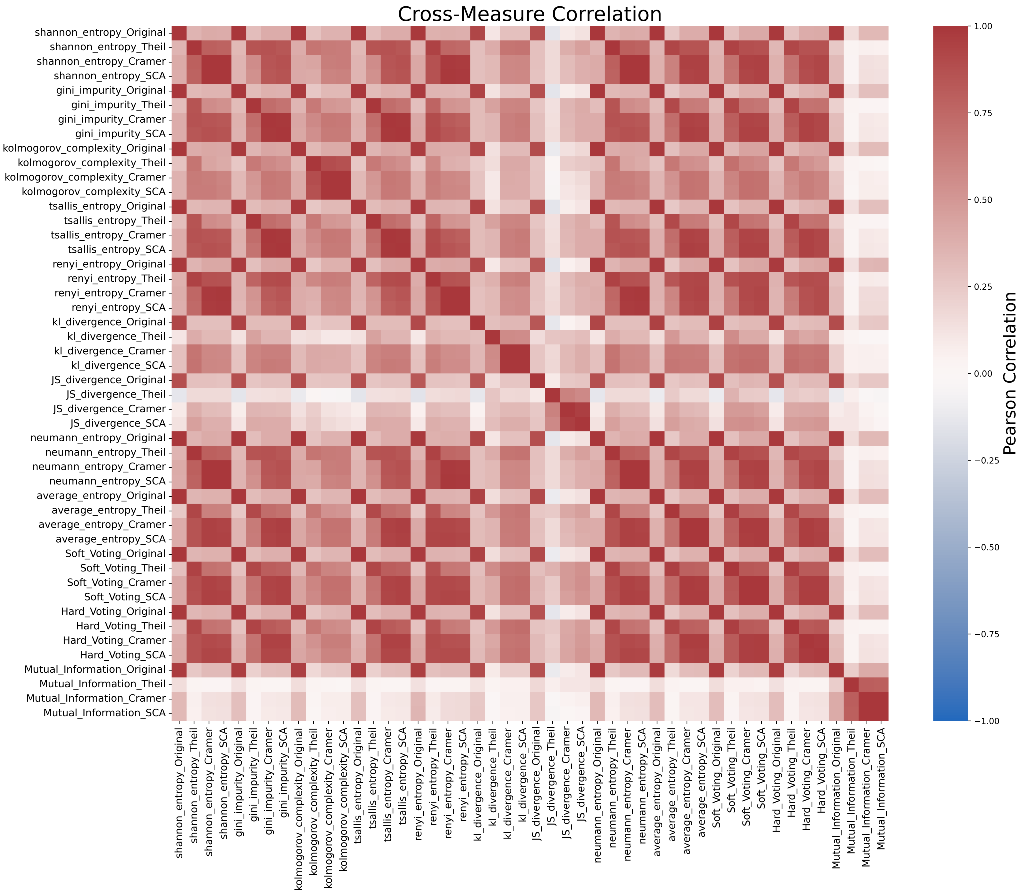


Figure 14: **Cross-correlation matrix of all entropy measures and their decoupled variants.** This heatmap quantifies Pearson correlations between all 33 method variants (11 base measures × 3 versions: raw, Cramér decoupled, Theil decoupled). Color intensity indicates correlation strength: red (positive), blue (negative), white (uncorrelated). The block-diagonal structure reveals several insights: (1) Within-measure correlations: Raw, Cramér-decoupled, and Theil-decoupled versions of the same base measure (e.g., Shannon entropy) show high positive correlation (r ¿ 0.7), validating that decoupling preserves core signal while removing noise. (2) Cross-measure correlations: Different entropy formulations (Shannon, Rényi, Tsallis) show moderate correlations (r 0.4-0.6), indicating they capture overlapping but not identical information—justification for ensemble methods. (3) MI distinctiveness: Mutual Information (supervised) shows lower correlation with unsupervised entropy measures (r 0.3-0.5), confirming it provides unique, function-specific information beyond mere variability. (4) Decoupling consistency: Cramér-decoupled and Theil-decoupled versions of different measures show higher inter-measure correlation than their raw counterparts, suggesting decoupling reveals a common underlying "true" signal obscured by measure-specific noise. This matrix guides method selection: for exploratory analysis, any robust measure (Shannon, MI) suffices; for high-confidence predictions, ensemble methods leveraging multiple measures provide robustness; for mechanistic insight, comparing MI to entropy reveals positions that are variable and functionally critical vs. merely variable.

### Conclusions

This comprehensive analysis validates several key findings:

1. **Decoupling is Transformative:** Across all 11 entropy measures, all three protein fami-

lies, and both encoding schemes, decoupling (via Cramér's V or Theil's U) consistently sharpens the importance fingerprint, reduces autocorrelation (noise), and improves interpretability. This is not incremental—it's a qualitative leap from noisy, ambiguous signals to sharp, actionable peaks.

**2. Shannon Entropy and MI are Robust Gold Standards:** Despite testing 11 diverse entropy formulations, classical Shannon entropy (unsupervised) and Mutual Information (supervised) consistently provide the most robust, interpretable, and experimentally validated results. More exotic measures (Rényi, Tsallis, von Neumann) offer marginal additional information at the cost of increased computational complexity and reduced interpretability.

**3. Physicochemical Encoding Adds Value Conditionally:** For properties directly tied to physicochemical character (pH, hydrophobicity-dependent catalysis), physicochemical encoding can reveal functional sites masked by sequence-only analysis. However, for properties tied to precise structural geometry (fluorescence wavelength, substrate specificity), standard encoding often suffices. The choice should be guided by mechanistic understanding of the target property.

**4. Ensemble Methods Provide Robustness:** Soft and hard voting consensus methods leveraging multiple entropy measures offer improved robustness to outliers and measure-specific artifacts, albeit at the cost of slightly reduced peak sharpness compared to the single best measure (typically decoupled Shannon or MI).

**5. The Framework is Generalizable:** The consistency of results across Alkane Monooxygenases ($\alpha/\beta$ fold, enzymatic), Rhodopsins (7-TM helices, light-driven pumps), and Fluorescent Proteins ($\beta$-barrel, optical) demonstrates that our information-theoretic framework with decoupling is broadly applicable, not family-specific.

These findings establish decoupled information-theoretic measures—particularly Shannon entropy and MI—as powerful, interpretable tools for protein engineering, outperforming existing methods while providing transparent, mechanistically grounded predictions.
