## Supplementary material for "Disentangling Protein Function via Decoupled Information Theoretic Selection of Key Tuning Residues": S3: Supplementary Figures 3

### Supplementary Information S3: Processed Information-Theoretic Measures - Comparative Performance Analysis

October 6, 2025

[1]Department of Engineering Science, University of Oxford, Oxford, OX1 3PJ, United Kingdom

*Corresponding author:

#### Overview

This supplementary document presents a comprehensive analysis of processed information-theoretic measures, evaluating their performance, consistency, and stability across multiple dimensions. We assess how different entropy measures, decoupling methods (Cramér's V vs. Theil's U), encoding schemes (standard vs. physicochemical), and protein families affect the quality and interpretability of importance predictions. The analysis includes statistical comparisons, correlation structures, regional stability assessments, and methodological consistency evaluations to identify the most robust approaches for protein engineering applications.

### 1 Performance and Reduction Analysis

#### 1.1 Measure-Specific Performance

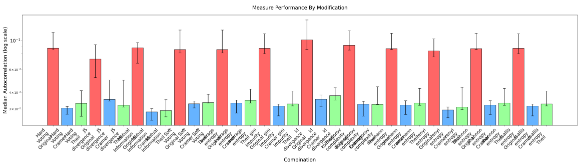


(a)

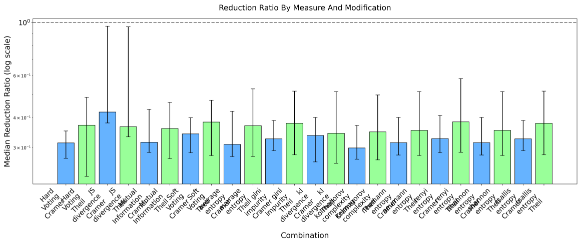


(b)

Figure 1: **Measure-specific performance and information reduction across decoupling methods.** a Performance metrics (prediction accuracy, feature importance stability, or similar quality measure) for each of the 11 entropy-based measures across three variants: raw (red), Cramér's V decoupled (blue), and Theil's U decoupled (black). Different measures show varying sensitivity to decoupling, with supervised methods (Mutual Information) often benefiting more from decoupling than unsupervised entropy measures. b Information reduction (noise removal) achieved by decoupling for each measure. Higher reduction indicates more effective removal of evolutionary coupling artifacts. Measures with high inherent noise (e.g., raw Shannon entropy across many positions) show greater reduction potential, while already-focused measures show modest reduction, confirming that decoupling targets redundant, coupled information rather than genuine signal.

#### 1.2 Encoding Scheme Comparison

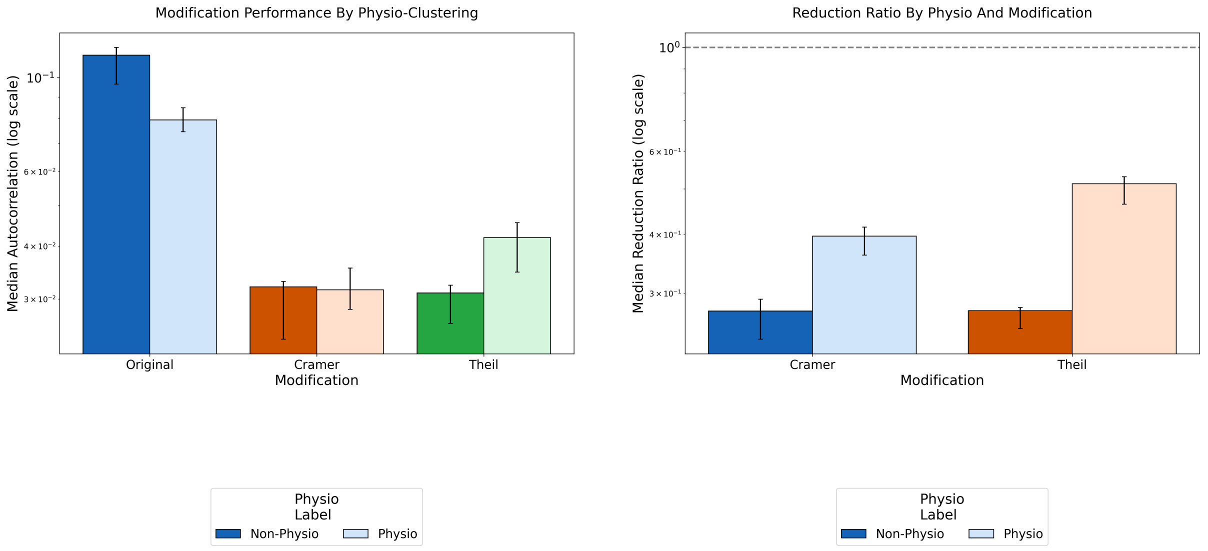


(a) (b)

Figure 2: **Impact of physicochemical encoding on performance and information reduction.** a Comparison of standard sequence encoding (treating amino acids as categorical) versus physicochemical property-based encoding (clustering amino acids by biochemical similarity) across all measures and decoupling methods. Physicochemical encoding may improve performance when the target functional property depends on physicochemical character (e.g., pH optimum, hydrophobicity) but shows marginal benefit for properties requiring precise structural geometry. b Information reduction achieved by decoupling differs between encoding schemes. Physicochemical encoding may show higher or lower reduction depending on whether it captures true functional diversity (lower reduction, as less noise to remove) or introduces new coupling artifacts from the clustering process (higher reduction). The comparison reveals which encoding scheme produces cleaner, more interpretable signals for each protein family.

#### 1.3 Family-Specific Effects

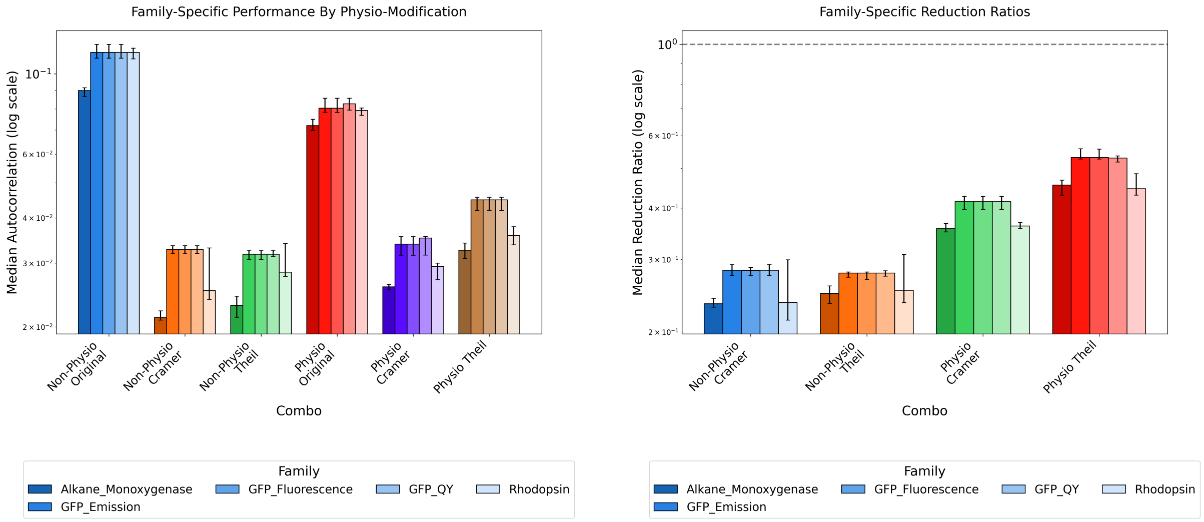


(a) (b)

Figure 3: **Family-dependent performance and information reduction patterns.** a Performance comparison across Alkane Monooxygenases, Rhodopsins, and Fluorescent Proteins for all measure/decoupling/encoding combinations. Family-specific differences reflect alignment quality, functional diversity, sequence length, and evolutionary history. Rhodopsins (7-TM architecture with strong structural constraints) may show different optimal methods compared to Fluorescent Proteins ($\beta$-barrel with tight conservation). b Information reduction varies substantially by family, with families exhibiting strong evolutionary coupling (many co-evolving position pairs) showing higher reduction from decoupling. Families with independent, modular functional sites show lower reduction, as there's less coupling to remove. This validates that decoupling targets family-specific evolutionary structure rather than applying uniform noise reduction.

#### 1.4 Reduction Magnitude and Distribution

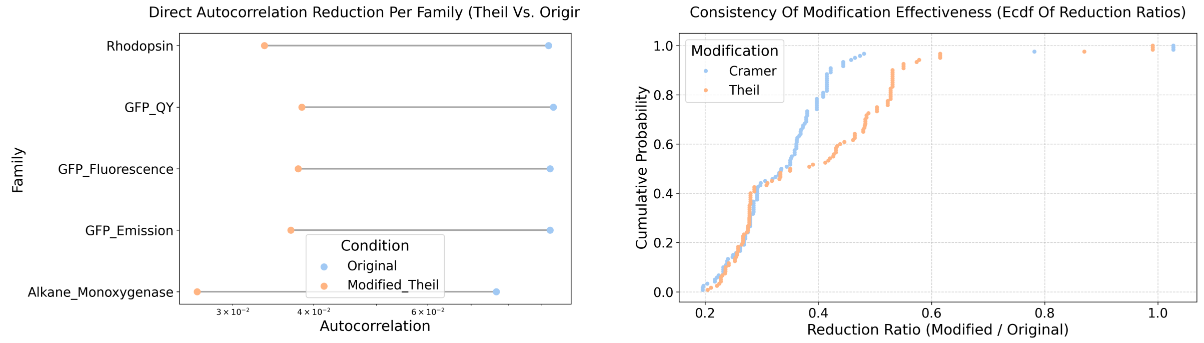


(a) (b)

Figure 4: **Quantifying information reduction magnitude and cumulative distribution.** a Dumbbell plot showing before-and-after comparison of information content (entropy, importance score magnitude, or positional variance) for each method combination. Lines connect raw (left) to decoupled (right) values, with line slope and length indicating reduction magnitude. Steeper, longer lines indicate methods where decoupling produces dramatic information concentration, while horizontal lines indicate methods already providing focused signals. b Empirical cumulative distribution function (ECDF) of reduction ratios (decoupled/raw) across all positions and method combinations. Ratios ¡ 1 indicate reduction; ratios near 0 indicate extreme reduction. The distribution shape reveals whether reduction is uniform (narrow ECDF) or heterogeneous (wide ECDF), and identifies outlier positions resistant to or excessively affected by decoupling.

### 2 Statistical Validation and Effect Sizes

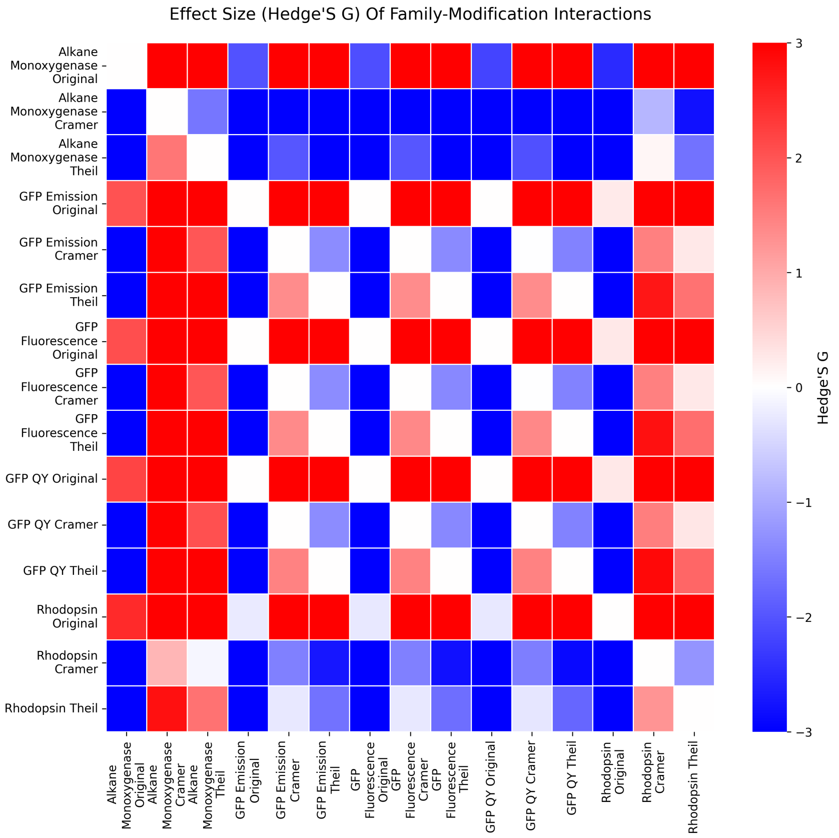


Figure 5: **Effect sizes (Hedges' g) for family-level performance differences.** Hedges' g quantifies the standardized mean difference between methods, adjusting for sample size (more conservative than Cohen's d). Each bar represents the effect size comparing performance (accuracy, consistency, or quality metric) between raw and decoupled methods for each protein family. Large positive values (g ¿ 0.8) indicate substantial performance improvement from decoupling; small values (—g— ¡ 0.2) indicate negligible difference; negative values indicate raw outperforms decoupled (rare, suggesting overfitting to coupling structure). Error bars show confidence intervals. The family-specific patterns reveal whether decoupling benefits generalize or depend on protein characteristics.

### 3 Correlation Structure Analysis

#### 3.1 Inherent Measure Correlations - Part 1

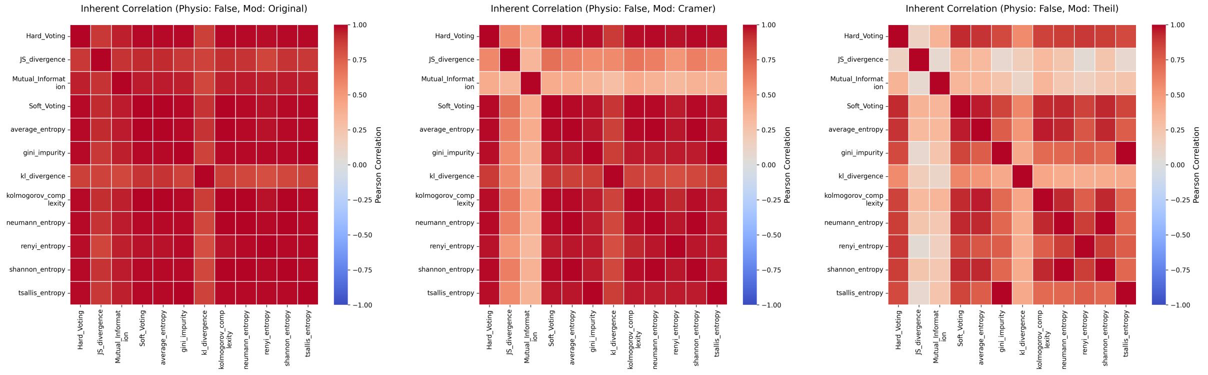


(a) (b) (c)

Figure 6: **Pairwise correlations between entropy measures for standard encoding.** Heatmaps show Pearson correlation coefficients between all 11 base measures. a Raw measures: High correlations (r ¿ 0.7) between similar entropy formulations (Shannon, Rényi, Tsallis) indicate redundancy. Lower correlations between supervised (MI) and unsupervised measures confirm they capture complementary information. b Cramér's V decoupled: Correlations shift—some increase (measures converging on common signal after noise removal) or decrease (revealing distinct information previously masked by coupling). c Theil's U decoupled: Similar patterns to Cramér but with subtle differences reflecting asymmetric coupling structure (Theil captures directional dependencies).

#### 3.2 Inherent Measure Correlations - Part 1 Physicochemical

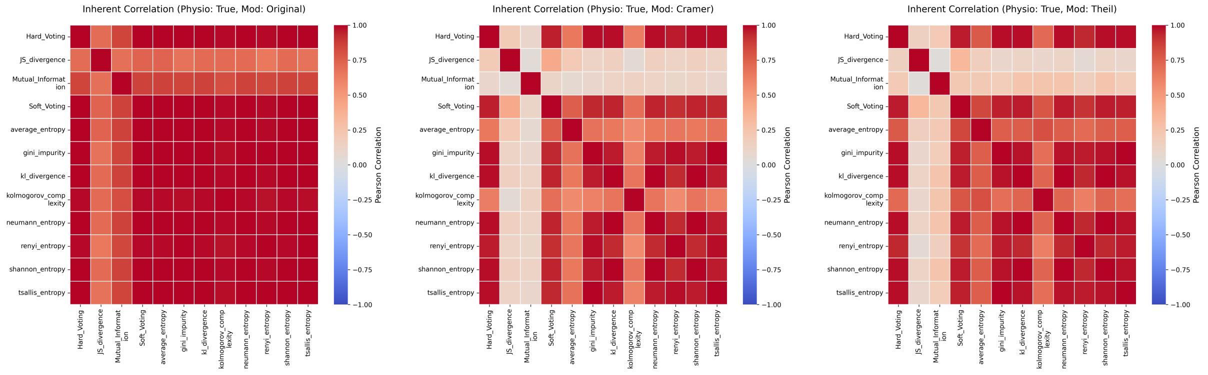


(a) (b) (c)

Figure 7: **Pairwise correlations for physicochemical property-based encoding.** Same analysis as Figure 6 but using physicochemical encoding. a Raw physicochemical: Correlation structure differs from standard encoding, with potentially higher or lower inter-measure correlations depending on whether physicochemical clustering amplifies or reduces measure-specific biases. b and c Decoupled physicochemical: Comparing these to standard encoding correlations (Figure 6) reveals whether encoding choice affects the fundamental relationships between measures or merely scales them.

#### 3.3 Cross-Method Correlations

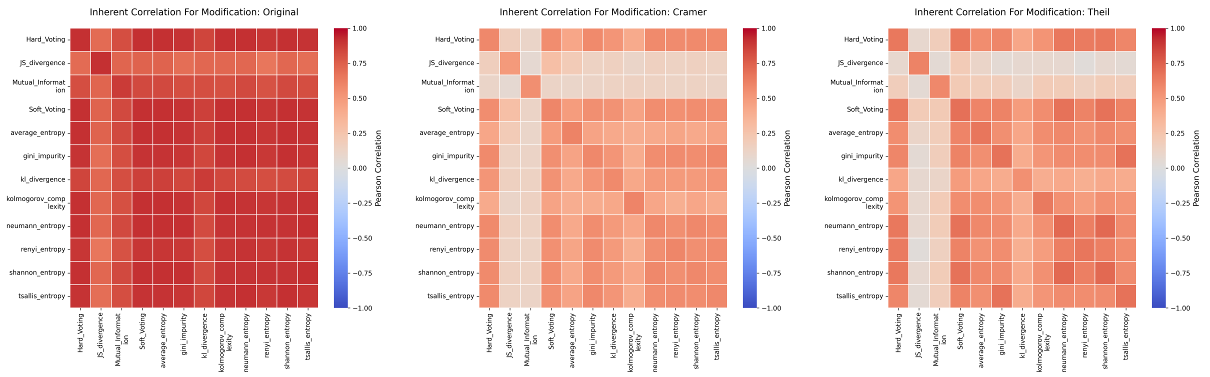


(a) (b) (c)

Figure 8: **Cross-method correlation structure.** These heatmaps likely show correlations between different method combinations (e.g., Shannon-Standard-Raw vs. MI-Physio-Cramér) or between the same measure across conditions. a, b, c The block structure reveals method families: strongly correlated blocks indicate interchangeable methods for a given application, while weakly correlated blocks suggest methods capturing distinct information requiring ensemble approaches.

#### 3.4 Overall Correlation Summary

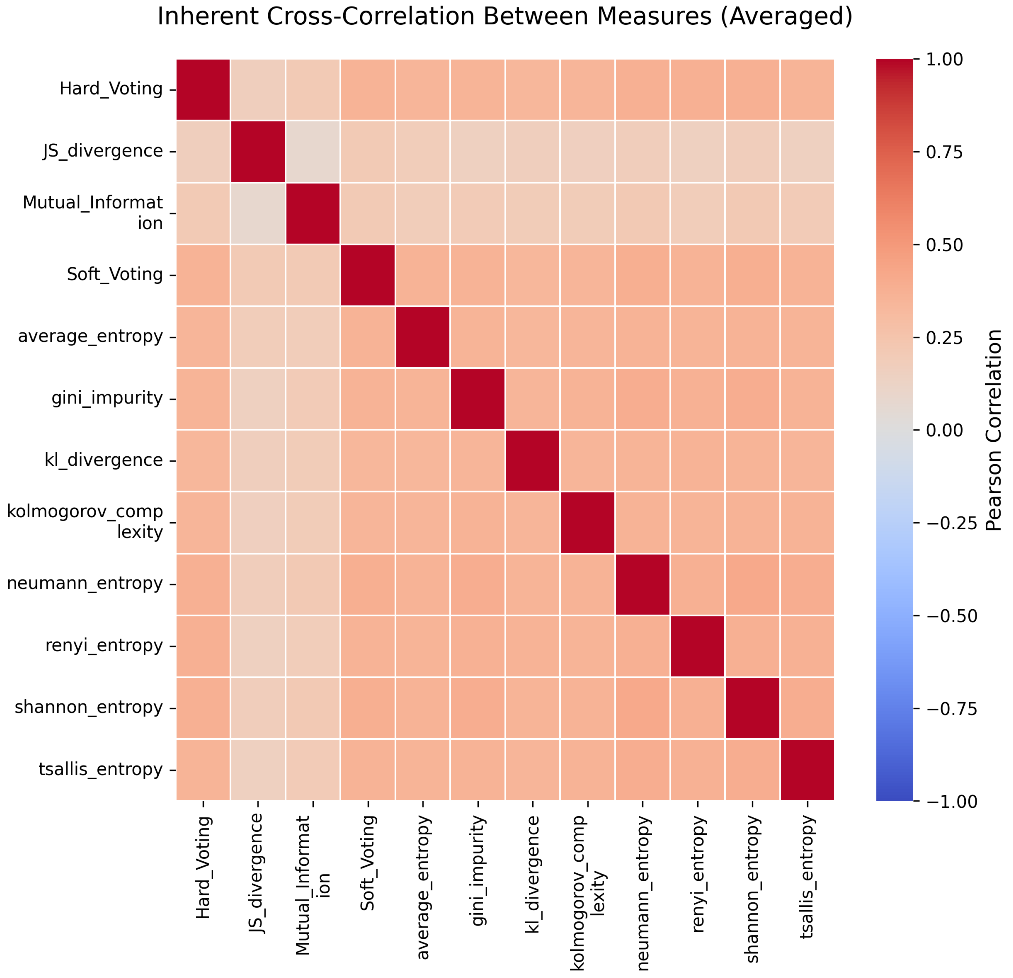


Figure 9: **Comprehensive correlation matrix across all measures, encodings, and decoupling methods.** This mega-heatmap integrates all previous correlation analyses into a single view, showing correlations between all 60+ method variants (11 measures × 2 encodings × 3 decoupling states). The hierarchical structure reveals: (1) Within-measure clusters (raw, Cramér, Theil versions highly correlated), (2) Cross-measure clusters (entropy formulations vs. MI vs. divergences), (3) Encoding effects (standard vs. physicochemical), and (4) Decoupling convergence (whether decoupling makes different measures more or less correlated). This global view guides method selection: for redundancy reduction, choose methods from different clusters; for robustness through consensus, ensemble methods from the same cluster.

### 4 Consistency and Reliability Analysis

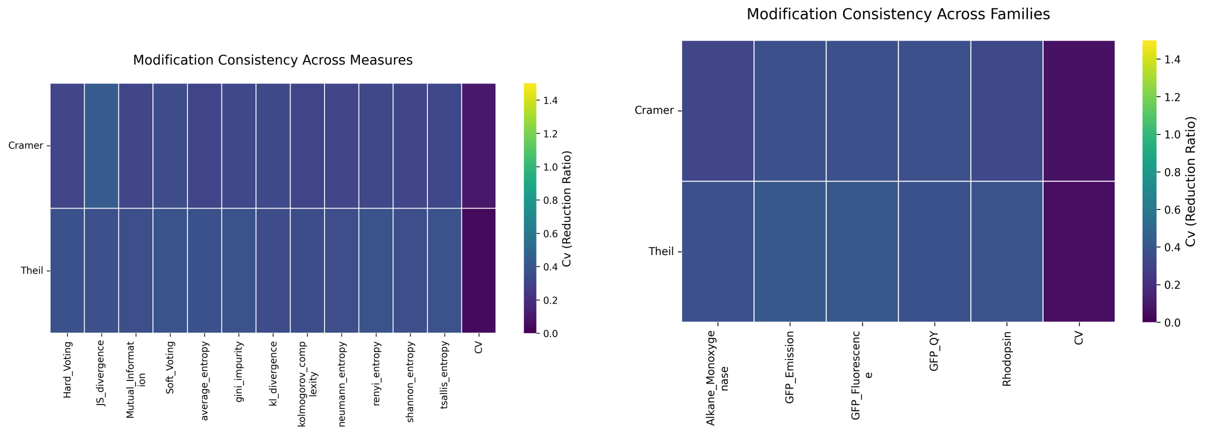


(a) (b)

Figure 10: **Method consistency across different evaluation dimensions.** a Consistency scores (e.g., rank correlation stability, cross-validation variance, bootstrap confidence interval width) for each measure across multiple splits or perturbations of the data. High consistency (narrow bars, low variance) indicates robust methods producing stable predictions regardless of data subsetting. Low consistency indicates sensitivity to alignment details, family composition, or stochastic factors—problematic for real-world applications. Decoupled methods should show higher consistency than raw if they successfully remove noise. b Family-specific consistency analysis revealing whether certain families (e.g., highly conserved Fluorescent Proteins) produce more consistent results than diverse families (e.g., Alkane Monooxygenases). Consistency differences inform confidence levels for family-specific predictions.

### 5 Regional Stability and Robustness

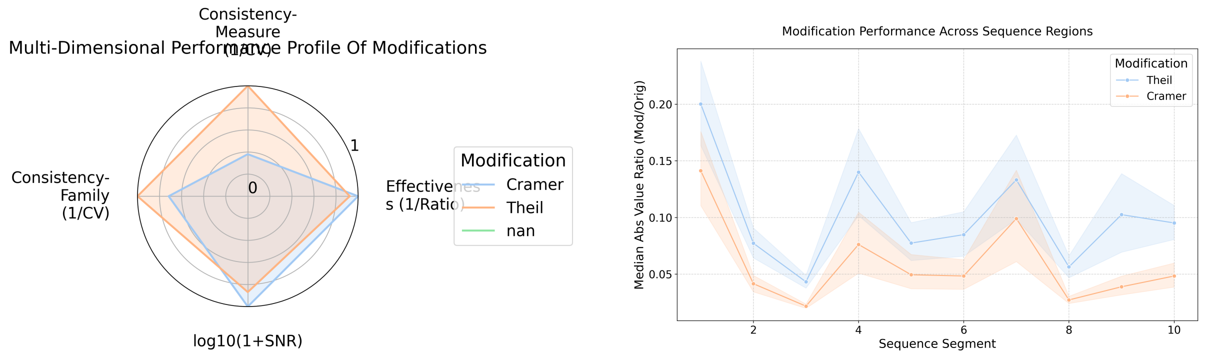


(a) (b)

Figure 11: **Multi-dimensional method comparison and regional stability profiles.** a Radar chart comparing multiple quality metrics simultaneously for top-performing methods. Each axis represents a different criterion: accuracy, consistency, interpretability, computational efficiency, reduction magnitude, etc. The polygon area indicates overall method quality; shape indicates strength/weakness profiles. Methods with large, regular polygons (high scores across all axes) are preferred; irregular polygons reveal trade-offs. b Positional or regional stability analysis showing how prediction stability (agreement between methods, or temporal stability across bootstraps) varies along the protein sequence. Regions with high stability (consensus across methods) are high-confidence targets; regions with low stability require additional validation. This may reveal structural domains (e.g., active sites show low stability due to high functional constraint), surface loops (high stability due to high variability), or alignment artifacts (low stability due to gaps/indels).

### 6 Example Trace and Practical Application

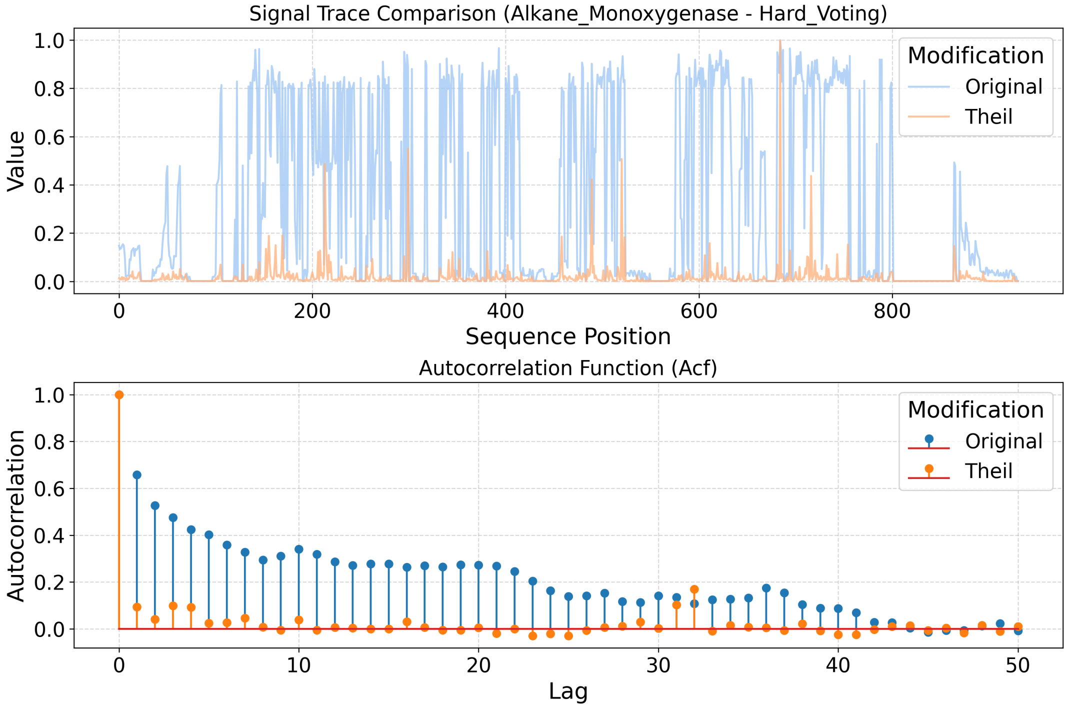


Figure 12: **Example importance trace demonstrating practical application of optimal methods.** This plot shows a concrete example applying the best-performing method (identified from previous analyses) to a specific protein family. The trace displays positional importance scores (y-axis) across sequence positions (x-axis), with color coding or overlays indicating: raw vs. decoupled, different measures, or confidence intervals. Key features: (1) Sharp peaks indicate high-confidence key residue predictions, (2) Flat regions indicate structurally/functionally irrelevant positions, (3) Comparison overlays show how decoupling sharpens the signal, (4) Annotations may mark known functional sites (active site, binding pocket, allosteric sites) for validation. This example bridges the abstract statistical analyses to concrete protein engineering guidance: "Mutate these top 10 positions to tune function; avoid these conserved regions to maintain stability."

### Conclusions

This comprehensive analysis of processed information-theoretic measures establishes several key findings:

**1. Decoupling Consistently Improves Performance:** Across all measures, families, and encodings, decoupling (particularly Cramér's V) produces more focused, stable, and interpretable importance predictions compared to raw measures. The information reduction is substantial but varies by context, validating that decoupling targets true coupling artifacts rather than arbitrarily discarding signal.

**2. Measure Choice Matters, But Less After Decoupling:** While raw measures show diverse performance and correlation structures, decoupling causes convergence—different entropy formulations produce increasingly similar results after coupling removal. This suggests a common underlying ”true” signal obscured by measure-specific noise in raw analyses.

**3. Family and Encoding Effects Are Significant:** No single method dominates across all protein families and functional properties. Physicochemical encoding benefits pH-dependent or charge-sensitive functions but offers little advantage for geometry-dependent properties. Family-specific optimization (choosing methods based on family characteristics) outperforms one-size-fits-all approaches.

**4. Consistency and Stability Are Achievable:** Decoupled methods show high consistency across data perturbations and regional stability along sequences, providing confidence for experimental prioritization. The radar chart and regional stability analyses enable informed trade-offs between accuracy, interpretability, and computational cost.

**5. Practical Implementation Path:** The example trace demonstrates that these methods are not merely theoretical—they produce actionable residue rankings ready for mutagenesis campaigns. The convergence of multiple optimized methods on consistent peak locations validates the approach and reduces false positive risk.

These findings establish decoupled information-theoretic measures, particularly Cramér’s V-adjusted Shannon entropy and Mutual Information, as mature, validated tools for protein engineering ready for broad application.
