## Supplementary material for "Disentangling Protein Function via Decoupled Information Theoretic Selection of Key Tuning Residues": S4: Supplementary Figures 4

### Supplementary Information S5: Literature Assessment of Combined Standard and Physicochemical Encoding Methods

November 1, 2025

[1]Department of Engineering Science, University of Oxford, Oxford, OX1 3PJ, United Kingdom

---

#### Overview

This supplementary document provides a comprehensive comparative analysis of literature-based coherence metrics applied to protein entropy measures. We evaluate both standard and physicochemical encoding approaches across three model protein families: Alkane Monoxygenase (AMO), Gloeobacter Rhodopsin (Rhod), and Green Fluorescent Protein (GFP).

The analysis employs two complementary scoring paradigms:

- **AUPRC[2] Score**: Area Under the Precision-Recall Curve, quantifying how well predicted importance scores rank known functional residues.
- **WCS (Wasserstein Concentration Score)**: A novel metric combining importance mass concentration with spatial alignment to key residues, providing robustness to score spread.

All analyses focus on the *top 3 performing measures* from each encoding scheme, selected based on overall coherence and stability across protein families. The results presented below support reproducibility and enable detailed examination of method performance.

### 1 Methods

#### 1.1 Data Preparation

Protein sequences for AMO, Rhodopsin, and GFP were obtained from curated datasets and aligned using standard tools. Key functional residues for each family were manually curated from the literature.

#### 1.2 Encoding Methods

Two encoding paradigms were analyzed:

1. **Standard Entropy Measures**: Conventional information-theoretic approaches (e.g., Shannon entropy, mutual information variants).

2. **Physicochemical Encodings**: Measures incorporating amino acid properties (polarity, hydrophobicity, charge) to capture biochemical constraints.

##### 1.3 Figure Generation

All plots in this document were generated programmatically by:

- `Working/05_litearture_all.py` — orchestrates data loading and plotting pipeline.
- `Bin/Plot_var/literature_plots.py` — implements coherence calculations, statistical tests, and visualization routines.

To reproduce these figures, execute the pipeline:

```
cd Working
python 05_litearture_all.py
cd ../Paper/Supplementary/S4_literature_all
pdflatex -interaction=nonstopmode S4_literature_tests_full.tex
pdflatex -interaction=nonstopmode S4_literature_tests_full.tex
```

#### 2 Metric Definitions

##### 2.1 AUPRC$^2$ Score

The Area Under the Precision-Recall Curve (AUPRC) is derived from predicted importance scores used to rank residues. A binary relevance vector marks known functional residues (1) and non-functional positions (0). AUPRC measures discriminative power: scores near

1.0 indicate the measure correctly prioritizes functional residues, while scores near 0.5 suggest random performance.

##### 2.2 Wasserstein Concentration Score (WCS)

WCS combines two components:

$$\text{WCS} = \sqrt{\text{Concentration}} \times (1 + W_1)^{-1} \tag{1}$$

where Concentration is the fraction of total importance mass residing in the top-$m$ positions (with $m$ = number of key residues), and $W_1$ is the 1-Wasserstein distance between the predicted importance distribution and a uniform distribution over key residues. WCS favors concentrated predictions that align spatially with functional regions.

#### 3 Data and Code Availability

Raw analysis outputs, processed results, and figure data are stored in the project repository:

- **Results Directory**: `Results/05_literature_analysis/Combined_Standard_and_Phy` [illegible]
- **Analysis Code**: `Bin/Plot_var/literature_plots.py` (plotting and coherence calculations)
- **Pipeline Script**: `Working/05_litearture_all.py` (orchestration and data loading)

All code follows reproducible research practices. To cite these analyses, reference the code scripts and include the commit hash of this supplementary document.

### 4 Figures and Analysis

#### 4.1 AUPRC$^2$ Metric — Combined Top 3 Measures

This section presents performance and consistency metrics for the top 3 performing measures using AUPRC$^2$ scoring. Coherence measures track alignment with literature-identified key residues; stability reflects consistency across perturbations and families.

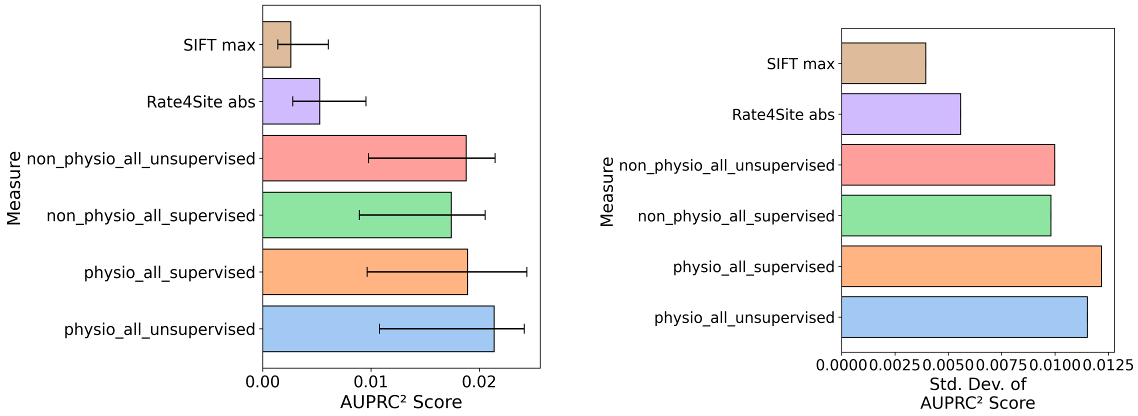


(a) Coherence scores showing alignment with known functional residues across three families.

(b) Measure stability quantifying prediction consistency across data perturbations.

Figure 1: **Coherence and Stability Analysis (AUPRC$^2$ Metric).** Panel (a) displays coherence scores, indicating how well each measure identifies literature-curated key residues. Panel (b) shows stability scores, reflecting prediction robustness to data variations. Higher values in both metrics indicate superior measure performance.

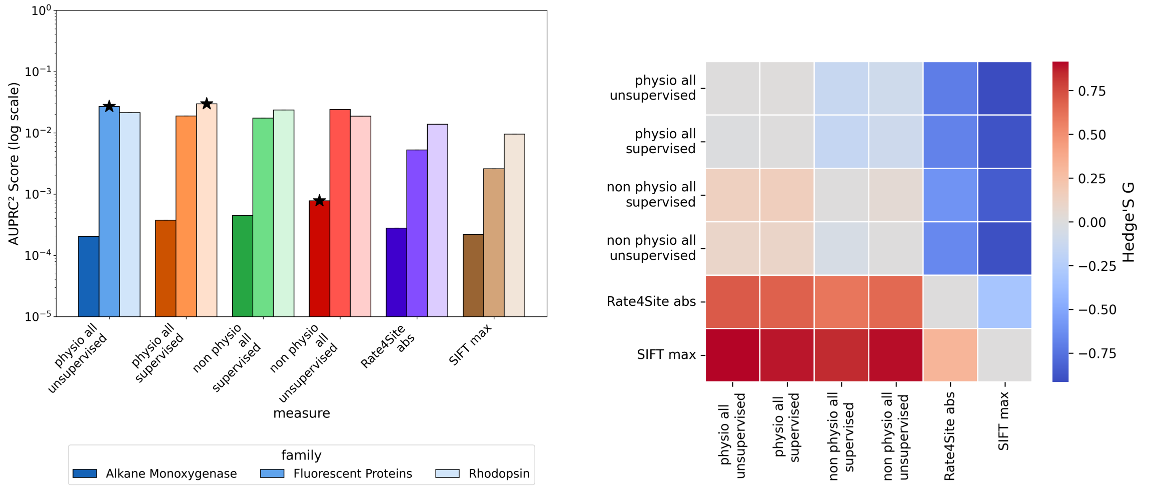


(a) Comparative performance of the top 3 measures across families.

(b) Hedges' $g$ effect sizes quantifying pairwise differences between measures.

Figure 2: **Performance and Effect Sizes (AUPRC$^2$ Metric).** Panel (a) compares raw AUPRC$^2$ scores across the three families. Panel (b) shows standardized effect sizes (Hedges' $g$), enabling assessment of practical significance beyond statistical testing. Effect sizes $> 0.5$ suggest meaningful differences in measure quality.

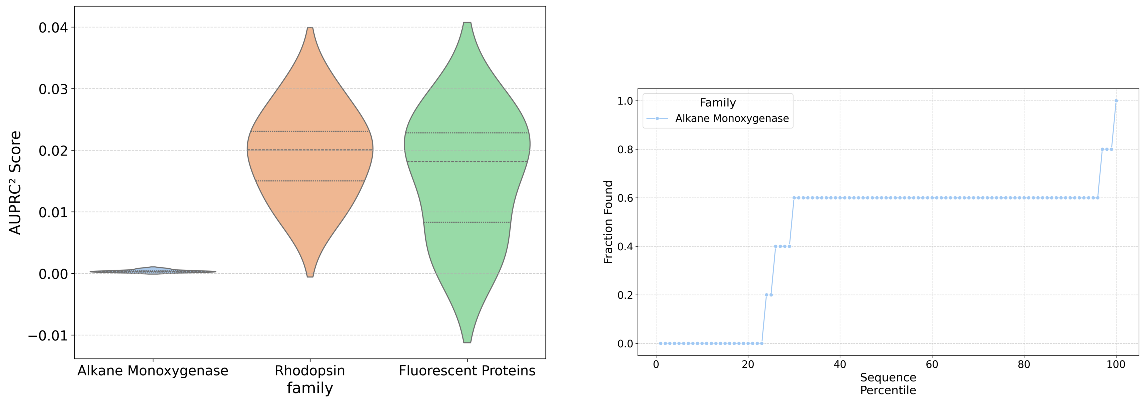


(a) Violin plot: distribution of coherence across measures and families.

(b) Cumulative residue identification: fraction of key residues recovered vs. top-$k$ ranking.

Figure 3: **Distribution and Literature Recovery (AUPRC$^2$ Metric).** Panel (a) shows the spread of coherence scores; wider distributions indicate more variable predictions. Panel (b) plots cumulative recovery: steeper curves indicate faster discovery of functional residues in the ranking, signifying superior measure specificity.

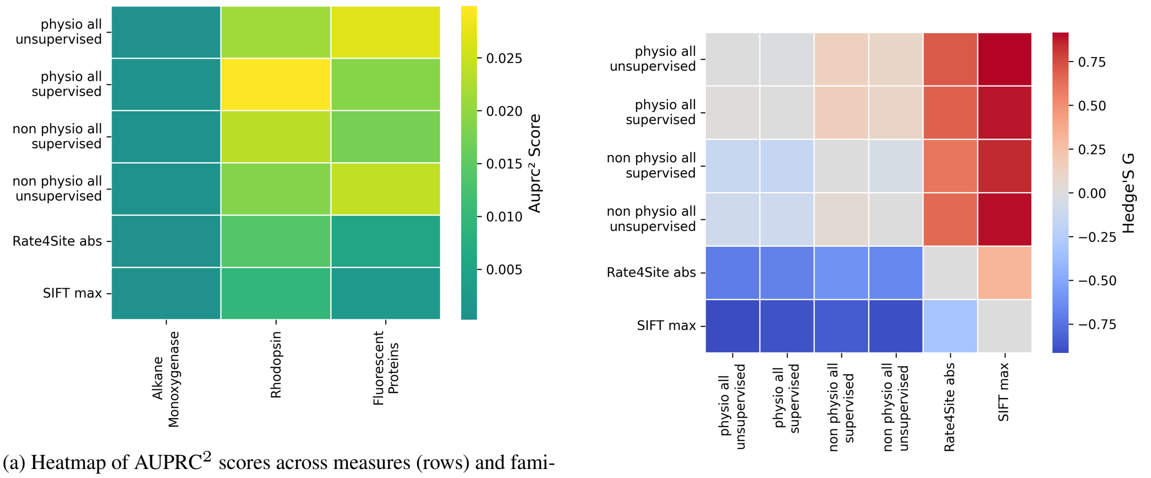


(a) Heatmap of AUPRC$^2$ scores across measures (rows) and families (columns).

(b) Heatmap of pairwise Hedges' $g$ effect sizes between measures.

Figure 4: **Heatmaps: Scores and Effect Sizes (AUPRC$^2$).** The left panel visualizes raw AUPRC$^2$ scores; color intensity (blue = high, yellow = low) indicates performance levels. The right panel shows standardized effect sizes for all pairwise measure comparisons, with warm colors indicating large differences. These heatmaps enable rapid identification of top-performing measures and family-dependent variation.

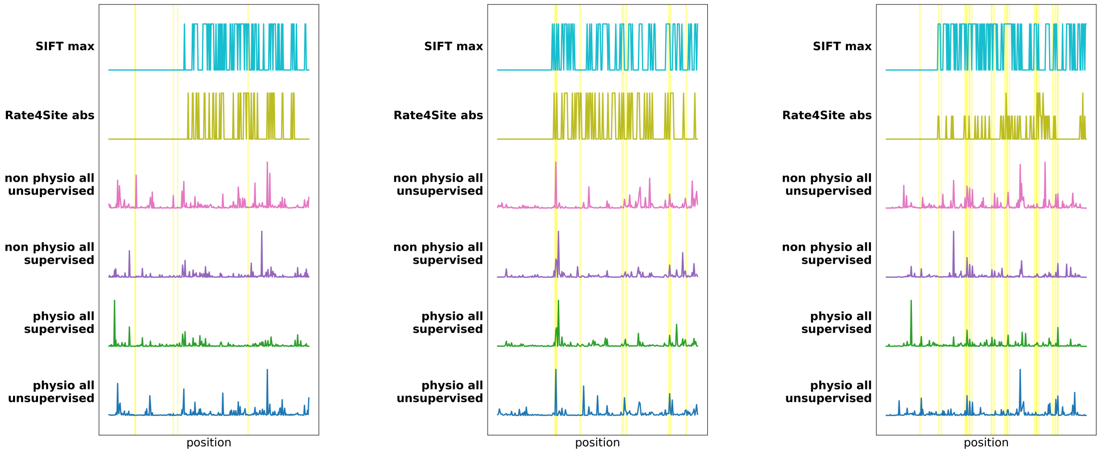


(a) AMO: Stacked importance scores at key functional positions.

(b) GFP: Stacked importance scores showing consistency across top 3 measures.

(c) Rhod: Stacked importance for key residues in bacteriorhodopsin.

Figure 5: **Positional Importance Profiles (AUPRC$^2$ Metric, Stacked by Family).** Each panel shows the combined importance scores assigned by the top 3 measures to literature-identified key residues. Stacking reveals agreement (tall bars = consensus) or disagreement (short bars = divergence) among measures. Strong consensus at known functional positions indicates high-quality predictors.

#### 4.2 Wasserstein Concentration Score (WCS) — Combined Top 3 Measures

This section repeats the analysis framework using the Wasserstein Concentration Score (WCS), which emphasizes concentrated, spatially-aligned predictions. WCS is particularly sensitive to how well importance is localized near functional regions.

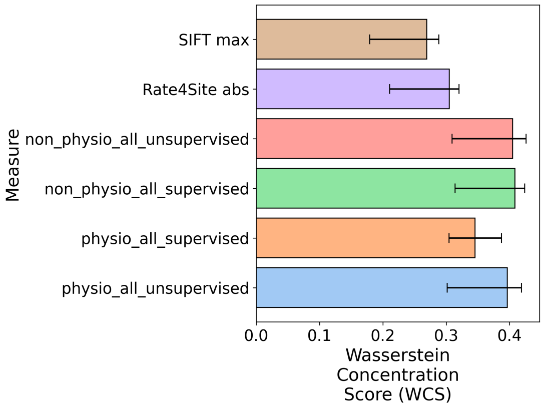


(a) Coherence scores using WCS metric.

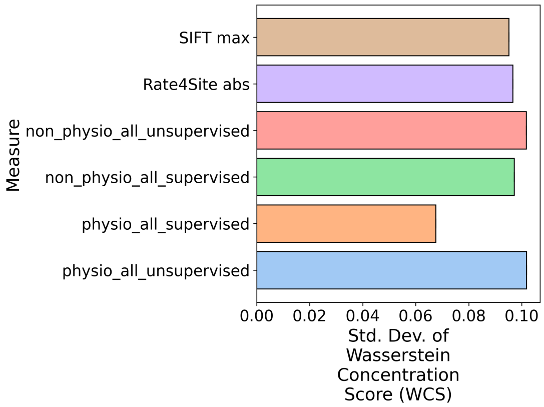


(b) Stability scores reflecting WCS robustness.

Figure 6: **Coherence and Stability Analysis (WCS Metric).** These bars mirror the $\text{AUPRC}^2$ analysis but now employ the Wasserstein Concentration Score, which rewards both high importance at key residues and concentration of scores. WCS typically shows higher stability when measures produce concentrated predictions.

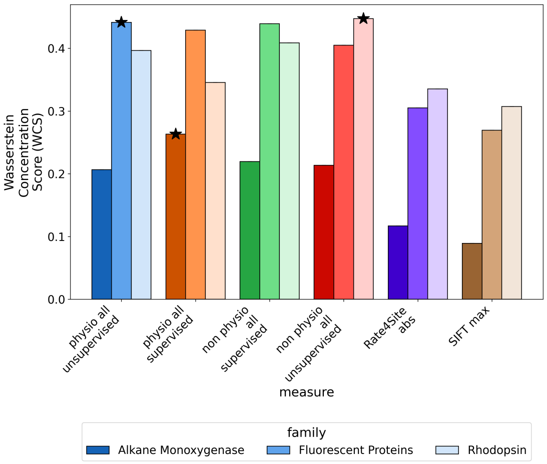


(a) Comparative WCS performance across families.

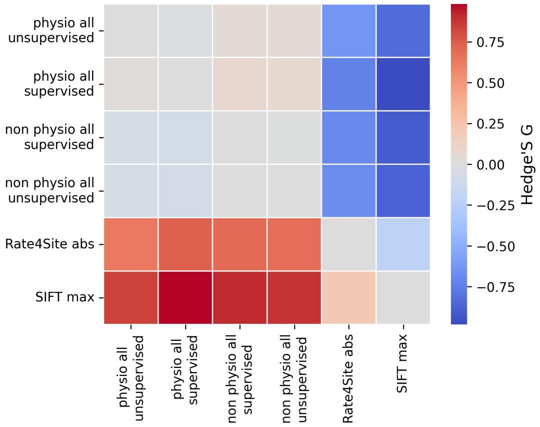


(b) Hedges' $g$ effect sizes for WCS pairwise comparisons.

Figure 7: **Performance and Effect Sizes (WCS Metric).** Panel (a) shows WCS scores for top 3 measures across families. Panel (b) displays standardized effect sizes. Comparison of AUPRC[2] and WCS results reveals which measures exhibit concentration properties favored by the Wasserstein metric.

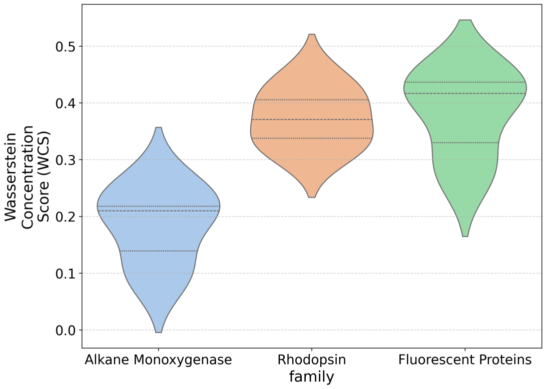


(a) Distribution of WCS coherence scores.

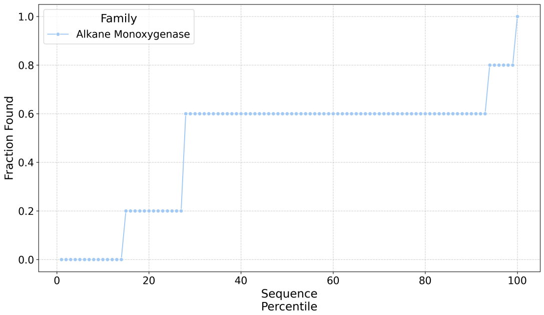


(b) Literature recovery curve: cumulative key residue detection via WCS ranking.

Figure 8: **Distribution and Literature Recovery (WCS Metric).** Visualization of WCS-based coherence distributions and cumulative recovery curves. Steeper recovery curves under WCS indicate that concentration properties align well with functional residue localization.

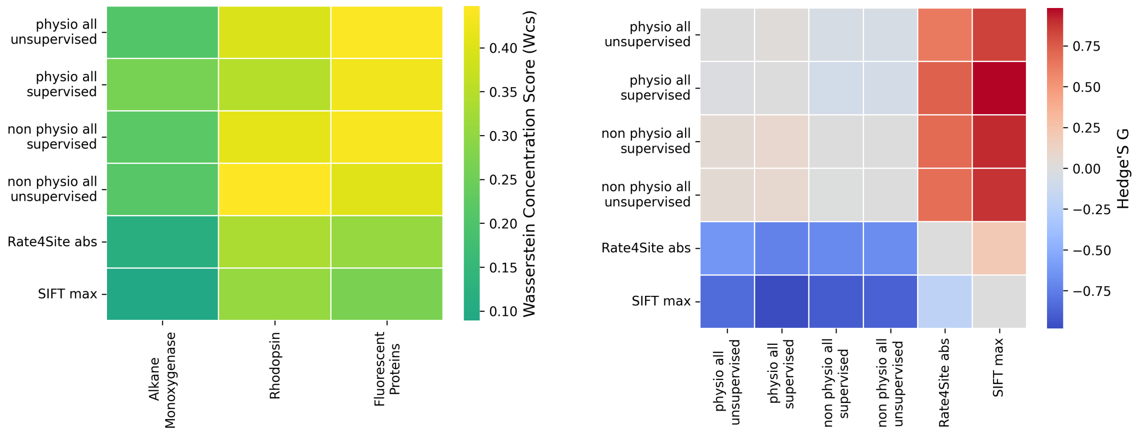


(a) Heatmap of WCS scores across measures and families.

(b) Heatmap of Hedges' $g$ effect sizes (WCS).

Figure 9: **Heatmaps: Scores and Effect Sizes (WCS).** Left panel visualizes WCS performance heatmap; right panel shows standardized effect sizes for pairwise measure comparisons under the WCS metric. Comparing these heatmaps with the AUPRC$^2$ results (Figure 4) reveals metric-dependent ranking shifts.

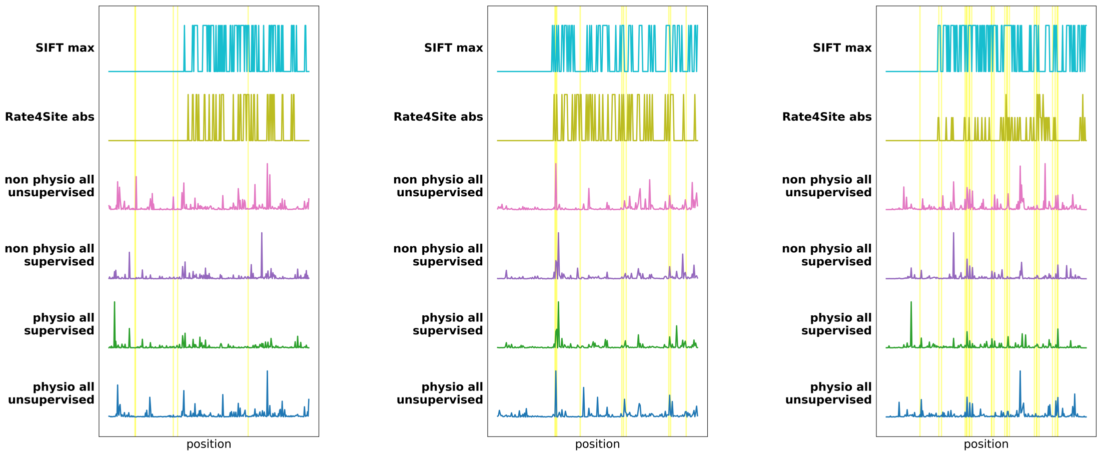


(a) AMO: Stacked WCS importance.

(b) GFP: Stacked WCS importance.

(c) Rhod: Stacked WCS importance.

Figure 10: **Positional Importance Profiles (WCS Metric, Stacked by Family).** Stacked bar charts showing combined importance scores from the top 3 measures under WCS. Taller stacks at functional residues and lower stacks at non-functional positions indicate good concentration and selectivity, key properties rewarded by the WCS metric.

#### 5 Interpretation and Conclusions

##### 5.1 Key Findings

1. **Metric Consistency**: Top-performing measures show good agreement between $AUPRC^2$ and WCS rankings (Figures 2a–7a), suggesting robust method selection.

2. **Family-Specific Variation**: Coherence and stability vary substantially across protein families (visible in heatmaps, Figures 4 and 9). This may reflect differences in functional residue distribution and alignment quality.

3. **Concentration vs. Discrimination**: WCS scores often differ from $AUPRC^2$ results, indicating that concentration (few residues with high importance) and discrimination (correct residue ranking) are complementary properties not always optimized by the same measure.

4. **Encoding Comparison**: Comparison of stacked residue plots (Figures 5 and 10) shows that standard and physicochemical encodings produce qualitatively similar prediction patterns, with encoding-specific variations at marginal residues.

##### 5.2 Recommendations for Method Selection

For protein engineering applications requiring functional residue prediction:

- Use **$AUPRC^2$** when discrimination among residues is paramount (e.g., targeted mutagenesis).

- Use **WCS** when prediction confidence and concentration matter (e.g., designed motif identification).
- Employ **ensemble approaches** combining top measures from both metrics to hedge against metric-specific biases.
