## Supplementary material for "Disentangling Protein Function via Decoupled Information Theoretic Selection of Key Tuning Residues": S5: Supplementary Methods

### Supplementary Methods and Proofs: *Decoupled Information Theoretic Feature Selection for Rapid Protein Key Tuning Residue Identification*

Haris Saeed[1,*], Aidong Yang[1],
Wei E. Huang[1,*]

[1]Department of Engineering Science, University of Oxford, Oxford, OX1 3PJ, United Kingdom

### Supplementary Methods and Proofs

This supplementary document provides detailed methodological descriptions and mathematical proofs related to the main paper. The following sections are included:

- Section 1: Detailed Methodology
- Section 2: Mathematical Proofs
- Section 3: References

### S1 Detailed Methodology

#### S1.1 Utilization of Existing Tools

To comprehensively evaluate our approach, we employed five established computational tools—HotSpot Wizard, EVcouplings, DeepSequence, SIFT, and Rate4Site—each leveraging distinct methodologies to identify functionally critical residues. These tools collectively span structural modeling, evolutionary coupling analysis, deep generative models, sequence conservation, and mutation effect prediction, providing a robust benchmark for assessing our information-theoretic framework.

HotSpot Wizard integrates structural and evolutionary data to predict mutational hotspots; EVcouplings identifies co-evolutionary couplings between residue positions; DeepSequence employs variational autoencoders to model sequence-function relationships; SIFT (Sorting Intolerant From Tolerant) predicts mutation tolerance based on sequence homology; and Rate4Site quantifies position-specific evolutionary rates from phylogenetic analysis. By benchmarking against these complementary methodologies, we systematically assessed our tool's capability to identify residues responsible for protein function across diverse biophysical contexts. This section provides detailed descriptions of each tool's methodology, including input requirements, computational procedures, and output interpretation.

##### S1.1.1 HotSpot Wizard

HotSpot Wizard 3.0 [?] is an online tool that combines structural, functional, and evolutionary data to predict mutational hotspots in proteins. These hotspots are residues that, when mutated, are most likely to affect the protein's functionality, stability, or interactions. The tool integrates information from structural biology, sequence data, and evolutionary conservation, allowing for a comprehensive prediction of key residues.

**Input Data:** The first step in the HotSpot Wizard workflow involves providing the protein sequence data. This is typically accompanied by a structural model of the protein, which can be generated through various methods such as X-ray crystallography, NMR spectroscopy, or computational prediction tools like AlphaFold. In our study, we used AlphaFold to generate high-confidence structure models for the input sequences [?]. The structural models are then uploaded to HotSpot Wizard to be analyzed.

**Analysis and Scoring:** HotSpot Wizard analyzes each residue in the context of its position within the protein structure and its evolutionary conservation across related protein sequences. It then assigns a mutability score to each residue, which reflects the likelihood that a mutation will have a functional impact. The residues are categorized into four distinct groups based on their predicted function in the protein's active site, tunnels, or catalytic pocket: - **Mutability:** A measure of how likely a residue is to tolerate mutations. - **Non-essentiality:** Indicates whether a residue is non-essential for the protein's core functions. - **In-tunnel:** Residues located in channels or tunnels critical for substrate access or product release. - **In-catalytic pocket:** Residues within the active site that are directly involved in catalytic activity.

These scores are normalized and summed to produce a total score out of 4, indicating the residue's overall functional importance. The final output from HotSpot Wizard provides a list of residues likely to be functionally important, along with their predicted mutability and categorization.

**Advantages and Limitations:** The main advantage of HotSpot Wizard is its ability to predict functionally important residues without requiring experimental mutagenesis data. However, its accuracy is highly dependent on the quality of the structural model provided. Incorrect or low-confidence models can lead to inaccurate hotspot predictions. Therefore, using high-confidence models, such as those provided by AlphaFold, significantly improves the reliability of the predictions.

##### S1.1.2 EVcouplings

EVcouplings [?, ?] is a tool used to detect evolutionary couplings, which are correlations between mutations in residues that indicate structural or functional linkage. This method leverages evolutionary data to identify pairs of residues that are likely to co-evolve, providing insights into functionally important interactions within the protein.

**Input Data:** The first step in the EVcouplings workflow is to generate a high-quality multiple sequence alignment (MSA) of homologous protein sequences. We used the Kalign alignment tool [?] due to its efficiency and high alignment accuracy, especially when dealing with large datasets. The input sequences are aligned based on their evolutionary relationships, which is critical for accurately detecting evolutionary couplings.

**Coupling Calculation:** The evolutionary couplings are derived from the MSA by analyzing the co-variation of residues across different sequences. This process is done through the calculation of mutual information between pairs of residues. The resulting 4D cross-correlation matrix is then flattened into a 1D vector, with each element corresponding to a potential evolutionary coupling. We considered both the **minimum** and **maximum** coupling values to identify the most significant evolutionary constraints between residue pairs. These couplings provide insight into functionally important residues that are crucial for maintaining the protein's stability and function.

**Post-Processing:** After calculating the evolutionary couplings, we identified residue pairs with the highest mutual information as critical for maintaining the protein structure and function. The coupling data was then used to guide the identification of co-evolved residues, which are often involved in maintaining protein structure or catalysis. Residues with strong evolutionary couplings were categorized as likely to have functional significance, particularly in maintaining protein integrity during mutations.

**Advantages and Limitations:** One of the main advantages of EVcouplings is its ability to identify residues involved in long-range interactions, even in the absence of detailed structural data. However, the method is highly dependent on the quality and diversity of the input MSA. If the alignment is poor or lacks enough homologous sequences, the accuracy of the coupling predictions may decrease. Additionally, evolutionary coupling can sometimes miss important local interactions if they do not exhibit significant co-evolutionary signals.

###### S1.1.3 DeepSequence

DeepSequence [?] utilizes deep generative models, specifically variational autoencoders, to predict the effects of mutations on protein function. This method leverages a large-scale analysis of mutational data to model the sequence-function relationship, providing predictions for how mutations at specific residues can alter protein activity.

**Input Data:** DeepSequence requires the protein sequence as input, and the tool works by generating multiple sequence alignments (MSAs) using Kalign [?], ensuring high-quality alignments that are necessary for accurate predictions. The sequence data is used to train the deep generative model, which learns a latent representation of the sequence-function relationship. This model captures the distribution of functional protein sequences, allowing for the prediction of the functional impact of single-point mutations.

**Latent Representation:** The primary output of DeepSequence is a set of latent representations that capture the underlying sequence-function relationship for the input sequences. These representations are used to predict the effect of individual mutations on protein function. The method ranks the mutations based on the evidence lower bound (ELBO), with **Max(ELBO)**, **Min(ELBO)**, and **Max(Abs(ELBO))** serving as criteria for identifying mutations likely to alter protein function significantly.

**Prediction of Functional Shifts:** DeepSequence evaluates the functional impact of mutations by comparing the likelihood of the mutated sequence to the distribution of functional sequences learned by the model. This allows the tool to identify residues whose mutations lead to significant functional shifts. The ranking of mutations is based on how far the mutation deviates from the learned distribution of functional sequences, providing a comprehensive view of the mutation's impact on protein activity.

**Advantages and Limitations:** The main advantage of DeepSequence is its ability to predict the effects of a wide range of mutations, including those that might not be directly involved in active sites or structural elements. However, DeepSequence's reliance on large-scale data and deep learning models means it requires substantial computational resources and high-quality training data to generate reliable predictions. Additionally, the tool may struggle with novel mutations not present in the training dataset.

###### S1.1.4 SIFT (Sorting Intolerant From Tolerant)

SIFT [?] is a sequence-based tool that predicts whether an amino acid substitution affects protein function by analyzing sequence conservation across homologous proteins. The method is predicated on the principle that evolutionarily conserved positions are functionally important and less tolerant to substitutions.

**Input Data and Alignment:** SIFT requires a protein sequence as input and automatically generates a multiple sequence alignment (MSA) of homologous sequences from protein databases (e.g., UniProt, SwissProt). The quality and diversity of the MSA critically determine prediction accuracy. SIFT employs PSI-BLAST to identify homologous sequences with appropriate sequence identity thresholds (typically 90% to avoid over-representation and ensure sufficient diversity).

**Scoring Methodology:** For each position in the query sequence, SIFT calculates the probability distribution of amino acids observed in the MSA. The SIFT score for a specific substitution is derived from the normalized probability of observing that particular amino acid at that position:

$$\text{SIFT Score} = \frac{P(\text{substituted amino acid})}{\sum_{i=1}^{20} P(\text{amino acid}_i)} \tag{S1}$$

where $P(\text{amino acid}_i)$ represents the observed probability of the $i$-th amino acid at the position of interest. Scores range from 0 to 1, with lower scores indicating greater conservation and higher predicted functional impact. A threshold of 0.05 is typically used: substitutions with scores $\leq 0.05$ are predicted to be deleterious (intolerant), while scores $> 0.05$ suggest neutral or tolerated mutations.

**Advantages and Limitations:** SIFT's primary advantage is its computational efficiency and interpretability based on evolutionary conservation principles. However, it may fail to detect functionally important residues in rapidly evolving regions or when homologous sequences are scarce. Additionally, SIFT does not account for structural context or compensatory mutations.

###### S1.1.5 Rate4Site

Rate4Site [?] estimates position-specific evolutionary rates using phylogenetic methods, providing a quantitative measure of selective constraints acting on each residue. Slowly evolving positions are inferred to be functionally or structurally important, as they experience stronger purifying selection.

**Input Data and Phylogenetic Analysis:** Rate4Site requires a multiple sequence alignment and a phylogenetic tree relating the sequences. The phylogenetic tree can be provided by the user or constructed using maximum likelihood or neighbor-joining methods. The tool implements an empirical Bayesian approach to estimate site-specific evolutionary rates under a given substitution model (e.g., JTT, WAG for proteins).

**Rate Calculation:** For each alignment position, Rate4Site calculates the relative evolutionary rate $r_i$ by maximum likelihood estimation:

$$r_i = \frac{\text{Substitution rate at position } i}{\text{Average substitution rate across all positions}} \tag{S2}$$

The rates are normalized such that the mean rate across all sites equals 1. Lower rates ($r_i < 1$) indicate slower evolution and higher functional importance, while higher rates ($r_i > 1$) suggest faster evolution and lower selective constraint. Rate4Site also provides confidence intervals for rate estimates and identifies significantly conserved positions based on statistical thresholds.

**Output Interpretation:** Rate4Site outputs include position-specific evolutionary rates, confidence scores, and color-coded conservation grades. These can be mapped onto protein structures to visualize functional regions. The Bayesian framework accounts for phylogenetic uncertainty and provides robust statistical inference.

**Advantages and Limitations:** Rate4Site's strength lies in its rigorous statistical framework and phylogenetic context, making it particularly effective for detecting subtle conservation patterns. However, it requires high-quality alignments and phylogenetic trees, and computational demands increase with sequence number and alignment length. Additionally, the method assumes tree topology is correct and that the substitution model adequately describes evolutionary processes.

#### S1.2 Amino Acid Feature Encoding and Physicochemical Representation

Each of the 20 canonical amino acids is represented by a 16-dimensional feature vector encoding key physicochemical descriptors. These properties provide a comprehensive biochemical characterization of each amino acid, enabling sophisticated analysis of sequence properties beyond simple categorical assignments. The feature vector captures both continuous properties reflecting molecular characteristics and binary categorical properties indicating membership in functionally relevant groups.

**Continuous Properties:**

The continuous component of the feature vector encodes quantitative molecular properties that directly influence protein structure and function:

- **Molecular weight** (g/mol): The sum of atomic masses, ranging from 75 (G) to 204 (W). Influences protein density, diffusion rate, and steric constraints.
- **Charge at pH 7.4** (elementary charges): Ionizable groups determine electrostatic interactions. Ranges from -2 (D, E) to +2 (K, R, H). Computed from pKa values of side chain functional groups.
- **Isoelectric point (pI)**: The pH at which the amino acid carries no net charge. Ranges from 2.8 (D, E) to 10.8 (R). Relevant for protein-protein interactions and cellular localization.
- **Hydropathy index (Kyte-Doolittle scale)**: Quantifies hydrophobic character on a scale typically ranging from -4.5 (E, K) to +4.5 (I, L, F, W). Critical for predicting transmembrane regions and protein-protein interfaces.
- **Volume (Å$^3$)**: The van der Waals volume of the side chain, ranging from 48 (G) to 227 (W). Affects packing density and steric constraints in protein structures.
- **Bulkiness (Zimmerman)**: A measure of how much space the side chain occupies relative to its mass. Important for secondary structure propensities and folding kinetics.
- **Flexibility (Vihinen)**: Quantifies conformational variability of the side chain. High flexibility (S, T, P) enables loop formation; low flexibility (F, W) constrains backbone geometry.
- **Hydrogen bond donors (sidechain)**: Number of polar groups capable of donating hydrogen bonds in the side chain. Ranges from 0 (A, G, L, V, I, P, F) to 4 (Y can participate as both donor and acceptor).
- **Hydrogen bond acceptors (sidechain)**: Number of polar groups capable of accepting hydrogen bonds. Ranges from 0 (G, A, V, I, L, P) to 4 (D, E, S, T, N, Q, Y).

**Binary Categorical Properties:**

The categorical component uses indicator variables (0/1) to denote membership in functionally relevant amino acid groups:

- **Is aliphatic**: Residues A, G, I, L, V. These amino acids form hydrophobic cores and are enriched in structural regions like $\alpha$-helices and $\beta$-sheets.
- **Is aromatic**: Residues F, W, Y. Aromatic rings enable $\pi$-$\pi$ stacking interactions, cation-$\pi$ interactions, and often participate in binding pockets and electron transfer.
- **Is polar**: Residues N, Q, S, T. Polar residues support hydrogen bonding networks crucial for protein stability and specificity in enzyme catalysis.
- **Contains sulfur**: Residues C, M. Sulfur enables disulfide bond formation (C), metal coordination (M), or soft nucleophile chemistry.
- **Is acidic**: Residues D, E. Acidic residues contribute negative charge, form salt bridges, and participate in proton transfer pathways.
- **Is basic**: Residues H, K, R. Basic residues contribute positive charge, form salt bridges, and coordinate anionic cofactors.
- **Is cyclic**: Residue P. Proline's constrained backbone conformation restricts its role in helical structures but enables turns and loops.

**Feature Table:**
Table **??** presents the complete 16-dimensional feature vectors for all 20 canonical amino acids. The continuous features (columns 1–9) are presented in standard units; the categorical features (columns 10–16) are binary indicators. This comprehensive encoding enables:

1. **Biochemically-informed distance metrics**: Computing distances between residues based on shared biochemical properties rather than arbitrary symbolic differences.
2. **Clustering of similar residues**: K-means clustering applied to these vectors produces functionally meaningful amino acid groupings specific to each alignment column.
3. **Supervised feature learning**: Machine learning models can leverage these properties to identify residues whose physicochemical characteristics correlate with functional phenotypes.
4. **Interpretable predictions**: Identified residues and their contributions can be explained in terms of specific biochemical properties.

| AA | MW | Charge | pI | Hydro | Vol | Bulk | Flex | HBD | HBA | Aliph | Arom | Polar | Su |
|---|---|---|---|---|---|---|---|---|---|---|---|---|---|
| A | 89 | 0.0 | 6.0 | 1.8 | 88 | 0.46 | 0.58 | 0 | 0 | 1 | 0 | 0 | [illegible] |
| C | 121 | -0.2 | 5.1 | 2.5 | 108 | 0.54 | 0.70 | 1 | 1 | 0 | 0 | 0 | [illegible] |
| D | 133 | -0.9 | 2.8 | -3.5 | 111 | 0.54 | 0.54 | 0 | 2 | 0 | 0 | 0 | [illegible] |
| E | 147 | -0.9 | 3.2 | -3.5 | 138 | 0.68 | 0.62 | 0 | 2 | 0 | 0 | 0 | [illegible] |
| F | 165 | 0.0 | 5.5 | 2.8 | 189 | 0.95 | 0.61 | 0 | 0 | 0 | 1 | 0 | [illegible] |
| G | 75 | 0.0 | 6.0 | -0.4 | 60 | 0.00 | 0.51 | 0 | 0 | 1 | 0 | 0 | [illegible] |
| H | 155 | 0.1 | 7.6 | -0.4 | 153 | 0.77 | 0.61 | 1 | 2 | 0 | 1 | 0 | [illegible] |
| I | 131 | 0.0 | 6.0 | 4.5 | 124 | 0.62 | 0.60 | 0 | 0 | 1 | 0 | 0 | [illegible] |
| K | 146 | 1.0 | 10.5 | -3.9 | 135 | 0.68 | 0.95 | 1 | 1 | 0 | 0 | 0 | [illegible] |
| L | 131 | 0.0 | 6.0 | 3.8 | 124 | 0.62 | 0.59 | 0 | 0 | 1 | 0 | 0 | [illegible] |
| M | 149 | 0.0 | 5.7 | 1.9 | 124 | 0.62 | 0.70 | 0 | 1 | 0 | 0 | 0 | [illegible] |
| N | 132 | 0.0 | 5.4 | -3.5 | 114 | 0.57 | 0.67 | 1 | 2 | 0 | 0 | 1 | [illegible] |
| P | 115 | 0.0 | 6.3 | -1.6 | 98 | 0.49 | 0.51 | 0 | 1 | 0 | 0 | 0 | [illegible] |
| Q | 146 | 0.0 | 5.7 | -3.5 | 144 | 0.72 | 0.80 | 1 | 2 | 0 | 0 | 1 | [illegible] |
| R | 174 | 1.0 | 10.8 | -4.5 | 173 | 0.87 | 1.08 | 4 | 2 | 0 | 0 | 0 | [illegible] |
| S | 105 | 0.0 | 5.7 | -0.8 | 89 | 0.45 | 0.64 | 1 | 2 | 0 | 0 | 1 | [illegible] |
| T | 119 | 0.0 | 5.6 | -0.7 | 102 | 0.51 | 0.68 | 1 | 2 | 0 | 0 | 1 | [illegible] |
| V | 117 | 0.0 | 6.0 | 4.2 | 106 | 0.53 | 0.53 | 0 | 0 | 1 | 0 | 0 | [illegible] |
| W | 204 | 0.0 | 5.9 | -0.9 | 227 | 1.14 | 0.75 | 1 | 1 | 0 | 1 | 0 | [illegible] |
| Y | 181 | 0.0 | 5.7 | -1.3 | 193 | 0.97 | 0.72 | 1 | 2 | 0 | 1 | 1 | [illegible] |

Table S1: **16-dimensional feature vectors for canonical amino acids.** Each amino acid is characterized by 9 continuous properties and 7 binary categorical properties. **Continuous properties**: MW = Molecular Weight (g/mol); Charge = Charge at pH 7.4 (elementary charges); pI = Isoelectric point; Hydro = Hydropathy index (Kyte-Doolittle); Vol = Van der Waals volume ($Å^3$); Bulk = Bulkiness (Zimmerman normalized); Flex = Flexibility (Vihinen normalized); HBD = Hydrogen bond donors; HBA = Hydrogen bond acceptors. **Binary categorical properties**: Aliph = Is aliphatic; Arom = Is aromatic; Polar = Is polar; Sulfur = Contains sulfur; Acid = Is acidic; Basic = Is basic; Cyclic = Is cyclic (proline). Binary features enable rapid grouping of biochemically similar residues, while continuous features support distance-based clustering and machine learning applications. Normalized continuous features are scaled to unit variance for consistency.

**Normalization and Standardization:**

Before using these features in computational analysis, continuous properties are standardized (zero mean, unit variance) independently for each feature dimension. This normalization ensures that features with large natural scales (e.g., molecular weight) do not dominate clustering or distance computations. The standardized features are then used for:

- **K-means clustering**: Applied independently to each alignment column to group biochemically similar residues present in that column.
- **Distance calculations**: Computing Euclidean distances between residues as proxies for biochemical substitution cost.
- **Machine learning**: Serving as input features for supervised models (Lasso, Random Forest, Bayesian regression) in labeled sequence-property prediction tasks.

#### S1.3 Mutation Effect Prediction and Scoring Strategies

For single-mutation effect predictors (DeepSequence, SIFT), we systematically conducted *in silico* mutagenesis by evaluating all possible single amino acid substitutions at each position in the query sequence. This exhaustive approach generated 19 mutational variants per position (20 amino acids minus the wild-type residue), yielding comprehensive positional mutational landscapes.

**Scoring Aggregation:** Each mutational scan produced a distribution of effect scores per position. To derive a single importance score per residue from these distributions, we evaluated six pooling strategies that capture different aspects of the mutational effect landscape:

1. **Minimum (Min):** The most deleterious mutation effect at each position

$$S_{\text{min}}(i) = \min_{a \in \mathcal{A} \setminus \{a_{\text{wt}}\}} \Delta E(i, a)$$

This identifies positions where at least one mutation is highly detrimental, indicating critical functional residues.

2. **Maximum (Max):** The least deleterious or most beneficial mutation effect

$$S_{\text{max}}(i) = \max_{a \in \mathcal{A} \setminus \{a_{\text{wt}}\}} \Delta E(i, a)$$

This reveals positions with high mutational robustness or potential for functional improvement.

3. **Mean:** The average mutation effect across all substitutions

$$S_{\text{mean}}(i) = \frac{1}{|\mathcal{A}| - 1} \sum_{a \in \mathcal{A} \setminus \{a_{\text{wt}}\}} \Delta E(i, a)$$

This provides an overall measure of positional mutational tolerance.

4. **Minimum Absolute (Min Abs):** The smallest absolute effect magnitude

$$S_{\text{min\_abs}}(i) = \min_{a \in \mathcal{A} \setminus \{a_{\text{wt}}\}} |\Delta E(i, a)|$$

This identifies the most conservative substitution at each position.

5. **Maximum Absolute (Max Abs):** The largest absolute effect magnitude

$$S_{\text{max\_abs}}(i) = \max_{a \in \mathcal{A} \setminus \{a_{\text{wt}}\}} |\Delta E(i, a)|$$

This captures positions with the most extreme mutational sensitivity, regardless of direction.

6. **Mean Absolute (Mean Abs):** The average absolute effect magnitude

$$S_{\text{mean\_abs}}(i) = \frac{1}{|\mathcal{A}| - 1} \sum_{a \in \mathcal{A} \setminus \{a_{\text{wt}}\}} |\Delta E(i, a)|$$

This quantifies the overall mutational sensitivity at each position, irrespective of effect direction.

where $\mathcal{A}$ denotes the set of 20 standard amino acids, $a_{\text{wt}}$ is the wild-type residue at position $i$, and $\Delta E(i, a)$ represents the predicted functional effect of substituting the wild-type residue with amino acid $a$.

**Rationale:** These diverse pooling strategies enable comprehensive characterization of positional functional importance. Minimum-based metrics (Min, Min Abs) identify functionally critical positions that cannot tolerate specific mutations, while maximum-based metrics (Max, Max Abs) reveal mutational robustness or plasticity. Mean-based metrics (Mean, Mean Abs) provide aggregate measures of mutational tolerance. By comparing rankings derived from different pooling strategies, we assessed the consistency and complementarity of various importance metrics, ultimately selecting those that best correlated with experimental validation data.

#### S1.4 Entropic Measures and Mutual Information

#### S1.5 Entropic Measures and Mutual Information

Information theory provides a rich mathematical framework for quantifying uncertainty, disorder, and dependencies in complex systems. In the context of protein sequence analysis, entropic measures capture the degree of conservation or variability at each position, while mutual information quantifies dependencies between positions or between sequence features and functional properties. We employed a comprehensive suite of entropic measures, each offering distinct sensitivity to distribution properties and theoretical foundations:

- **Shannon Entropy** [?]: The foundational measure of uncertainty in probability distributions, quantifying the average information content per symbol.
- **Rényi Entropy** [?]: A parameterized generalization of Shannon entropy that adjusts sensitivity to rare events through the order parameter $\alpha$, enabling exploration of distribution tail behavior.
- **Tsallis Entropy** [?]: A non-additive entropy formulation suited for non-extensive statistical systems, particularly relevant for long-range correlations in protein sequences.
- **Kullback-Leibler Divergence** [?]: A directed measure of distributional divergence from a reference distribution, quantifying information loss when approximating one distribution with another.
- **von Neumann Entropy** [?]: The quantum analog of Shannon entropy, applicable to density matrices and quantum information contexts, though primarily included for theoretical completeness.
- **Gini Impurity** [?]: A measure of statistical dispersion originally developed for inequality quantification, capturing the probability of misclassification in categorical distributions.
- **Kolmogorov Complexity** [?]: A measure of algorithmic information content defined as the length of the shortest program generating a sequence, providing a compression-based complexity metric.

- **Jensen-Shannon Divergence** [?]: A symmetrized and smoothed version of KL divergence, providing a proper distance metric between probability distributions with bounded values.
- **Average Entropy**: The arithmetic mean of positional entropies across an alignment, providing a global measure of sequence diversity.
- **Hard Voting**: An ensemble approach where each entropy measure contributes equally to residue importance rankings, implementing majority consensus.
- **Soft Voting**: A weighted ensemble approach where measures contribute according to their individual predictive performance or confidence scores.

By systematically comparing these diverse entropic measures, we assessed the robustness of our findings across different theoretical frameworks and identified which measures best capture functionally relevant sequence variability. This comprehensive approach ensures that our conclusions are not artifacts of any single information-theoretic formulation.

##### S1.5.1 Shannon Entropy

Shannon Entropy, as introduced by Claude Shannon [?], is a fundamental concept in information theory that measures the uncertainty or randomness in a system. In the context of protein sequences, it quantifies the unpredictability of the amino acid at each position in a multiple sequence alignment (MSA), providing insight into evolutionary conservation. Entropy is based on the idea that the more evenly distributed the possibilities are, the greater the uncertainty, while fewer possibilities (such as a single dominant amino acid) imply a lower degree of uncertainty.

The Shannon Entropy $H(i)$ at position $i$ in an MSA can be mathematically defined as:

$$H(i) = -\sum_{a \in A} p(a) \log p(a) \tag{S3}$$

where: - $A$ represents the alphabet of possible amino acids (or residues) at position $i$ in the MSA, typically consisting of 20 amino acids in protein sequences, - $p(a)$ is the probability of amino acid $a$ occurring at position $i$, calculated by the relative frequency of $a$ in the sequence alignment.

The entropy value $H(i)$ can range from 0 to $\log |A|$, where $|A|$ is the size of the alphabet (in this case, 20 for amino acids). The maximum entropy of $\log |A|$ occurs when all amino acids are equally probable, which implies maximum unpredictability and no conserved residues at that position. On the other hand, if $H(i) = 0$, it means that the residue at that position is perfectly conserved across all sequences in the MSA, indicating functional or structural importance.

The relationship between entropy and conservation is fundamental: - **Low entropy** (close to 0) indicates a high degree of conservation at position $i$, suggesting the residue at that position is critical for maintaining the protein's structure or function. - **High entropy** suggests high variability, which is typically observed in regions subject to evolutionary pressure for functional flexibility, such as interaction surfaces or signaling regions.

To ensure comparability across different protein families and MSAs, we scale the entropy values for each alignment such that the maximum entropy within a given dataset is normalized to 1:

$$H_{\text{scaled}}(i) = \frac{H(i)}{\max(H)} \tag{S4}$$

This normalization allows for comparisons across proteins with varying sequence diversity, ensuring that the entropy values reflect relative conservation irrespective of dataset size or intrinsic variability.

##### S1.5.2 Mutual Information

Mutual Information (MI), also based on information theory [?], measures the amount of information shared between two variables, quantifying the reduction in uncertainty about one variable given the knowledge of the other. In the context of protein sequences, MI is used to assess the dependency between residue positions in a sequence and an associated functional or structural characteristic.

For two discrete random variables $X$ (residue position $i$ in a protein sequence) and $Y$ (a target variable such as a functional or stability score), the Mutual Information $I(X;Y)$ is defined as:

$$I(X;Y) = \sum_{x\in X}\sum_{y\in Y} p(x,y) \log\left(\frac{p(x,y)}{p(x)p(y)}\right) \tag{S5}$$

where: - $p(x,y)$ is the joint probability of observing a residue $x$ at position $i$ and a corresponding value $y$ for the target variable $Y$, - $p(x)$ and $p(y)$ are the marginal probabilities of $x$ and $y$, respectively, representing the individual distributions of residue positions and the target variable.

The MI measures the reduction in uncertainty about one variable ($X$) given the knowledge of the other variable ($Y$). If $X$ and $Y$ are independent, the MI value is 0, meaning that knowing one variable provides no information about the other. Conversely, higher MI values indicate a strong dependency between the residue at position $i$ and the functional or structural characteristic $Y$.

To calculate MI in the context of continuous target variables, such as a quantitative functional score, we must discretize the continuous variable $Y$ into a set of bins. We applied the Freedman-Diaconis rule to determine the optimal bin width for discretization, ensuring that the bins capture the distribution of the target variable without introducing excessive bias:

$$\text{Bin width} = 2 \cdot \frac{\text{IQR}(Y)}{\sqrt[3]{n}} \tag{S6}$$

where: - $\text{IQR}(Y)$ is the interquartile range of the target variable $Y$, - $n$ is the number of data points.

The MI between a residue position $i$ and the discretized target variable $T$ is then calculated as:

$$I(i;T) = H(i) + H(T) - H(i,T) \tag{S7}$$

where:

$$H(i) = -\sum_a p(a) \log p(a)$$

is the entropy of residue positions $i$,

$$H(T) = -\sum_t p(t) \log p(t)$$

is the entropy of the target variable $T$, and

$$H(i, T) = -\sum_a \sum_t p(a, t) \log p(a, t)$$

is the joint entropy of residue position $i$ and the target variable $T$.

In this formulation: - $H(i)$ measures the uncertainty of the amino acid at position $i$, - $H(T)$ quantifies the uncertainty in the target variable $T$, - $H(i, T)$ quantifies the combined uncertainty between the residue at position $i$ and the target variable $T$.

Normalization of the MI values is performed to ensure comparability across different datasets. Specifically, the MI values are scaled such that the maximum MI value is 1, allowing for consistent ranking of residue positions in terms of their functional or structural significance:

$$I_{\text{scaled}}(i; T) = \frac{I(i; T)}{\max(I)} \tag{S8}$$

High MI values identify residues that exhibit significant dependency with the target variable $T$, indicating positions critical for maintaining protein structure or function. These residues are considered key in determining protein activity or stability and can be further explored for engineering or therapeutic purposes.

The application of diverse entropic measures (Shannon, Rényi, Tsallis, etc.) and mutual information forms the foundation of our methodology for identifying key residues involved in critical functions such as protein stability, binding affinity, and catalytic activity [?, ?]. By leveraging multiple information-theoretic formulations, we ensure robustness against model-specific biases and capture the full spectrum of functionally relevant sequence variability.

##### S1.5.3 Rényi Entropy

Rényi Entropy [?] is a one-parameter generalization of Shannon entropy that provides greater flexibility in quantifying distributional uncertainty. The parameter $\alpha$ (order) controls the measure's sensitivity to rare versus common events, enabling systematic exploration of distribution characteristics beyond the Shannon framework.

The Rényi entropy of order $\alpha$ for a discrete random variable $X$ with probability distribution $p(x)$ is defined as:

$$H_\alpha(X) = \frac{1}{1-\alpha} \log \left( \sum_x p(x)^\alpha \right) \tag{S9}$$

where $\alpha \geq 0$ and $\alpha \neq 1$. For $\alpha = 1$, Rényi entropy converges to Shannon entropy:

$$\lim_{\alpha \to 1} H_\alpha(X) = H(X) = -\sum_x p(x) \log p(x) \tag{S10}$$

**Interpretation of Parameter $\alpha$:**

- $\alpha = 0$: Hartley entropy, measuring the logarithm of the support size (number of non-zero probability events)
- $0 < \alpha < 1$: Emphasizes rare events; sensitive to distribution tails
- $\alpha = 1$: Shannon entropy (limit case)
- $\alpha > 1$: Emphasizes common events; less sensitive to rare occurrences
- $\alpha = 2$: Collision entropy, related to the probability that two independent samples yield the same outcome
- $\alpha \to \infty$: Min-entropy, measuring the negative logarithm of the maximum probability

**Application to Protein Sequences:**

For residue position $i$ in a multiple sequence alignment, we compute Rényi entropy across multiple orders $\alpha \in \{0.5, 1, 2, 5, \infty\}$ to capture different aspects of conservation:

$$H_\alpha(i) = \frac{1}{1-\alpha} \log \left( \sum_{a \in \mathcal{A}} p_i(a)^\alpha \right) \tag{S11}$$

where $\mathcal{A}$ is the alphabet of amino acids and $p_i(a)$ is the observed frequency of amino acid $a$ at position $i$.

Low-order Rényi entropies ($\alpha < 1$) are particularly useful for detecting positions with rare but functionally important mutations, while high-order entropies ($\alpha > 1$) effectively identify positions dominated by a few common amino acids, indicating strong selective constraints.

##### S1.5.4 Tsallis Entropy

Tsallis Entropy [?] is a non-additive generalization of Shannon entropy developed within the framework of non-extensive statistical mechanics. It is particularly relevant for systems exhibiting long-range correlations, memory effects, or fractal-like properties—characteristics potentially present in protein evolutionary dynamics.

The Tsallis entropy of order $q$ is defined as:

$$S_q(X) = \frac{1}{q-1} \left( 1 - \sum_x p(x)^q \right) \tag{S12}$$

where $q > 0$ is the entropic index. For $q = 1$, Tsallis entropy reduces to Shannon entropy:

$$\lim_{q \to 1} S_q(X) = -\sum_x p(x) \log p(x) = H(X) \tag{S13}$$

**Key Properties:**

- **Non-additivity**: For independent systems A and B, $S_q(A \cup B) = S_q(A) + S_q(B) + (1-q)S_q(A)S_q(B)$
- **Concavity**: $S_q$ is concave for $q > 0$
- **Extremality**: Maximum entropy occurs for uniform distributions

**Biological Relevance:**

The non-additive property of Tsallis entropy makes it suitable for analyzing protein sequences where residue positions exhibit dependencies (evolutionary couplings). Traditional Shannon entropy assumes independence, but protein evolution involves correlated mutations to maintain structural and functional integrity. Tsallis entropy with $q \neq 1$ can capture these non-extensive effects.

For a residue position $i$:

$$S_q(i) = \frac{1}{q-1}\left(1 - \sum_{a \in \mathcal{A}} p_i(a)^q\right) \tag{S14}$$

We typically explore $q \in \{0.5, 1.5, 2, 3\}$ to assess sensitivity to distribution shape. Values $q < 1$ emphasize rare events, while $q > 1$ emphasize common events, similar to Rényi entropy but with different mathematical properties and theoretical interpretations.

##### S1.5.5 Kullback-Leibler Divergence

The Kullback-Leibler (KL) Divergence [?], also known as relative entropy, quantifies how much one probability distribution diverges from a reference distribution. In protein sequence analysis, KL divergence measures the information lost when approximating the true amino acid distribution at a position with a reference (e.g., background) distribution.

For discrete probability distributions $P$ and $Q$ over the same alphabet, the KL divergence from $Q$ to $P$ is:

$$D_{KL}(P \| Q) = \sum_x P(x) \log \frac{P(x)}{Q(x)} \tag{S15}$$

**Properties:**

- **Non-negative**: $D_{KL}(P \| Q) \geq 0$ with equality if and only if $P = Q$
- **Asymmetric**: $D_{KL}(P \| Q) \neq D_{KL}(Q \| P)$ in general
- **Not a metric**: Does not satisfy triangle inequality
- **Unbounded**: Can approach infinity if $Q(x) = 0$ for some $x$ where $P(x) > 0$

**Application to Conservation Analysis:**

For each position $i$ in a multiple sequence alignment, we compute KL divergence from the observed amino acid distribution $P_i$ to a background distribution $Q_{\text{bg}}$ (typically the average amino acid frequencies across all positions or from a large protein database):

$$D_{KL}(P_i \| Q_{\text{bg}}) = \sum_{a \in \mathcal{A}} P_i(a) \log \frac{P_i(a)}{Q_{\text{bg}}(a)} \tag{S16}$$

High KL divergence indicates that the position has a distinctive amino acid composition compared to the background, suggesting functional specialization or strong selective constraints. Low KL divergence indicates the position follows typical amino acid frequencies, suggesting lower functional importance.

**Relation to Mutual Information:**

KL divergence is fundamentally related to mutual information. For position $i$ and target variable $T$:

$$I(i;T) = \sum_{t} P(t) D_{KL}(P(i|t) \| P(i)) \tag{S17}$$

This expresses mutual information as the expected KL divergence between the conditional distribution $P(i|t)$ and the marginal distribution $P(i)$, weighted by $P(t)$.

##### S1.5.6 Jensen-Shannon Divergence

The Jensen-Shannon (JS) Divergence [?] is a symmetrized and smoothed version of KL divergence that addresses its asymmetry and potential unboundedness. JS divergence is a true distance metric (satisfies triangle inequality after taking square root) and is always bounded.

For two probability distributions $P$ and $Q$, the JS divergence is defined as:

$$D_{JS}(P \| Q) = \frac{1}{2} D_{KL}(P \| M) + \frac{1}{2} D_{KL}(Q \| M) \tag{S18}$$

where $M = \frac{1}{2}(P+Q)$ is the average distribution. This can be generalized with weights $\pi_1, \pi_2$ such that $\pi_1 + \pi_2 = 1$:

$$D_{JS}^{\pi_1,\pi_2}(P \| Q) = \pi_1 D_{KL}(P \| M) + \pi_2 D_{KL}(Q \| M) \tag{S19}$$

where $M = \pi_1 P + \pi_2 Q$.

**Properties:**

- **Symmetric**: $D_{JS}(P \| Q) = D_{JS}(Q \| P)$
- **Bounded**: $0 \leq D_{JS}(P \| Q) \leq \log 2$ (for equal weights and base-2 logarithm)
- **Metric**: $\sqrt{D_{JS}(P \| Q)}$ is a true distance metric
- **Smooth**: Always well-defined, even when distributions have non-overlapping supports

**Application:**

We use JS divergence to compare amino acid distributions between functional subgroups or to measure the distinctiveness of a position's distribution relative to background. For comparing two sequence subsets with amino acid distributions $P_{\text{group1}}(i)$ and $P_{\text{group2}}(i)$ at position $i$:

$$D_{JS}(P_{\text{group1}}(i) \| P_{\text{group2}}(i)) = \frac{1}{2} \sum_{a} P_{\text{group1}}(i,a) \log \frac{2P_{\text{group1}}(i,a)}{P_{\text{group1}}(i,a) + P_{\text{group2}}(i,a)} + \frac{1}{2} \sum_{a} P_{\text{group2}}(i,a) \log \frac{[illegible]}{P_{\text{grou}}[illegible]} \tag{S20}$$

High JS divergence indicates that different functional variants have distinct amino acid preferences at this position, suggesting its role in determining functional specificity.

##### S1.5.7 Gini Impurity

Gini Impurity [?], originally developed in economics to measure income inequality, quantifies the degree of "impurity" or heterogeneity in a categorical distribution. In machine learning, it is widely used in decision tree algorithms (e.g., CART) for split selection.

For a discrete random variable $X$ with probability distribution $p(x)$, the Gini impurity is:

$$\text{Gini}(X) = 1 - \sum_x p(x)^2 = \sum_x p(x)(1 - p(x)) \tag{S21}$$

**Interpretation:**

Gini impurity represents the probability that two randomly selected items from the distribution belong to different categories. It ranges from 0 (perfect purity, single category) to $1 - 1/|X|$ (maximum impurity for uniform distribution over $|X|$ categories).

**Relation to Other Measures:**

Gini impurity is closely related to Rényi entropy of order 2 (collision entropy):

$$\text{Gini}(X) = 1 - \sum_x p(x)^2 = 1 - \exp(-H_2(X)) \tag{S22}$$

It is also related to Shannon entropy through the approximation:

$$H(X) \approx 2 \cdot \text{Gini}(X) \quad \text{for small Gini values} \tag{S23}$$

**Application to Protein Sequences:**

For a residue position $i$ with amino acid distribution $p_i(a)$:

$$\text{Gini}(i) = 1 - \sum_{a \in \mathcal{A}} p_i(a)^2 \tag{S24}$$

Low Gini impurity (close to 0) indicates strong conservation with one or few dominant amino acids. High Gini impurity (close to $1 - 1/20 = 0.95$ for amino acids) indicates high variability with near-uniform distribution. Gini impurity is computationally efficient and provides similar rankings to Shannon entropy for identifying conserved positions, while being less sensitive to very rare amino acids.

##### S1.5.8 Average Entropy and Ensemble Methods

Beyond individual entropic measures, we employ ensemble approaches that combine multiple measures to improve robustness and capture complementary aspects of sequence variability.

**Average Entropy:**

Average entropy aggregates entropic values across a multiple sequence alignment to provide a global measure of sequence diversity:

$$H_{\text{avg}} = \frac{1}{N} \sum_{i=1}^{N} H(i) \tag{S25}$$

where $N$ is the alignment length and $H(i)$ is any entropic measure (Shannon, Rényi, etc.) at position $i$. This provides a single scalar characterizing overall alignment conservation.

**Hard Voting:**

Hard voting implements majority consensus across multiple entropic measures. Each measure assigns a rank to each residue position based on its importance score. The final ranking is determined by majority vote or rank aggregation:

$$\text{Rank}_{\text{hard}}(i) = \text{mode}\{\text{Rank}_{\text{Shannon}}(i), \text{Rank}_{\text{Rényi}}(i), \text{Rank}_{\text{Tsallis}}(i), \ldots\} \quad \text{(S26)}$$

Alternatively, we can use Borda count, where each measure's ranking contributes equally:

$$\text{Score}_{\text{Borda}}(i) = \sum_{m \in \mathcal{M}} (N - \text{Rank}_m(i)) \quad \text{(S27)}$$

where $\mathcal{M}$ is the set of all entropic measures and $N$ is the number of positions.

**Soft Voting:**

Soft voting weights each measure's contribution according to its individual performance or confidence:

$$\text{Score}_{\text{soft}}(i) = \sum_{m \in \mathcal{M}} w_m \cdot \text{Score}_m(i) \quad \text{(S28)}$$

where $w_m$ are weights typically learned from validation data. Weights can be determined by:

- **Cross-validation performance**: Measures that better predict known key residues receive higher weights
- **Correlation with functional data**: Measures with stronger correlation to experimental measurements are upweighted
- **Inverse variance weighting**: More stable measures (lower variance across replicates) receive higher weights

For example, if we have literature-validated key residues for a protein family, we optimize weights to maximize WCS:

$$\mathbf{w}^* = \arg\max_{\mathbf{w}} \text{WCS}\left(\sum_m w_m \cdot \text{Score}_m\right) \quad \text{subject to} \quad \sum_m w_m = 1,\ w_m \geq 0 \quad \text{(S29)}$$

**Rationale for Ensemble Approaches:**

Different entropic measures emphasize different aspects of distributions:

- Shannon entropy: Balanced sensitivity to all probability levels
- Rényi/Tsallis entropy: Tunable sensitivity to rare vs. common events
- KL/JS divergence: Deviation from background or reference distributions
- Gini impurity: Computational efficiency with similar interpretability

By combining these measures through voting schemes, we achieve more robust residue importance rankings that are less sensitive to idiosyncrasies of any single measure. Empirically, we find that soft voting with performance-based weights typically outperforms individual measures and hard voting, particularly when validated against experimental mutagenesis data.

#### S1.6 Removing the Effect of Coupling

When analyzing amino acid sequences using measures such as Mutual Information (MI) or Shannon Entropy, we must account for the multidimensional dependencies inherent in sequence data. Residue positions in protein sequences are often coupled through evolutionary pressures, meaning that certain residues co-evolve or display interdependencies. As a result, conventional ranking methods based purely on MI or entropy can incorrectly highlight residues as critical due to these indirect evolutionary couplings, rather than genuine functional significance.

To address this challenge, we employ statistical techniques designed to decouple these effects and more accurately identify residues that play critical roles in protein function. We use two key measures: **Thiel's U** and the **Chi-squared ($\chi^2$) test**.

##### S1.6.1 Thiel's U

Theil's U is a statistical measure that quantifies the amount of uncertainty in one variable ($X$) that remains even after knowing another variable ($Y$). It is defined as the ratio of the conditional entropy of $X$ given $Y$ to the entropy of $X$. The formula for Theil's U is:

$$U(X \mid Y) = \frac{H(X \mid Y)}{H(X)} \tag{S30}$$

Where:

- $H(X) = -\sum_i P(x_i) \log P(x_i)$ is the entropy of $X$,
- $H(X \mid Y) = \sum_j P(y_j) H(X \mid Y = y_j)$ is the conditional entropy of $X$ given $Y$,
- $P(x_i)$ is the probability of the $i$-th category of $X$,
- $P(y_j)$ is the probability of the $j$-th category of $Y$.

The numerator, $H(X \mid Y)$, represents the uncertainty in $X$ that is removed by knowing $Y$, while the denominator, $H(X)$, represents the total uncertainty in $X$. Thus, $U(X \mid Y)$ measures the proportion of $X$'s uncertainty that can be explained by $Y$.

- If $U(X \mid Y) = 1$, then $Y$ perfectly explains $X$, and no uncertainty remains in $X$.
- If $U(X \mid Y) = 0$, then $Y$ provides no information about $X$, and the uncertainty in $X$ is unchanged. item This is an asymmetrical measure: $U(X \mid Y) \neq U(Y \mid X)$ in general.

##### S1.6.2 Chi-squared ($\chi^2$) Test

The Chi-squared ($\chi^2$) test is a statistical method for assessing the association between two categorical variables. It compares the observed frequencies of co-occurrence of amino acid residues with the expected frequencies under the assumption of independence. The formula for the Chi-squared statistic is:

$$\chi^2 = \sum \frac{(O_i - E_i)^2}{E_i} \tag{S31}$$

where: - $O_i$ represents the observed frequency of co-occurrence between residues $i$ and $j$, - $E_i$ is the expected frequency under the null hypothesis (assuming no association).

A large $\chi^2$ value indicates a significant deviation from expected frequencies, which suggests that there is a strong association between the residues. This association may represent evolutionary coupling, which needs to be accounted for when ranking residue importance.

##### S1.6.3 Decoupling Algorithm

To effectively remove the confounding effects of residue coupling, we employed an algorithm that adjusts the rankings of residue importance by considering the evolutionary coupling matrix. This decoupling process ensures that residues identified as important are not simply artifacts of evolutionary coupling. The algorithm is summarized in Algorithm **??**.

**Algorithm S1** Decoupling of Developed Measures

**Require:**

$S$ ▷ Aligned sequences

$H = \{H_i \mid i = 1, 2, \ldots, n\}$ ▷ Initial importance values for each residue

$C = \{C_{ij} \mid i, j = 1, 2, \ldots, n\}$ ▷ Coupling matrix between residues

**Ensure:**

$A = \{A_i \mid i = 1, 2, \ldots, n\}$ ▷ Adjusted importance values for each residue

Initialize $A_i \leftarrow H_i \forall i = \{1, 2, \ldots, n\}$

Initialize $L \leftarrow [1, 2, 3, \ldots, n]$ ▷ List of unprocessed residues

**while** L is not empty **do**

  $i_{\max} \leftarrow \arg\max_{i \in L} H_i$ ▷ Select residue with highest importance

  $L \leftarrow L \setminus \{i_{\max}\}$ ▷ Remove selected residue from the list

  **for** j in L **do**

    $A_j \leftarrow A_j - H_i \times C_{ij}$ ▷ Adjust entropy based on coupling matrix

  **end for**

**end while**

**return** adjusted importance values $A$

This algorithm refines the rankings by iteratively adjusting the importance values for residues, accounting for their evolutionary coupling. Residues that are highly coupled with others are adjusted down in importance, providing a more accurate ranking based on functional relevance rather than evolutionary correlation.

##### S1.6.4 Proof of Algorithm Correctness

**Theorem 1** (Decoupling Algorithm Correctness). *The decoupling algorithm produces adjusted importance values $A$ that remove indirect coupling effects while preserving direct functional relationships.*

*Proof.* Let's prove this by induction on the number of iterations of the algorithm.

**Definitions:**

- Let $H_i$ be the initial importance value of residue $i$

- Let $C_{ij}$ be the coupling coefficient between residues $i$ and $j$
- Let $A_i^k$ be the adjusted importance value of residue $i$ after $k$ iterations

**Base Case:** $k = 0$

- Initially, $A_i^0 = H_i$ for all residues $i$
- This preserves the original importance values before adjustment

**Inductive Hypothesis:** Assume after $k$ iterations, the algorithm has correctly adjusted $k$ residues by removing their coupling effects.

**Inductive Step:** For iteration $k + 1$:

1. Select residue $i_{\max}$ with highest remaining importance:

$$i_{\max} = \arg\max_{i \in L} H_i$$

2. For each remaining residue $j$, the new adjusted value is:

$$A_j^{k+1} = A_j^k - H_{i_{\max}} \times C_{i_{\max} j}$$

3. This adjustment:
   - Removes the influence of $i_{\max}$ on $j$ proportional to their coupling
   - Preserves any importance of $j$ that is independent of $i_{\max}$
   - Maintains transitivity: if $i$ influences $j$ which influences $k$, both paths are considered

**Correctness Properties:**

1. **Termination:** The algorithm terminates after $n$ iterations as $|L|$ decreases by 1 each iteration
2. **Preservation of Direct Effects:** For any residue $j$, its final adjusted value $A_j$ represents its importance minus all indirect coupling effects
3. **Order Independence:** The final adjusted values are independent of the order of processing equal-importance residues

□

##### S1.6.5 Proof of Entropy-Based Prediction Relationship

**Theorem 2** (Entropy Prediction Asymmetry)**.** *If residue A has higher entropy than residue B, then residue B cannot fully predict A, while A may potentially predict B.*

*Proof.* Let's prove this using information theory principles.

**Given:**

- $H(A)$ = entropy of residue A
- $H(B)$ = entropy of residue B

- $H(A) > H(B)$

**Part 1:** Proving B cannot fully predict A
1) By the Data Processing Inequality:

$$I(A;B) \leq \min\{H(A), H(B)\}$$

2) Since $H(A) > H(B)$:

$$I(A;B) \leq H(B) < H(A)$$

3) For perfect prediction of A by B, we would need:

$$I(A;B) = H(A)$$

4) However, from (2), we know:

$$I(A;B) < H(A)$$

Therefore, B cannot contain sufficient information to fully predict A.
**Part 2:** Showing A may predict B
1) The mutual information is symmetric:

$$I(A;B) = I(B;A)$$

2) For B to be predicted by A, we need:

$$I(A;B) = H(B)$$

3) This is possible because:

$$H(A) > H(B)$$

So A has sufficient information capacity to potentially encode all of B's information.

□

**Corollary 1.** *The entropy-based ranking in the decoupling algorithm naturally preserves the information flow direction from high-entropy residues to low-entropy residues.*

###### S1.6.6 Impact of Decoupling

Incorporating these decoupling techniques refines the identification of key residues by removing false positives caused by evolutionary coupling. By ensuring that the detected residue interactions are statistically significant, rather than being artifacts of indirect correlations, the approach improves the accuracy of predicting functionally important residues.

This methodology, implemented in tools like EVcouplings, enhances residue interaction predictions and helps deepen our understanding of protein function and evolution. By removing cross-correlations, the combined use of MI and entropy analyses becomes more robust, allowing for more precise functional annotations and guiding experimental designs with greater confidence [**?**, **?**].

#### S1.7 Quantifying Performances of Different Methods

To evaluate the alignment and consistency of rankings produced by different methods, we employed two statistical techniques: Fourier Transform-based Autocorrelation and Cross-correlation. These methods are crucial in analyzing how different rankings may be aligned or exhibit similar periodic patterns, offering a more nuanced understanding of the ranking structures generated by various methods.

**Fourier Transform-based Autocorrelation:** Fourier transforms are powerful tools for examining periodicity in sequences. By converting the ranking sequence $R = (r_1, r_2, \ldots, r_n)$ into the frequency domain, we can identify periodic components that might otherwise go unnoticed in the time domain. The autocorrelation function is computed as:

$$\text{Autocorrelation}(R, \tau) = \mathcal{F}^{-1}\left\{\mathcal{F}(R) \cdot \overline{\mathcal{F}(R)}\right\}(\tau) \tag{S32}$$

Here, $\mathcal{F}(R)$ denotes the Fourier transform of the ranking sequence $R$, and $\overline{\mathcal{F}(R)}$ is the complex conjugate. The inverse Fourier transform $\mathcal{F}^{-1}$ is used to return the autocorrelation function in the time domain. This calculation allows us to capture the repeating patterns in the rankings, indicating periodic trends or consistent shifts between ranks.

We also calculate the normalized autocorrelation:

$$\text{Normalized Autocorrelation}(R, \tau) = \frac{\text{Autocorrelation}(R, \tau)}{\text{Autocorrelation}(R, 0)} \tag{S33}$$

This normalization ensures that the autocorrelation at lag 0 (which indicates the overall consistency of the ranking without any shift) serves as the baseline for comparison. A low value for the autocorrelation at any non-zero lag indicates that the ranking does not exhibit significant periodic behavior.

Furthermore, we compute the average autocorrelation over all lags, which provides an overall measure of the ranking's cyclic patterns and periodicity:

$$\text{Average Autocorrelation}(R) = \frac{1}{n}\sum_{\tau=1}^{n} \text{Normalized Autocorrelation}(R, \tau) \tag{S34}$$

In this context, an average autocorrelation close to 1 suggests that the ranking is close to white noise, indicating a lack of significant patterns or dependencies between consecutive rankings. Lower values would indicate detectable periodicity or structure in the ranking.

**Cross-correlation:** To compare two ranking sequences $R_1$ and $R_2$, cross-correlation measures how similar the two sequences are across all positions. It is calculated as:

$$\text{CrossCorrelation}(R_1, R_2) = \frac{\sum_{i=1}^{n}(r_{1,i} - \bar{r}_1)(r_{2,i} - \bar{r}_2)}{\sqrt{\sum_{i=1}^{n}(r_{1,i} - \bar{r}_1)^2 \sum_{i=1}^{n}(r_{2,i} - \bar{r}_2)^2}} \tag{S35}$$

Where $\bar{r}_1$ and $\bar{r}_2$ are the means of the ranking sequences. The cross-correlation value ranges from -1 to 1, with 1 indicating perfect similarity between the two rankings and -1 indicating perfect inverse similarity. This metric is essential for comparing how consistently two different methods identify key residues, shedding light on their relative alignment.

#### S1.8 Assessing Performance

extbfLiterature-based Assessment: Wasserstein Concentration Score (WCS)

To rigorously assess how well computational methods identify literature-validated key residues, we employ the **Wasserstein Concentration Score (WCS)** as the sole metric for literature-based evaluation. WCS integrates both the concentration of importance on literature-identified residues and the optimality of their ranking, providing a unified, interpretable score between 0 and 1.

extbfDefinition:

Let $\mathcal{L}$ denote the set of residues identified as key in the literature, $\mathcal{R}$ the set of all residues, and $\hat{I}_i$ the normalized importance of residue $i$. The WCS is defined as:

$$\text{WCS} = \left( \frac{\sum_{i \in \mathcal{L}} \hat{I}_i}{\sum_{i \in \mathcal{R}} \hat{I}_i} \right) \cdot \left( 1 - \frac{d_{\text{swap}}}{n(L-n)} \right) \tag{S36}$$

where:

- $\hat{I}_i$ is the normalized importance of residue $i$,
- $\mathcal{L}$ is the set of literature-identified key residues,
- $\mathcal{R}$ is the set of all residues in the protein,
- $d_{\text{swap}}$ is the swap distance between the model and literature rankings (see below),
- $n = |\mathcal{L}|$ is the number of literature-identified key residues,
- $L = |\mathcal{R}|$ is the total number of residues.

extbfSwap Distance $d_{\text{swap}}$:

The swap distance $d_{\text{swap}}$ quantifies the minimum number of adjacent swaps required to transform the model's ranking of key residues into the literature ranking. This term penalizes misordering, ensuring that both the presence and correct prioritization of key residues are rewarded. The maximum possible swap distance is $n(L-n)$, which occurs when the rankings are completely reversed. Thus, the second term in the WCS formula is always between 0 and 1.

extbfInterpretation:

- The first term measures the concentration of importance on literature-validated residues (maximized when all importance is assigned to these residues).
- The second term measures the optimality of their ranking (maximized when the model's ranking matches the literature exactly).
- WCS = 1 indicates perfect agreement: all importance is concentrated on literature-validated residues, and their ranking matches the literature.
- WCS = 0 indicates no agreement: no importance is assigned to literature residues, or their ranking is maximally misordered.

This unified metric provides a rigorous, interpretable, and normalized assessment of how well computational predictions align with established biological knowledge, superseding previous coherence-based approaches.

**Machine Learning-based Performance:** This metric evaluates the predictive capability of a model using a machine learning framework, specifically Bayesian Ridge Regression. The model leverages labeled data to predict key residues, utilizing a one-hot encoded representation of protein sequences. The performance score is computed over subsets of residues as follows:

$$\text{Performance} = \frac{\sum_{i=1}^{N}\left((R_i^2 - R_0^2)\cdot\frac{(n-i)}{R_{\max}^2 - R_0^2}\right)^2}{\sum_{i=1}^{N}(n-i)^2} \tag{S37}$$

Where:

- $R_i^2$ is the $R^2$ score for the model using the top $i$-th ranked residues.
- $R_0^2$ is the baseline $R^2$ score when no residues are included (i.e., predicting using just the intercept).
- $R_{\max}^2$ is the theoretical maximum $R^2$ score the model could achieve.
- $n$ is the total number of residues.

**Theoretical Basis:** The idea behind this metric is that an ideal ranking of residues should maximize information gain early in the sequence, with each subsequent residue adding progressively less new information. In a perfect scenario:

- The $R^2$ score should increase rapidly with the inclusion of the top-ranked residues and then plateau, indicating that the most critical residues have been captured early.
- The ideal curve would show a steep initial increase (high $R^2$) after including the very first residue, followed by a flat line (plateau) as additional residues contribute negligible new information.
- Thus, a model that reaches this plateau quickly (after only a few residues) would be considered superior, as it implies a more accurate ranking of residue importance.

**Explanation of the Metric:** The performance score is designed to reflect this ideal behavior:

- **Gradient Analysis:** The term $(R_i^2 - R_0^2)$ measures the incremental predictive power gained by including the top $i$-th residue. For a perfect ranking, the difference $(R_i^2 - R_0^2)$ should be large for early residues and diminish as $i$ increases.
- **Weighted Contribution:** The weight $\frac{(n-i)}{R_{\max}^2 - R_0^2}$ assigns higher significance to early-ranked residues, aligning with the expectation that early residues are more critical for accurate prediction. This weight decreases linearly, giving diminishing importance to later residues.

- **Performance Plateau:** The squared term $(\cdot)^2$ emphasizes the significance of residues that deviate from the ideal ranking behavior. A model that reaches the performance plateau earlier will have a lower cumulative score, indicating higher efficacy.
- **Normalization:** The denominator $\sum_{i=1}^{N}(n-i)^2$ ensures the performance metric remains normalized, allowing comparisons across different models and datasets. It accounts for the total possible contribution, thus scaling the metric to a consistent range.

In summary, this performance metric quantifies the efficiency with which a model ranks residues in order of their predictive importance. An effective model should achieve a high $R^2$ score rapidly, plateauing after identifying the most informative residues, thus demonstrating both accuracy and efficiency in residue ranking.

#### S1.9 Identifying Interface Residues

To identify interface residues between antibodies and antigens, we employed a distance-based approach coupled with solvent accessible surface area (SASA) analysis. Residues from the antibody and antigen were considered part of the interface if any of their atoms were within a threshold distance ($d \leq 5.0$ Å) of atoms from the binding partner. For each interface residue, we recorded the minimum distances to interacting residues and the corresponding spatial vectors.

Interface residues were defined as:

$$I_{ab-ag} = \{r_{ab} \in Ab \mid \exists r_{ag} \in Ag, \exists a_{ab} \in r_{ab}, \exists a_{ag} \in r_{ag} : d(a_{ab}, a_{ag}) \leq d_{\text{threshold}}\} \quad \text{(S38)}$$

where $r_{ab}$ and $r_{ag}$ represent residues from antibody and antigen respectively, $a_{ab}$ and $a_{ag}$ represent atoms within these residues, and $d(a_{ab}, a_{ag})$ is the Euclidean distance between atoms.

For increased specificity, we filtered interface residues based on SASA, retaining only those with significant exposure to solvent (SASA $\geq 10.0$ Å$^2$), calculated using the Shrake-Rupley algorithm. This approach efficiently eliminates buried residues unlikely to participate in binding interactions.

#### S1.10 Energy Contribution Estimation with Conformational Flexibility

For each antibody interface residue, we estimated the maximum binding energy contribution by optimizing over both amino acid identity and small conformational adjustments. Our approach incorporates local flexibility by allowing residues to adopt optimal positions within a biologically reasonable radius ($\delta_{\max} = 4.0$ Å), simulating the induced-fit model of protein-protein interactions.

The binding energy for each residue pair was calculated as:

$$\Delta G_{\text{bind}}(r_{ab}, r_{ag}, \vec{\delta}) = \sum_{i \in \mathcal{I}} E_i \cdot f_i(d_{ab-ag} + \|\vec{\delta}\|, \vec{v}_{ab-ag} + \vec{\delta}) \quad \text{(S39)}$$

where $\mathcal{I} = \{\text{hbond}, \text{salt}, \text{hydrophobic}, \text{vdw}\}$ represents the set of interaction types, $E_i$ represents the estimated energy contribution of interaction type $i$, $f_i$ is a distance-dependent decay function that accounts for the specific properties of each interaction type, $d_{ab-ag}$ is the original distance between residues, $\vec{v}_{ab-ag}$ is the original contact vector, and $\vec{\delta}$ is the displacement vector representing conformational adjustment.

The optimal residue identity and displacement were determined through constrained optimization:

$$(\mathrm{aa}^*_{\mathrm{opt}}, \vec{\delta}^*_{\mathrm{opt}}) = \arg \min_{\mathrm{aa}\in\mathcal{A}, \|\vec{\delta}\|\leq\delta_{\max}} \Delta G_{\mathrm{bind}}(r_{ab}, r_{ag}, \vec{\delta}) \tag{S40}$$

where $\mathcal{A}$ is the set of all 20 standard amino acids. The optimization was performed using the SLSQP (Sequential Least Squares Programming) algorithm, which efficiently handles the nonlinear constraint $\|\vec{\delta}\| \leq \delta_{\max}$.

#### S1.11 Energy Function Components

The energy function incorporates four primary interaction types with their respective distance-dependent behavior:

##### S1.11.1 Hydrogen Bonds

For polar residues (Ser, Thr, Asn, Gln) and charged residues (Arg, Lys, Asp, Glu, His), hydrogen bond energy was modeled with both distance and angular dependencies:

$$E_{\mathrm{hbond}}(d, \theta) = E^0_{\mathrm{hbond}} \cdot e^{-\alpha_h \frac{d-d_{\min}}{d_{\max}-d_{\min}}} \cdot e^{-\frac{\theta^2}{2\sigma_\theta^2}} \tag{S41}$$

where $E^0_{\mathrm{hbond}} = -4.0$ kcal/mol, $d_{\min} = 1.5$ Å, $d_{\max} = 3.5$ Å, $\alpha_h = 3.0$, $\theta$ is the angle deviation from ideal hydrogen bond geometry, and $\sigma_\theta = 30°$. The angular term accounts for the directional nature of hydrogen bonds, which significantly influences their strength.

##### S1.11.2 Salt Bridges

For charged residue pairs, salt bridge energy was modeled with exponential distance decay:

$$E_{\mathrm{salt}}(d) = E^0_{\mathrm{salt}} \cdot e^{-\alpha_s \frac{d-d_{\min}}{d_{\max}-d_{\min}}} \tag{S42}$$

where $E^0_{\mathrm{salt}} = -3.0$ kcal/mol, $d_{\min} = 2.0$ Å, $d_{\max} = 5.5$ Å, and $\alpha_s = 3.0$. Salt bridges contribute significantly to binding specificity through electrostatic interactions between oppositely charged groups.

##### S1.11.3 Hydrophobic Interactions

For hydrophobic residues (Ala, Val, Ile, Leu, Met, Phe, Trp, Pro):

$$E_{\mathrm{hydrophobic}}(d) = E^0_{\mathrm{hydrophobic}} \cdot e^{-\alpha_{hp} \frac{d-d_{\min}}{d_{\max}-d_{\min}}} \tag{S43}$$

where $E^0_{\mathrm{hydrophobic}} = -1.0$ kcal/mol, $d_{\min} = 2.5$ Å, $d_{\max} = 5.0$ Å, and $\alpha_{hp} = 3.0$. These interactions are entropic in nature, driven by the exclusion of water molecules from non-polar surfaces.

##### S1.11.4 Van der Waals Interactions

To capture both attractive and repulsive components of van der Waals interactions, we implemented a Lennard-Jones potential:

$$E_{\text{vdw}}(d) = E_{\text{vdw}}^0 \cdot 4\epsilon \left[ \left(\frac{\sigma}{d}\right)^{12} - \left(\frac{\sigma}{d}\right)^{6} \right] \tag{S44}$$

where $E_{\text{vdw}}^0 = -0.5$ kcal/mol, $\epsilon = 0.2$ kcal/mol, and $\sigma = 3.5$ Å. The $r^{-12}$ term models short-range repulsion due to electron cloud overlap, while the $r^{-6}$ term represents attractive dispersion forces.

#### S1.12 Statistical Validation

To validate our approach, we performed comprehensive statistical analyses comparing predicted binding energies with experimental data and assessing the significance of our findings through multiple complementary methods:

##### S1.12.1 Interface Enrichment Analysis

We calculated the ratio of energy contributions at interface residues to total energy contributions to evaluate spatial localization of binding hotspots:

$$R_{\text{interface}} = \frac{\sum_{i \in I_{\text{interface}}} E_i}{\sum_{j \in I_{\text{all}}} E_j} \tag{S45}$$

This ratio quantifies the concentration of binding energy at the interface, with higher values indicating more focused interactions.

##### S1.12.2 Nonlinear Correlation Testing

Using XGBoost regression with Box-Cox transformation to handle non-Gaussian distributions, we assessed the relationship between our predicted energies and experimental binding data. For a given binding energy dataset $Y$ with $n$ observations, we applied the Box-Cox transformation:

$$Y^{(\lambda)} = \begin{cases} \frac{Y^\lambda - 1}{\lambda} & \text{if } \lambda \neq 0 \\ \ln(Y) & \text{if } \lambda = 0 \end{cases} \tag{S46}$$

where $\lambda$ is the transformation parameter optimized to maximize the log-likelihood function.

Statistical significance was determined through permutation testing ($n = 100$ permutations) to calculate $p$-values, and model quality was evaluated using $R^2$ scores and Akaike Information Criterion (AIC):

$$\text{AIC} = -2\ln(L) + 2k + \frac{2k(k+1)}{n-k-1} \tag{S47}$$

where $L$ is the likelihood function adjusted for the Box-Cox transformation, $k$ is the effective number of parameters, and the last term represents a small-sample correction. For the nonlinear XGBoost model, we estimated $k$ as the product of the number of trees and their maximum depth.

### S2 Mathematical Proofs

#### S2.1 Proof: Decoupling Reduces Spurious Correlations

**Theorem 3.** *Let $I(X;Y)$ denote the mutual information between residue position $X$ and target property $Y$, and let $C_{XZ}$ represent the coupling between positions $X$ and $Z$. The decoupled importance score $I'(X;Y)$ obtained by removing coupling effects satisfies:*

$$I'(X;Y) = I(X;Y) - \sum_{Z \neq X} \beta_{XZ} \cdot C_{XZ}$$

*where $\beta_{XZ}$ represents the coupling strength. This decoupling reduces spurious correlations while preserving true functional signals.*

*Proof.* Consider the information decomposition framework where the observed mutual information $I(X;Y)$ can be partitioned into:

$$I(X;Y) = I(X;Y|Z = \emptyset) + \sum_{Z \neq X} I(X;Y|Z) \quad \text{(S48)}$$

$$= I_{\text{direct}}(X;Y) + I_{\text{indirect}}(X;Y) \quad \text{(S49)}$$

The indirect component $I_{\text{indirect}}(X;Y)$ represents information about $Y$ that $X$ shares through coupling with other positions $Z$. By definition:

$$I_{\text{indirect}}(X;Y) = \sum_{Z \neq X} \beta_{XZ} \cdot C_{XZ} \cdot I(Z;Y)$$

The decoupling algorithm removes this indirect component by adjusting rankings based on the coupling matrix, yielding:

$$I'(X;Y) = I(X;Y) - I_{\text{indirect}}(X;Y) = I_{\text{direct}}(X;Y)$$

This operation preserves the direct functional relationship between $X$ and $Y$ while eliminating spurious correlations mediated by coupled positions. □

#### S2.2 Proof: WCS Metric Properties

**Theorem 4.** *The Wasserstein Concentration Score (WCS) satisfies the following properties:*

1. ***Boundedness**: $0 \leq WCS \leq 1$*
2. ***Monotonicity**: Higher concentration in top-ranked residues yields higher WCS*
3. ***Ideal Maximum**: WCS = 1 if and only if all importance is concentrated in known key residues*

*Proof.* Let $\mathbf{s} = (s_1, s_2, \ldots, s_N)$ be the normalized importance scores sorted in descending order, where $\sum_{i=1}^{N} s_i = 1$ and $s_i \geq 0$.

**Property 1 (Boundedness):** The concentration term is defined as:

$$C_k = \sum_{i=1}^{k} s_i$$

Since $s_i \geq 0$ and $\sum_{i=1}^{N} s_i = 1$, we have $0 \leq C_k \leq 1$.

The Wasserstein distance $W(\mathbf{s}_{\text{norm}}, \mathbf{p}_{\text{pulse}})$ is bounded by the maximum displacement:

$$0 \leq W(\mathbf{s}_{\text{norm}}, \mathbf{p}_{\text{pulse}}) \leq 1$$

Therefore:

$$\text{WCS} = C_k \times (1 - W(\mathbf{s}_{\text{norm}}, \mathbf{p}_{\text{pulse}})) \in [0, 1]$$

**Property 2 (Monotonicity):** Consider two score distributions $\mathbf{s}^{(1)}$ and $\mathbf{s}^{(2)}$ where $\mathbf{s}^{(2)}$ has higher concentration in the top $k$ positions. Then:

$$C_k^{(2)} \geq C_k^{(1)} \quad \text{and} \quad W^{(2)} \leq W^{(1)}$$

This implies:

$$\text{WCS}^{(2)} = C_k^{(2)} \times (1 - W^{(2)}) \geq C_k^{(1)} \times (1 - W^{(1)}) = \text{WCS}^{(1)}$$

**Property 3 (Ideal Maximum):** The ideal distribution is $\mathbf{p}_{\text{pulse}} = (1/k, 1/k, \ldots, 1/k, 0, \ldots, 0)$ with $k$ non-zero entries.

If $\mathbf{s} = \mathbf{p}_{\text{pulse}}$, then:

- $C_k = k \times (1/k) = 1$
- $W(\mathbf{s}_{\text{norm}}, \mathbf{p}_{\text{pulse}}) = 0$ (distributions are identical)
- $\text{WCS} = 1 \times (1 - 0) = 1$

Conversely, if WCS $= 1$, then both $C_k = 1$ and $W = 0$, which requires $\mathbf{s}_{\text{norm}} = \mathbf{p}_{\text{pulse}}$ and all importance concentrated in the top $k$ positions. □

#### S2.3 Computational Complexity Analysis

**Theorem 5.** *The computational complexity of the complete pipeline is $O(N^2M + N^3)$ where $N$ is the sequence length and $M$ is the number of sequences in the alignment.*

*Proof.* The algorithm consists of the following steps:

**Step 1: Entropy and MI Calculation**

- Computing entropy for each position: $O(NM)$
- Computing pairwise MI for all position pairs: $O(N^2M)$

Total for Step 1: $O(N^2M)$

**Step 2: Coupling Matrix Computation**

- Computing Cramér's V or Theil's U for all pairs: $O(N^2M)$
- Building the coupling matrix: $O(N^2)$

Total for Step 2: $O(N^2M)$

**Step 3: Decoupling Algorithm**

- Sorting scores: $O(N \log N)$
- Iterative adjustment using coupling matrix: $O(N^2)$ per iteration

- Convergence typically requires $O(N)$ iterations: $O(N^3)$

Total for Step 3: $O(N^3)$

**Step 4: WCS Calculation**

- Normalizing scores: $O(N)$
- Computing Wasserstein distance: $O(k \log k)$ where $k \ll N$

Total for Step 4: $O(N)$

**Overall Complexity:** The dominant terms are $O(N^2M)$ from MI calculation and $O(N^3)$ from decoupling. In practice:

- For typical protein families: $N \approx 100 - 500$ residues, $M \approx 1000 - 10000$ sequences
- The $O(N^2M)$ term dominates when $M > N$
- The algorithm scales linearly with the number of sequences when $N$ is fixed

Therefore, the overall complexity is $O(N^2M + N^3)$. □

### S3 Nanobody-Antigen Docking Protocol

#### S3.1 Structural Preparation

Nanobody and antigen structures were prepared for docking using standard molecular dynamics protocols. PDB structures were protonated at physiological pH (7.4) using REDUCE reduce2009 to ensure correct histidine protonation states and optimal hydrogen bonding geometry. Side chains with ambiguous rotamers were assigned using the Dunbrack rotamer library at 90% confidence threshold. All structures were energy-minimized in vacuo (1000 steps, steepest descent followed by conjugate gradient) to relieve steric clashes while preserving the experimental structure as much as possible.

For heteromeric complexes (when initial PDB contained both nanobody and antigen), chains were separated by creating independent structure files from the PDB coordinates. For separately obtained structures, chain identities were verified by sequence alignment to ensure correct molecular entity was selected.

#### S3.2 Clus2Pro Docking Configuration

Protein-protein docking was performed using Clus2Pro, an information-driven docking server that integrates experimental data (NMR chemical shift perturbations, mutational constraints, literature-curated interface residues) with physics-based docking. The following standard configuration was applied to all nanobody-antigen pairs:

##### S3.2.1 Docking Space Definition

- **Binding pocket**: Predicted from antigen surface properties (electrostatic potential, hydrophobic patches) and sequence conservation. For antibody targets, the linear epitope was identified from sequence annotation or predicted using PISA (Proteins, Interfaces, Structures and Assemblies) interface prediction on homologous complexes.

- **Search radius**: 10 Å expansion around predicted epitope center to accommodate nanobody CDR loop positioning flexibility.
- **Sampling grid**: 1.2 Å spacing, allowing adequate conformational sampling while maintaining computational efficiency.

###### S3.2.2 Full Docking Settings

| Parameter | Setting |
|---|---|
| Algorithm | Systematic rigid-body search (FFT-accelerated) |
| Scoring function | CHARMM force field + SASA burial energy |
| van der Waals radii | CHARMM36 |
| Electrostatics | Distance-dependent dielectric ($\epsilon = 4r$) |
| Hydrogen bonding | Distance-dependent, directionality-weighted |
| Desolvation penalty | Linear: 0.01 kcal/mol/Å$^2$ exposed SASA |
| Rotamer sampling | Single lowest-energy rotamer per residue |
| Interface flexibility | Fixed-backbone docking (no side-chain repacking) |
| Number of initial poses | 10,000 |
| Clustering radius | 2.5 Å RMSD |
| Top poses retained | 100 (before filtering) |

Table S2: **Clus2Pro docking parameters.** All nanobody-antigen docking calculations used this standardized configuration to ensure reproducibility and cross-complex comparability.

###### S3.2.3 Pose Selection Rules

From the 100 top-ranked poses, final representative poses were selected using a hierarchical filtering strategy:

1. **Energy threshold** (Tier 1): Retain poses with CHARMM score $< -300$ kcal/mol (typically 20–50 poses). This threshold empirically captures near-native poses while excluding severely misfolded or non-physical conformations. Poses above this threshold are rejected outright.
2. **Buried surface area criterion** (Tier 2): Among energy-filtered poses, select those with buried surface area (BSA) between 800–2000 Å$^2$. This range corresponds to typical antibody-antigen interfaces (1000 Å$^2$ median, with single-domain nanobodies often showing 800–1500 Å$^2$ due to smaller binding surface). Poses outside this range are assumed to represent spurious docking or partial binding modes.
3. **Interfacial geometry validation** (Tier 3): Inspect geometry of poses passing Tiers 1–2 for:
    - Absence of atomic clashes (minimum contact distance $\geq 2.4$ Å for any heavy atom pair)
    - Presence of at least 5 intermolecular hydrogen bonds (polar contacts)
    - Salt bridge formation (within-distance Coulombic interactions $< 0.5$ kcal/mol penalty)

Poses failing any sub-criterion are rejected.

4. **Sequence-based CDR positioning** (Tier 4): For nanobodies with known CDR loops (FR1-CDR1-FR2-CDR2-FR3-CDR3-FR4), verify that CDR3 loop (primary contact region) makes $\geq$ 50% of interface contacts and CDR1/CDR2 contribute $\geq$ 25%. This ensures immunologically realistic binding modes. Poses with atypical CDR engagement are flagged.

5. **Clustering and consensus** (Final): Poses passing all filtering tiers are clustered by interface RMSD (threshold 2.0 Å). Clusters containing $\geq$ 3 members are considered robust predictions. The lowest-energy pose within the largest cluster is selected as the representative docking pose.

###### S3.2.4 Energy Scoring Thresholds

The multi-level energy scoring used by Clus2Pro combines van der Waals, electrostatic, desolvation, and entropy terms:

$$E_{\text{total}} = E_{\text{vdW}} + E_{\text{elec}} + E_{\text{desolv}} + T \cdot S_{\text{conf}} \tag{S50}$$

Reference thresholds for pose quality assessment:

| **Energy Score (kcal/mol)** | **Quality Assessment** | **Acceptance** |
|---|---|---|
| $< -350$ | Excellent: strong binding predicted | High confidence |
| $-350$ to $-300$ | Good: favorable binding geometry | Accept |
| $-300$ to $-250$ | Marginal: borderline geometry | Conditional (inspect manually) |
| $-250$ to $-150$ | Poor: weak interactions | Reject |
| $> -150$ | Very poor: non-specific or misfolded | Reject |

Table S3: **Energy scoring thresholds for Clus2Pro nanobody docking.** Energies below $-300$ kcal/mol are considered acceptable starting points for molecular dynamics refinement. The range $-300$ to $-350$ kcal/mol corresponds to typical near-native poses found in experimental nanobody-antigen complexes.

Individual energy components were also inspected:

- **Van der Waals** ($E_{\text{vdW}}$): Should be $< -100$ kcal/mol (favorable close packing). Values $> -50$ kcal/mol suggest excessive steric strain.
- **Electrostatic** ($E_{\text{elec}}$): Typically $-50$ to $-150$ kcal/mol depending on surface charge distribution. Highly positive values ($> -10$ kcal/mol) indicate unfavorable Coulombic interactions and should prompt geometry re-inspection.
- **Desolvation** ($E_{\text{desolv}}$): Usually $-80$ to $-200$ kcal/mol (favorable burial of hydrophobic surface). Low values ($> -50$ kcal/mol) suggest minimal interface burial, characteristic of weak or non-specific binding.

#### S3.3 Molecular Dynamics Refinement

Selected docking poses were refined using AMBER20 molecular dynamics simulations (ff14SB forcefield) in explicit TIP3P solvent with periodic boundary conditions. Each complex was solvated in an octahedral box with 10 Å clearance to nearest solute atom. Counterions ($Na^+/Cl^-$) were added to achieve ionic strength of 150 mM and net neutrality.

The MD protocol consisted of:

1. Minimization: 5000 steps (2500 steepest descent + 2500 conjugate gradient)
2. NVT equilibration: 100 ps, 310 K, weak position restraints ($k_B = 10$ kcal/mol/Å$^2$) on non-hydrogen protein atoms
3. NPT equilibration: 100 ps, 310 K, 1 atm, restraints maintained
4. Production: 50 ns unrestrained MD, Langevin thermostat ($\gamma = 1$ ps$^{-1}$), Berendsen barostat

Trajectory snapshots at 1 ps intervals were saved for post-simulation analysis. Final nanobody-antigen complex structures were taken as the average structure from the last 10 ns of the MD trajectory (after equilibration).

#### S3.4 Complex Validation Metrics

The quality of final docked and refined complexes was assessed using:

- **Interface RMSD**: Calculated for interface residues (heavy atoms within 4.5 Å of partner chain) relative to initial docked pose. Stable MD trajectories show RMSD plateau $< 3$ Å.
- **Hydrogen bond network**: Identified inter-chain H-bonds (donor-acceptor distance $< 3.5$ Å, angle $> 120$) and tracked occupancy over the MD trajectory. Persistent H-bonds (occupancy $> 60\%$) are considered reliable.
- **Binding interface stability**: Per-residue interaction energy decomposition (computed using MMPBSA in the last 10 ns) to identify key nanobody and antigen residues driving binding.
- **CDR loop geometry**: Analyzed CDR1–3 backbone conformations to ensure they remain in known solution ensembles (VH/VL domain databases).

These validation metrics ensure that predicted nanobody-antigen complexes are physically reasonable and suitable for subsequent energetic and structural analysis.

### References


[1] Sumbalova, L., Stourac, J., Martinek, T., Bednar, D., & Damborsky, J. (2018). HotSpot Wizard 3.0: web server for automated design of mutations and smart libraries based on sequence input information. *Nucleic Acids Research*, **46**(W1), W356-W362. doi:10.1093/nar/gky500.

[2] Jumper, J., et al. Highly accurate protein structure prediction with AlphaFold. *Nature* **596**, 583-589 (2021). `https://www.nature.com/articles/s41586-021-03819-2`

[3] Marks, D.S., et al. "Protein 3D structure computed from evolutionary sequence variation." *PLoS One*, vol. 6, no. 12, 2011, e28766.

[4] Hopf, T.A., et al. "Mutation effects predicted from sequence co-variation." *Nature Biotechnology*, vol. 35, no. 2, 2017, pp. 128-135.

[5] Lassmann, Timo. "Kalign 3: multiple sequence alignment of large datasets." *Bioinformatics*, vol. 36, no. 6, pp. 1928–1929, 2019. `https://doi.org/10.1093/bioinformatics/btz795`.

[6] Riesselman, A.J., et al. "Deep generative models of genetic variation capture the effects of mutations," *Nature Methods*, 2018, 15(10), pp. 816-822.

[7] Shannon, C. E. "A mathematical theory of communication." *Bell System Technical Journal*, vol. 27, 1948, pp. 379-423.

[8] Cover, T. M., & Thomas, J. A. "Elements of Information Theory." *Wiley-Interscience*, 2006.

[9] Göbel, U., Sander, C., Schneider, R., & Valencia, A. "Correlated mutations and residue contacts in proteins." *Proteins: Structure, Function, and Bioinformatics*, vol. 18, no. 4, 1994, pp. 309-317.

[10] Dorsam, R. T., & Gutkind, J. S. "G-protein-coupled receptors and cancer." *Nature Reviews Cancer*, vol. 7, no. 2, 2010, pp. 79-94.

[11] Ng, P. C., & Henikoff, S. "SIFT: Predicting amino acid changes that affect protein function." *Nucleic Acids Research*, vol. 31, no. 13, 2003, pp. 3812-3814.

[12] Pupko, T., Bell, R. E., Mayrose, I., Glaser, F., & Ben-Tal, N. "Rate4Site: an algorithmic tool for the identification of functional regions in proteins by surface mapping of evolutionary determinants within their homologues." *Bioinformatics*, vol. 18, suppl. 1, 2002, pp. S71-S77.

[13] Rényi, A. "On measures of entropy and information." *Proceedings of the Fourth Berkeley Symposium on Mathematical Statistics and Probability*, vol. 1, 1961, pp. 547-561.

[14] Tsallis, C. "Possible generalization of Boltzmann-Gibbs statistics." *Journal of Statistical Physics*, vol. 52, 1988, pp. 479-487.

[15] Kullback, S., & Leibler, R. A. "On information and sufficiency." *The Annals of Mathematical Statistics*, vol. 22, no. 1, 1951, pp. 79-86.

[16] von Neumann, J. "Mathematical Foundations of Quantum Mechanics." Princeton University Press, 1955.

[17] Gini, C. "Variabilità e mutabilità." *Reprinted in Memorie di metodologica statistica*, 1912.

[18] Kolmogorov, A. N. "Three approaches to the quantitative definition of information." *Problems of Information Transmission*, vol. 1, no. 1, 1965, pp. 1-7.

[19] Lin, J. "Divergence measures based on the Shannon entropy." *IEEE Transactions on Information Theory*, vol. 37, no. 1, 1991, pp. 145-151.
